# Single-cell Map of Diverse Immune Phenotypes Driven by the Tumor Microenvironment

**DOI:** 10.1101/221994

**Authors:** Elham Azizi, Ambrose J. Carr, George Plitas, Andrew E. Cornish, Catherine Konopacki, Sandhya Prabhakaran, Juozas Nainys, Kenmin Wu, Vaidotas Kiseliovas, Manu Setty, Kristy Choi, Rachel M. Fromme, Phuong Dao, Peter T. McKenney, Ruby C. Wasti, Krishna Kadaveru, Linas Mazutis, Alexander Y. Rudensky, Dana Pe’er

**Author notes:** These authors contributed equally (Elham Azizi, Ambrose J. Carr, George Plitas, Andrew E. Cornish). Co-senior author and lead contact (Alexander Y. Rudensky, Dana Pe’er).

## Abstract

Knowledge of immune cell phenotypes in the tumor microenvironment is essential for understanding mechanisms of cancer progression and immunotherapy response. We created an immune map of breast cancer using single-cell RNA-seq data from 45,000 immune cells from eight breast carcinomas, as well as matched normal breast tissue, blood, and lymph node. We developed a preprocessing pipeline, SEQC, and a Bayesian clustering and normalization method, Biscuit, to address computational challenges inherent to single-cell data. Despite significant similarity between normal and tumor tissue-resident immune cells, we observed continuous tumor-specific phenotypic expansions driven by environmental cues. Analysis of paired single-cell RNA and T cell receptor (TCR) sequencing data from 27,000 additional T cells revealed the combinatorial impact of TCR utilization on phenotypic diversity. Our results support a model of continuous activation in T cells and do not comport with the macrophage polarization model in cancer, with important implications for characterizing tumor-infiltrating immune cells.

## INTRODUCTION

Recent evidence suggests that cells of the innate and adaptive immune system serve an essential accessory function in non-lymphoid normal tissues and in tumors (Fan and Rudensky, 2016). Naïve, effector, and memory T lymphocytes, as well as chronically stimulated dysfunctional T lymphocytes, are considered to be the principal T cell differentiation states. These discrete states are characterized by their functional features as well as patterns of co-stimulatory and co-inhibitory receptor expression. Co-stimulatory receptors, such as CD28, ICOS, OX40, CD40L, and CD137, markedly enhance TCR-dependent T cell activation, whereas increasing levels of co-inhibitory receptors CTLA-4, PD-1, TIGIT, LAG3, TIM-3, and CD160 are characteristic of progressive T cell dysfunction and loss of self-renewal potential (exhaustion). The success of cancer immunotherapy based on CTLA-4 or PD-1 blockade has been attributed to prevention or reversal of intratumoral T cell exhaustion (Pauken and Wherry, 2015). Regulatory T (Treg) cells, expressing high amounts of the transcription factor Foxp3, curtail activity of effector T cells and other immune cell types under physiologic conditions and are found in markedly increased numbers in solid organ tumors (Josefowicz et al., 2012; Tanaka and Sakaguchi, 2017). This dedicated lineage of suppressive cells is thought to play a prominent role in cancer progression, and Treg cell targeting is considered a potentially promising strategy for tumor immunotherapy.

Likewise, two principal functional states are standardly recognized in tumor-associated macrophages: pro-inflammatory M1 macrophages, which partake in protective anti-bacterial responses and are thought to oppose tumor progression, and tissue reparative M2 macrophages, which promote tumor growth and metastasis (Mantovani and Locati, 2013; Mills et al., 2000). In addition, a heterogeneous group of monocytic cells and neutrophils, summarily classified as myeloid derived suppressor cells (MDSC), are capable of production of immunosuppressive mediators and have also been suggested to support tumor progression and limit tumor immunity (Kumar et al., 2016).

Although immunotherapy treatments targeting CTLA-4 and PD-1 have been successful in treating melanoma, lung cancer, and kidney cancer (Topalian et al., 2015), meaningful clinical responses have only been observed in a subset of patients and cancer types. The observed variation in treatment efficacy has been connected to heterogeneity in the immune status and immune cell type composition of individual tumors. Similarly, in breast cancer significant heterogeneity in immune composition is observed across tumor subtypes as well as patients (Dushyanthen et al., 2015). These observations raise the question of whether immune cell states differ in normal and tumor tissue, and whether they represent a limited number of discrete differentiation or activation intermediates with defined gene expression characteristics. Alternatively, these cell states may represent a continuum of cell states occupying a single contiguous spectrum, whose features, including “phenotypic volume,” may be affected by the tumor microenvironment.

Recent studies of human immune cells in lung adenocarcinoma and clear cell renal cell carcinoma using CyTOF mass cytometry (Chevrier et al., 2017; Lavin et al., 2017) and bulk RNA-seq analysis of tumor resident immune cells (Rooney et al., 2015; Senbabaoglu et al., 2016) have provided broad characterization of the composition of main immune cell subsets. Further studies employing single-cell RNA-seq analysis have begun to explore finer definitions of immune cell subsets in tumors (Tirosh et al., 2016; Zheng et al., 2017), but their scale has been insufficient to address the major questions above.

Thus, we sought to undertake a large-scale, high-dimensional analysis of cells of hematopoietic origin in human breast tumors of various types – as well as paired normal breast tissue, peripheral blood, and a lymph node – using single-cell RNA-seq and paired TCR-seq combined with novel computational approaches. Our analyses revealed significantly increased heterogeneity of intratumoral cells of both lymphoid and myeloid cell lineages, occupying markedly expanded contiguous phenotypic space in comparison to normal breast tissue. This heterogeneity can be viewed as the result of combinatorial responses to environmental stimuli, and in the case of T cells is shaped by TCR specificity. The observed continuum of cell states likely reflects their progressive cellular activation and differentiation and argues strongly against the classical notion of few discrete states of differentiation or activation of individual cell types shaping the tumor microenvironment.

## RESULTS

### Single-cell RNA-seq of breast carcinoma resident immune cells

To generate a deep transcriptional map of the immune cell states in human breast cancer, we constructed an atlas comprising 47,016 CD45^+^ cells collected from 8 primary breast carcinomas from treatment naïve patients including estrogen receptor (ER^+^) and progesterone receptor (PR^+^) positive, human epidermal growth factor receptor 2 amplified (Her2^+^), and triple negative (TNBC) cancers. To assess the common effects of the tumor microenvironment across breast cancer subtypes on immune cell phenotypes, we also analyzed CD45^+^ cells from matched normal breast tissue, peripheral blood, and lymph node obtained from fresh surgical specimens. The corresponding FACS-purified CD45^+^ cell populations were subjected to single-cell barcoding and RNA sequencing (scRNA-seq) using the inDrop platform (Figure 1A,B, STAR Methods) (Zilionis et al., 2017). We developed a scRNA-seq pipeline, SEQC, for processing the data, providing increased sensitivity and selectivity in its resulting single-cell profiles (STAR Methods).

**Figure 1:**
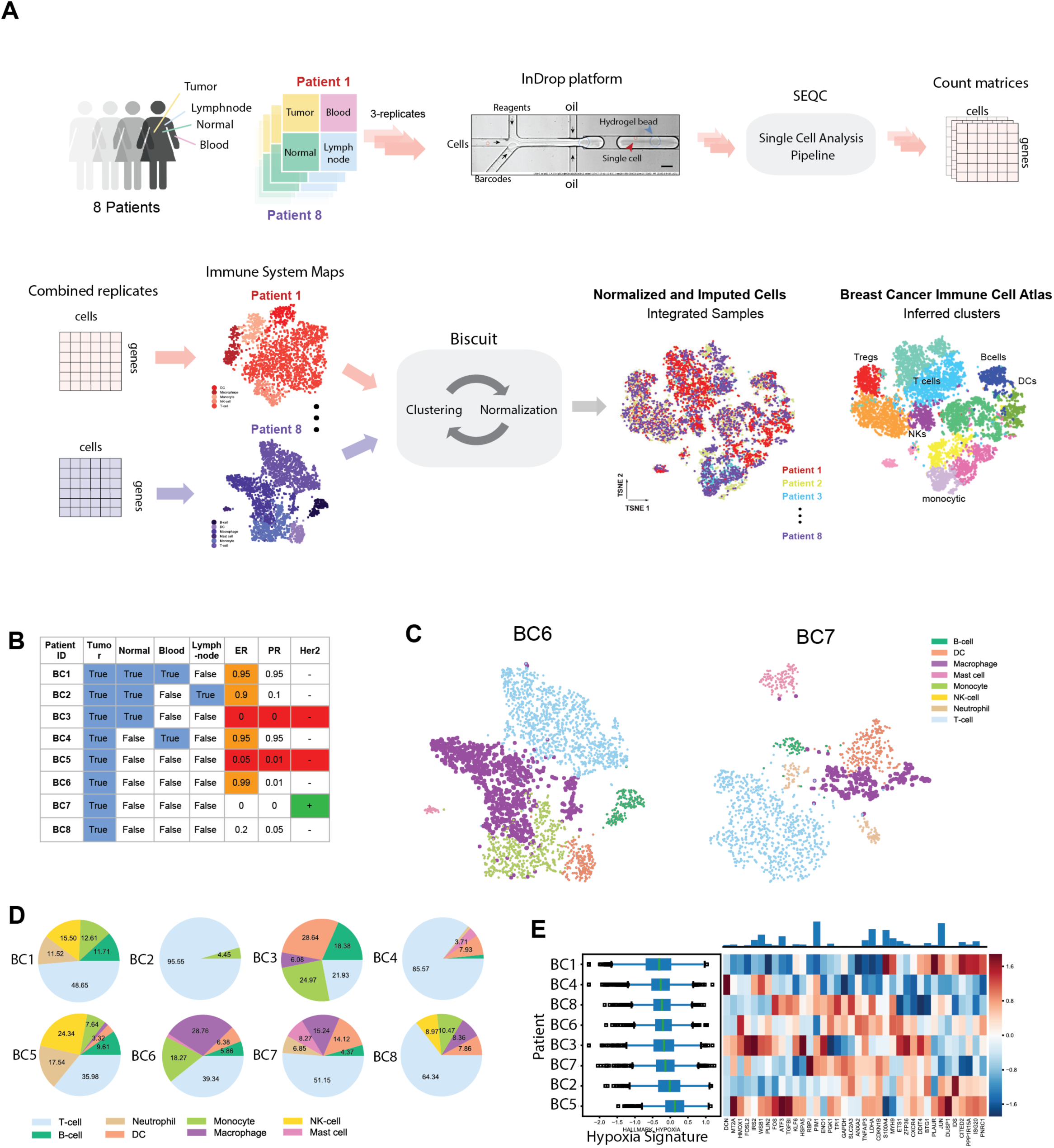
Single-Cell RNA Seq Experimental Design and Initial Data Exploration. (A) Flow chart displaying experimental design and analysis strategy. (B) Summary of samples obtained and patient metadata; more details in S1A. ER and PR values denote proportion of tumor cell nuclei that express the receptor. (C) t-SNE projection of complete immune systems from two example breast cancer tumors. scRNA-seq data for each tumor is processed with pipeline in Figure S1B and library size-normalized; each dot represents a single-cell colored by PhenoGraph clustering, and clusters are labeled by inferred cell types. Additional tumors are presented in in Figure S1D. (D) Pie charts of cell type fractions for each patient’s tumor-infiltrating immune cells, colored by cell type. (E) Left: Boxplots showing expression of Hypoxia signature (defined as the mean normalized expression of genes in the hypoxia signature listed in Table S4) across immune cells from each patient. Right: Heatmap displaying z-scored mean expression of genes in hypoxia signature. Top: Barplot showing total expression of each gene, across all patients. See Figure S1E-G for additional signatures.

We first verified that major immune cell types were identifiable in each patient using PhenoGraph clustering (Levine et al., 2015). We annotated clusters using genome-wide correlations between cluster mean expression levels and previously characterized transcriptional profiles of sorted immune cell subsets (Jeffrey et al., 2006; Novershtern et al., 2011), as well as their canonical markers (see STAR Methods). We were able to identify the majority of immune cell types expected to be present in human tumors, including monocytes, macrophages, dendritic cells (DC), T cells, B cells, mast cells, and neutrophils (Figure 1C, S1C). Thus, we were able to capture a comprehensive representation of the immune ecosystem from each individual tumor.

### Variation between individual tumor immune microenvironments

In agreement with recent studies (Chevrier et al., 2017; Lavin et al., 2017), we found a large degree of variation in the immune composition of each tumor (Figure 1D). For example, the myeloid and T cell fractions constituted 4-55% and 21%-96%, respectively. To determine the reliability of inDrop in representative sampling of heterogeneous populations, we compared the proportions of cell types as measured by flow cytometry and scRNA-seq. Although this comparison revealed a significant bias towards monocytic lineage cell subsets in scRNA-seq, we observed a high correlation between cell type frequencies across patient samples (r^2^ > 0.8, Figure S1D). The observed bias, likely due to the higher RNA yield of myeloid cells vs. T cells, was systemic, and did not adversely affect our analyses.

This genome-wide view allowed us to assess system-level differences between immune cell consortia in individual patients in, for example, metabolic signatures, including hypoxia (Figure 1E). It is notable that while all patients expressed a similar average degree of a hypoxia signature, they differed in expression at the level of individual genes in the signature. Similar variation was observed in fatty acid metabolism, glycolysis, and phosphorylation (Figure S1E-G).

### Integration of data across multiple tumors

To enable a systematic comparison across patients, we merged the data from all tumors. However, we observed that cells from the same patient were often more similar than cells of the same lineage across patients (Figure 2A). This was likely due both to batch effects and standard normalization procedures that conflate biological signal and technical differences. We also observed an increase in the number of molecules captured from activated cells, likely due to the previously observed increase in total RNA abundance upon activation (Cheadle et al., 2005; Marrack et al., 2000; Singer et al., 2017). In addition, our analyses showed an activation gradient in CD8 T cells in tumors, most pronounced in a TNBC tumor (BC3), in agreement with previous reports (Figure 2B) (Dushyanthen et al., 2015). To remove technical effects while retaining convolved biological variation, we clustered the combined data using an improved version of Biscuit, a hierarchical Bayesian model that infers clusters while simultaneously normalizing and imputing, allowing us to correct for cell- and batch-intrinsic variation (Prabhakaran et al., 2016) (STAR Methods). Using Biscuit, we normalized cells that are assigned to the same type together, and extracted interpretable parameters to characterize each cluster.

**Figure 2:**
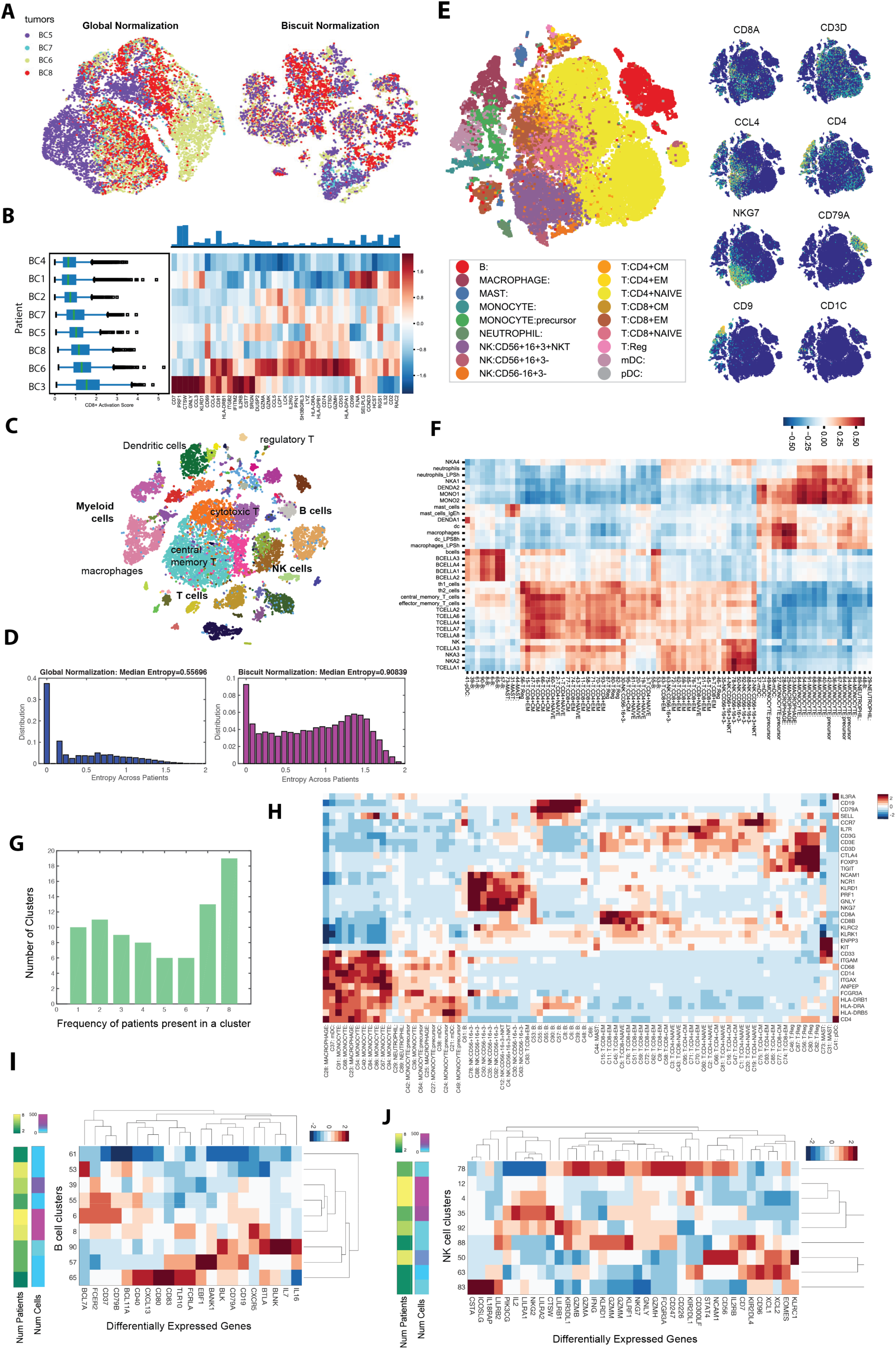
Unbiased Characterization of the Immune System Across Breast Cancer Patients. (A) t-SNE projection of immune cells from 4 breast cancer tumors after library-size normalization (left panel) and Biscuit normalization and imputation (right panel). Cells are colored by tumor (patient). Less mixing of tumors indicates either batch effects or patient-specific cell states. (B) Left: Boxplots showing expression of CD8 T cell activation signature (defined as the normalized mean expression of genes in the activation signature listed in Table S4) across immune cells from each patient, in the same format as Figure 1A. Expression of T cell activation signature shows variability across patients and increased expression in patients BC6 and BC3. (C) t-SNE map of immune cells from all 8 breast tumors after Biscuit normalization and imputation showing rich structure and diverse cell types. Cells colored by Biscuit clusters and labeled with inferred cell types. (D) Histogram depicting entropy of the patient distribution as a measure of sample mixing. Entropy is computed per cell, based on the distribution of patients in (30-NN) local cell neighborhoods after library-size normalization (left panel) as compared to Biscuit (right panel). (E) t-SNE projection of complete atlas of immune cells, post-Biscuit normalization, from all patients and tissues including tumor, blood, lymph, and contralateral normal tissue, labeled by inferred cell type (left panel) and normalized expression of 8 immune cell markers (right panel). Figure S2 presents further details on inferred clusters with complete annotations in Table S2. (F) Pearson correlations between cluster expression centroids and bulk RNA-seq data from purified immune populations from (Jeffrey et al., 2006; Novershtern et al., 2011). (G) Histogram of frequency of patients contributing to each cluster showing that 19 clusters are present in all 8 patients and 10 clusters are patient-specific. (H) Expression of canonical and cell type markers across clusters, z-score normalized across clusters. (I,J) Differentially expressed genes in B cell (I) and NK cell (J) clusters, standardized by z-scores within cell type to highlight clusters with higher or lower expression of the marker compared to the average B or NK cell cluster (refer to Table S3 for all DEGs in these and other clusters).

After applying Biscuit to the data from all tumors (Figure 2C), we found clusters covering T cell, macrophage, monocyte, B cell, and NK cell clusters. We used an entropy-based measure to quantify the mixing of samples (STAR Methods): while cells were most similar within individual samples using standard normalization, this was corrected after Biscuit normalization, with significantly improved mixing of cells across patients (U=1.7721e+09, p=0, Figure 2A,D). Using this approach, we successfully retained information on immune cell activation while stabilizing differences in library size, and uncovered a rich and robust structure in imputed data, suggesting diversity in immune cell subtypes (Figure 2A, C).

### Breast tumor immune cell atlas reveals substantial diversity of cell states

To construct a global atlas of immune cells we merged data from 47,016 cells across all tissues and patients revealing a diverse set of 83 clusters (Figure 2E, F; S2A, B). The large number of clusters prompted us to test their robustness using cross-validation (STAR Methods), showing robust cluster assignments (Figure S2C). Most clusters were shared across multiple patients, with only 10 being patient-specific (Figure 2G; STAR Methods). We used entropy as a quantitative metric and found that the clusters vary in their degree of patient mixing (Figure S2D).

To assign each cluster to a cell type, we compared cluster mean expression to sorted bulk datasets (Figure 2E, F) and identified 38 T cell, 27 myeloid lineage, 9 B cell, and 9 NK cell clusters (Table S2). These annotations were confirmed and honed using the expression of canonical markers (Figure 2H). Of the T cell clusters, we identified 15 CD8+ and 21 CD4+ clusters, which were split into 9 naive, 7 central memory, 15 effector memory, and 5 Treg clusters. The myeloid clusters divided into 3 macrophage, 3 mast cell, 4 neutrophil, 3 dendritic cell, 1 plasmacytoid dendritic cell, and 13 monocytic clusters. Finally, we identified 3 CD56^−^ NK cell clusters, and 6 CD56^+^ NK cell clusters, 2 of which are likely NKT cells (Figure 2I, J).

Broad cell types (e.g. 15 effector memory T cell clusters) are defined by a small set of surface markers (Figure 2F,H). To further differentiate these clusters, we used their genome-wide profiles. Biscuit identifies clusters based on both mean expression and covariance patterns (i.e. co-expression of genes). These parameters helped identify differentially expressed genes that characterized multiple subpopulations within major cell types (Table S2, S3). Moreover, we observed a prominent effect of covariance in defining the T cell clusters by measuring pairwise cluster similarity with and without the effect of mean expression (Figure S2E, F); large differences between most clusters remained even after mean gene expression was equalized. Thus, our approach robustly identified distinct cell states that were shared across multiple tumor microenvironments. As T and myeloid cells represented the most abundant cell subsets, and are considered the most clinically impactful, we focused subsequent in-depth analyses on these major cell types.

### Tissue environment impacts the diversity of immune phenotypic states

A key goal of this study was to quantify the extent to which variation in immune phenotypes is driven by their tissue of residence (i.e. cancerous vs. normal breast tissue), using peripheral blood or lymph node cells as references. We used t-SNE co-embedding (Amir el et al., 2013; van der Maaten and Hinton, 2008) to visualize phenotypic overlap between tissues, which showed that T cells in blood and lymph node exhibited significantly dissimilar phenotypes compared with T cells in cancerous and normal breast tissue (Figure 3A, B). While T and myeloid lineage cells exhibited considerable overlap between tumor and normal tissue samples, we observed increased phenotypic heterogeneity and expansion of these populations in the tumor (Figure 3B). Naive T cells were strongly enriched in three blood-specific clusters (*χ*^2^=361.4, df=1, p=3e-80), while B cells were more prevalent in the lymph node than in other tissues (*χ*^2^=1737.1, df=1, p=0.0) (Figure 3C). A subset of T cell clusters was present in both tumor and normal tissue, but cytotoxic T cell clusters were more abundant in tumor (*χ*^2^=93.7, df=1, p=3e-25), as were Treg clusters (*χ*^2^=336.0, df=1, p=5e-91). Moreover, some myeloid clusters were shared between normal and tumor tissue, whereas clusters of more activated macrophages were specific to tumor (tumor-associated macrophages, or TAMs) (*χ*^2^=2420.6, df=1, p=0.0). These findings highlight that tissue of residence is a significant determinant of immune phenotypes, and that biomarkers based on blood immune cells may not necessarily reflect immune cell composition in tumor.

**Figure 3:**
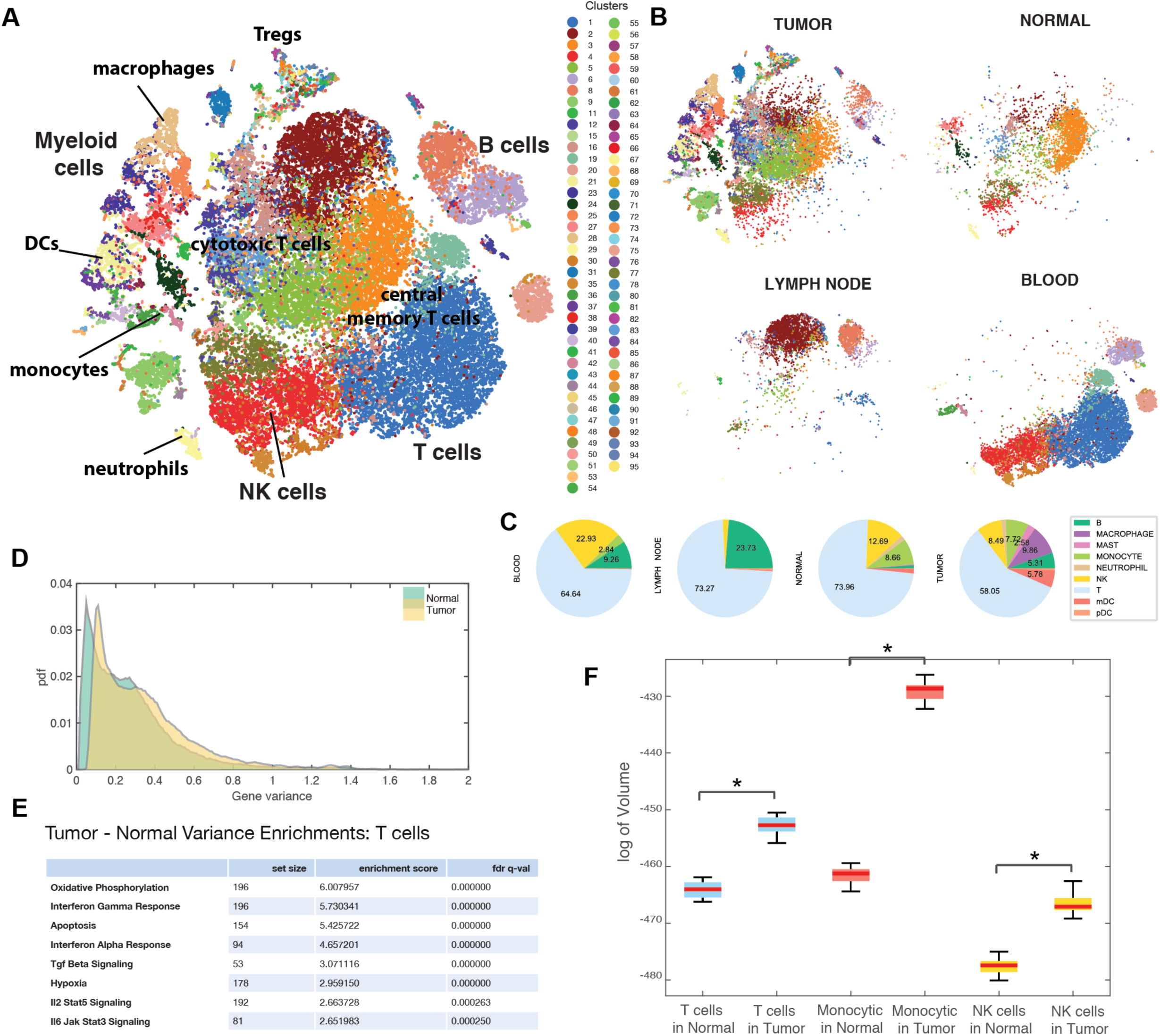
Impact of Environment on Breast Immune Cells. (A) Breast immune cell atlas constructed from combining all patient samples (BC1-8) and tissues using Biscuit and projected with t-SNE. Each dot represents a cell and is colored by cluster label; major cell types are marked according to Figure 2F,H and Table S2,3. (B) Subsets of immune atlas t-SNE projection in (A) showing cells from each tissue separately on the same coordinates as 3A to highlight the differences between tissue compartments. (C) Proportions of cell types across tissue types. (D) Distribution of variance of expression, computed for each gene across all immune cells (all patients), from tumor tissue compared to that in normal breast tissue. (E) Hallmark GSEA enrichment results on genes with highest difference in variance in tumor T cells vs normal tissue T cells. See Figure S3 for enrichment in monocytic and NK cells. Most significant results are shown; full lists of enrichments are presented in Table S5. (F) Phenotypic volume in log-scale (defined as determinant of gene expression covariance matrix, detailed in STAR) of immune cell types in tumor compared to normal breast tissue, controlled for number of cells. Massive expansion of volume spanned by independent phenotypes in tumor compared to normal tissue is shown for all three major cell types.

### Tumor microenvironment drives an expansion of immune cell phenotypes

We observed a large number of normal breast tissue resident immune cell states, including 13 myeloid and 19 T cell clusters that were not observed in circulation or in the secondary lymphoid tissue. Furthermore, the set of clusters found in normal breast tissue cells represented a subset of those observed in the tumors; 14 myeloid and 17 T cell clusters were only found in tumor, doubling the number of observed clusters relative to normal tissue. In distinction, there were no clusters specific to normal tissue.

This increased diversity of cell states was driven by a significant increase in the variance of gene expression in tumor compared to normal tissue (Figure 3D). We found that the genes with the largest increase in variance were enriched in signaling pathways activated in the tumor environment, including Type I (IFNα) and II interferons (IFNγ), TNFα, TGFβ, IL6/JAK/STAT signaling, and hypoxia (STAR Methods) (Figure 3E; S3A, B; Table S5).

To further explore this increase in variance, we defined a metric for the “phenotypic volume” occupied by cells. Specifically, this metric uses the covariance in gene expression to measure the volume spanned by independent phenotypes, which grows with the number of active phenotypes as well as the degree of independence between them (see STAR methods). Using this metric, we compared the phenotypic volume occupied by each cell type in normal vs. tumor tissue. Assessment of the change in volume showed a significant increase in the phenotypic volume of all major cell types, including T cells, myeloid cells, and NK cells (U-test, p = 0) in the tumor compared to normal mammary gland tissue (Figure 3F). Precisely, the fold change in volume was 7.39e4 in T cells, 1.18e14 in myeloid cells, and 6.08e4 in NK cells, indicating a massive increase in phenotypic volume in tumor compared to normal tissue. These data suggest that increased heterogeneity of cell states and marked phenotypic expansions found within the tumor were likely due to more diverse local microenvironments within the tumor, which differ in the extent of inflammation, hypoxia, expression of ligands for activating and inhibitory receptors, and nutrient supply (Finger and Giaccia, 2010; Jimenez-Sanchez et al., 2017).

### Intratumoral T cells reside on continuous components of variation

We used diffusion maps (Coifman et al., 2005; Haghverdi et al., 2015) to characterize the most significant sources of the observed phenotypic variation. This analysis was done separately on T and myeloid cells to avoid biases from cell type-specific capture rates (Figure S1D). For T cell clusters, while some components distinguished discrete clusters, the majority of components defined gradual trends of variation (Figure 4A, S4A). The top 3 informative components correlate, respectively, with signatures for activation, terminal differentiation, and hypoxia (Figure 4B-D; Figure S4B-F; STAR Methods).

**Figure 4:**
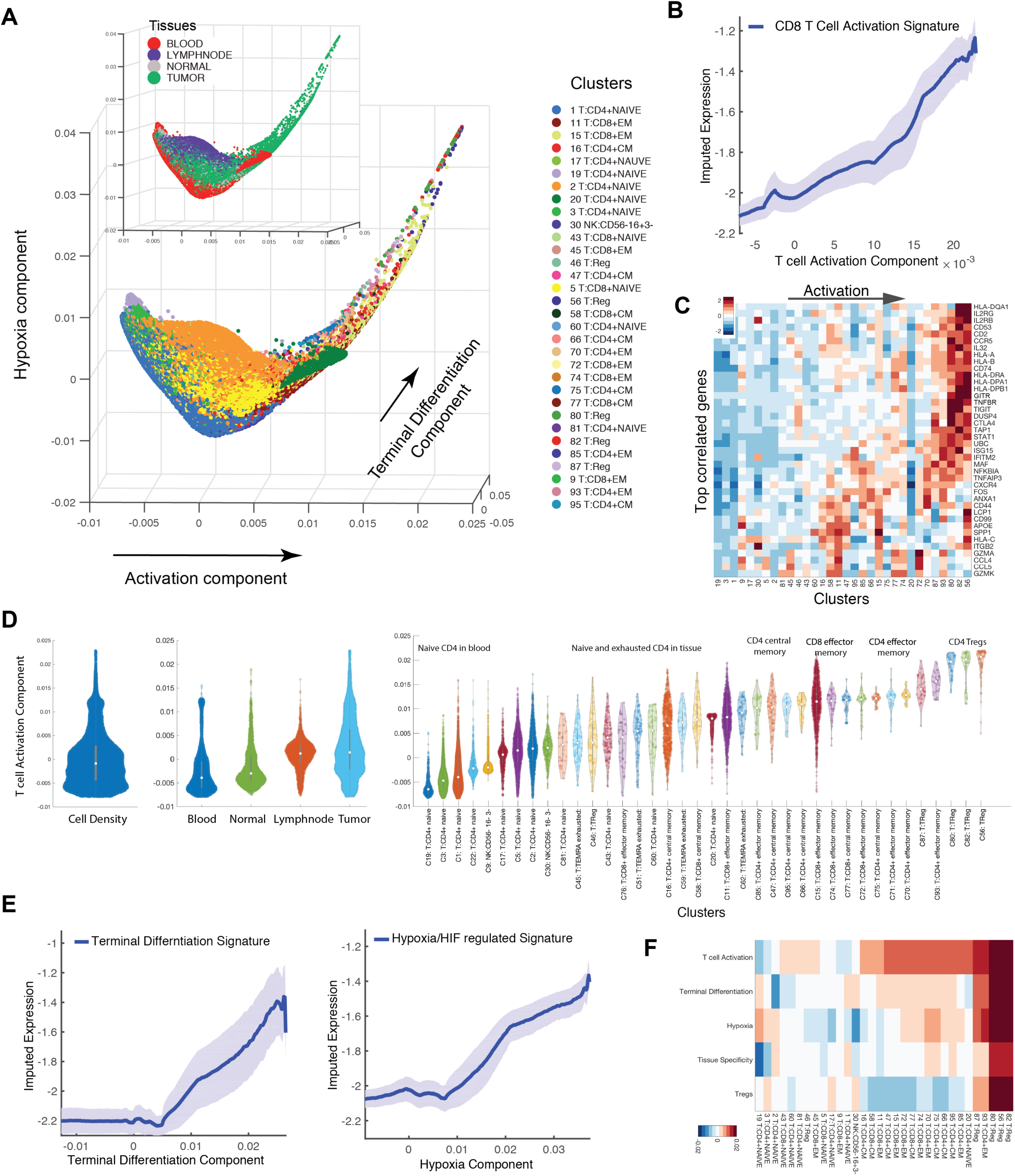
Detailed Characterization of T Cells. (A) Visualization of all cells from T Cell clusters using first, second, and third diffusion components (two uninformative components denoting isolated NKT and blood-specific clusters were removed). Each dot represents a cell colored by cluster, and by tissue type in insert. The main trajectories are indicated with arrows and annotated with the signature most correlated with each component. See Figure S4D for additional components. (B) Traceplot of CD8 T cell activation signature (defined as mean expression across genes in signature listed in Table S4) for all T cells along first diffusion component. Cells are sorted based on their projection along the component (x-axis), and the blue line indicates moving average over gene expression, using a sliding window of length equal to 5% of total number of T cells; shaded area displays standard error (y-axis). (C) Heatmap showing expression of immune-related genes with the largest positive correlations with activation component, averaged per cluster and z-score standardized across clusters; columns (clusters) are ordered by mean projection along the component. See Figure S4 for heatmaps for additional components. (D) Violin plot showing the projection of T-cells along activation component aggregated by total density (left), tissue type (middle), and cluster (right). See Figure S4 for violin plots for additional components. Number of dots inside each violin are proportional to number of cells. (E) Trace-plots (as in B) of (left) terminal differentiation signature along second component and (right) hypoxia signature along third component. List of genes associated with signatures are presented in Table S4. (F) Heatmap of cells projected on each diffusion component (rows) averaged by cluster (columns).

The most informative component of variation, labeled activation, is highly correlated with gene signatures of T cell activation and progressive differentiation, along with IFNγ signaling (p=0.0) (STAR Methods). The mean expression of the activation signature steadily increases along the component (Figure 4B), with a concomitant gradual increase in expression of specific activation-related genes (Figure 4C). Intratumoral T cell populations, including Treg and effector memory T cells, are enriched at the activated end of the component (t-test p=0.0, Figure 4A, D), while naïve peripheral blood T cells congregate at the least activated terminus, consistent with their quiescent state (t-test p=0.0, Figure 4D). Further, though the mean expression levels of clusters vary gradually along the component, there is a wide range of activation states within each cluster (Figure 4D). Examining genes most correlated with the component reveals genes known to increase upon activation and progressive differentiation, including cytolytic effector molecules granzymes A and K (GZMA and GZMK), pro-inflammatory cytokines (IL-32), cytokine receptor subunits (IL2RB), chemokines (CCL4, CCL5), and their receptors (CXCR4, CCR5) (Figure 4C).

The next most informative component of variation was labeled terminal differentiation (Figure 4E); the genes most correlated with it include co-stimulatory molecules (CD2, GITR, OX40, and 4-1BB) as well as co-inhibitory receptors (CTLA-4 and TIGIT) (Figure S4B). This set also included FOXP3, IL2RA, and ENTPD1 (CD39), genes characteristic of Tregs (Josefowicz et al., 2012). There is a moderate degree of overlap in the genes most correlated with the activation and terminal differentiation components, consistent with previous single-cell studies (Tirosh et al., 2016) (Figure 4A, C; S4B). However, there are also important exceptions, including the markers of exhaustion listed above, and the order of clusters differs along the two components (Figure 4F). Indeed, some clusters—notably lymph node T cells (e.g. cluster 16)—express higher levels of activation than terminal differentiation (t-test p=0.0; Figure S4B-D), consistent with T cell exhaustion and terminal differentiation being prominent in non-lymphoid tissues.

Interestingly, visualizing the T cell activation and terminal differentiation components together revealed remarkable continuity, in essence representing a single continuous trajectory of T cells towards a terminal state (Figure 4A, S4D). Thus, T cells reside along a broad continuum of activation, suggesting that their conventional classification into relatively few discrete activation or differentiation subtypes may grossly oversimplify the phenotypic complexity of T cell populations resident in tissues.

### Diverse environmental stimuli and covariance of gene expression define intratumoral T cell states

Noting that the strongest components of variation do not fullly explain cluster distinctness, we sought to understand the variation driving the observed clustering. While clusters were arranged in a continuous fashion along the activation component, each cluster appeared distinct when accounting for a combination of signatures representing responses to diverse environmental stimuli. Our data show that CD4 effector and central memory clusters (Figure 5A) exhibit variable levels of gene expression involved in type I and II interferon response (F-test, p=1e-54 and 0.008 respectively), hypoxia (F-test, p=4e-64), and anergy (F-test p=4e-69). Moreover, CD8 effector and central memory clusters (Figure 5B) varied in expression levels of activation (F-test p=2e-114), proinflammatory (F-test p=1e-39), and cytolytic effector pathway-related genes (F-test p=6e-32). These findings suggest that tumor-resident T cells are exposed to varying degrees of inflammation, hypoxia, and nutrient deprivation. While many of these responses (e.g. activation or hypoxia) individually represent phenotypic continuums, their combinations may result in more discrete states (STAR Methods).

**Figure 5.**
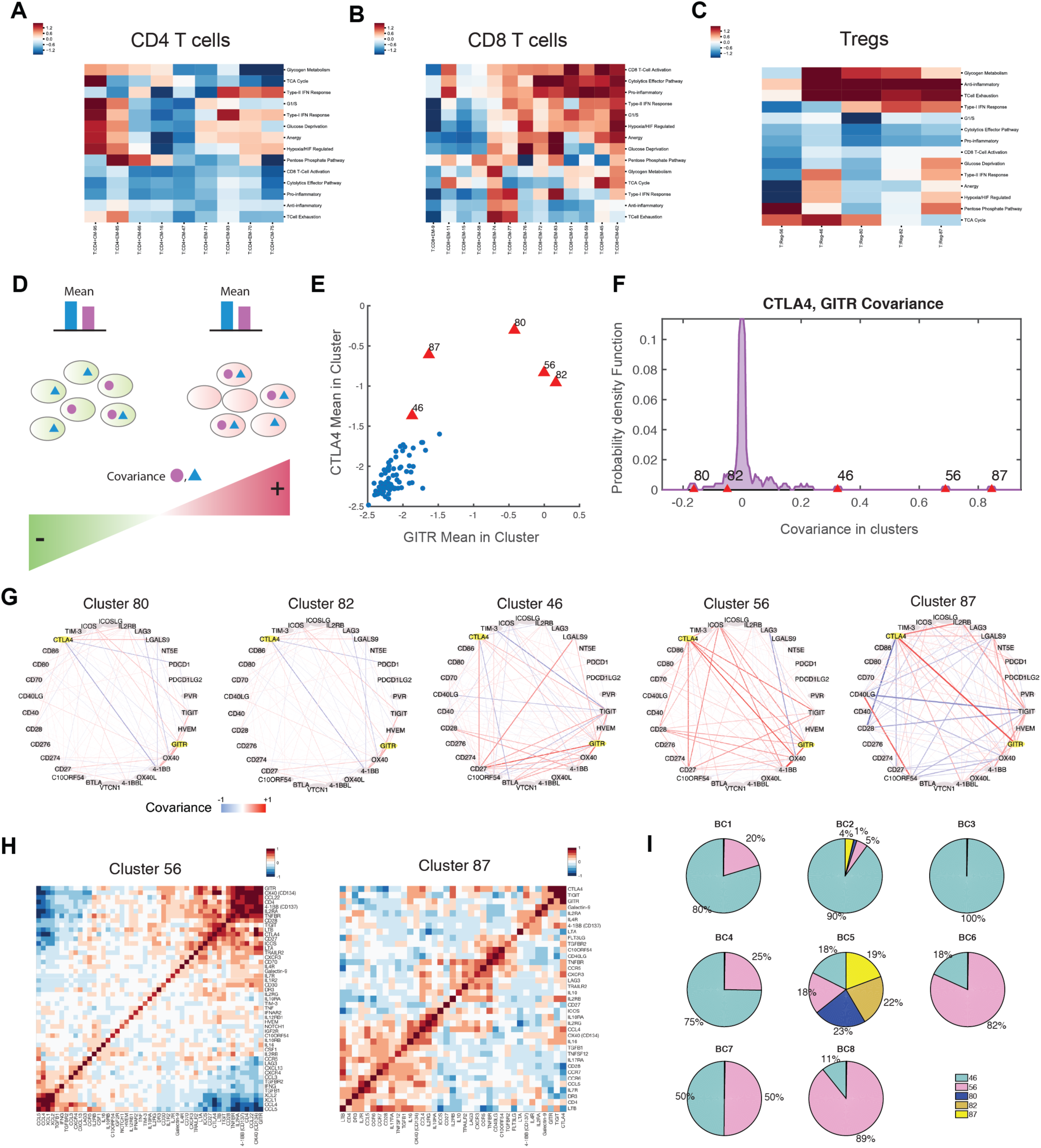
Covariance Patterns Help Define Distinct T Cell Clusters. (A, B, C) Heatmaps showing mean expression levels for a curated set of transcriptomic signatures (listed in Table S4) for (A) CD4 memory clusters, (B) CD8 memory clusters, and (C) Treg clusters. Only signatures with high expression in at least one T cell cluster are shown. Expression values are z-scored relative to all T cell clusters. (D) Cartoon illustration of two clusters of cells with similar mean expression for two example genes but opposite covariance between the same two genes. (E) Scatter plot showing mean expression of GITR vs. CTLA-4 for each T cell cluster (represented by a dot). Treg clusters, labeled in red, have high mean expression levels of both genes. (F) Distribution of covariance between GITR and CTLA-4 across all T cell clusters, with Treg clusters labeled in red. Note that Treg cluster covariance values exhibit differences despite sharing high mean expression levels. See Figure S5A for similar computation on the raw, un-normalized, data, verifying the result. (G) Network visualization illustrating strength of covariance between pairs of checkpoint receptor genes in Treg clusters. Edge width denotes absolute magnitude of covariance and red and blue colors denote positive and negative signs of covariance respectively; the case of CTLA-4 and GITR is highlighted in yellow. Distinct patterns in gene covariance, in addition to mean expression, define clusters in the Biscuit model. See figure S5C for networks for other T cell populations. (H) Heatmaps showing covariance between immune genes in two Treg clusters with different modules of covarying genes. (I) Proportion of the five Treg clusters in each patient, indicating that differences in covariance patterns between clusters translate to patients.

In contrast to effector T cells, Treg clusters displayed less variation in these gene signatures: the majority of these clusters featured comparable patterns for anti-inflammatory activity, exhaustion, hypoxia, and metabolism gene sets (Figure 5C). To identify features distinguishing the Treg clusters, we examined the Biscuit parameters differentiating them. We found that beyond mean expression levels, covariance parameters varied significantly between clusters. Specifically, two marker genes can exhibit similar mean expression in two different clusters (e.g. highly expressed in both), while the clusters show opposite signs in covariance between these genes. This occurs due to the genes typically being co-expressed in the same cells in one cluster (i.e. positive covariance), but expressed in a mutually exclusive manner in the other cluster (i.e. negative covariance) (Fig. 5D). It is noteworthy that clusters were inferred based on the expression of over 14,000 genes; hence, negative covariance between two specific genes does not necessarily imply the existence of sub-clusters.

As an example, our analysis showed that the prototypical co-inhibitory gene CTLA-4 exhibited rich covariance patterns with other mechanistically related genes (Figure 5E-G; S5A, B). CTLA-4 co-varied strongly with TIGIT and co-stimulatory receptor GITR in Treg clusters 46, 56, and 87; with CD27 in clusters 46 and 80; and with co-stimulatory receptor ICOS only in cluster 80 (Figure 5F,G). Covariance patterns between checkpoint receptors generally varied across Treg clusters (Figure 5G), and other important immune genes exhibited modular covariance structures, suggesting co-regulation and potential involvement in similar functional modalities (Figure 5H). Since varied proportions of Treg clusters were observed in individual patient samples, the differences in gene co-expression were also present at the patient level, and the majority of patients did not have all 5 subtypes of Treg cells (Figure 5I). We observed similar patterns for GITR and CTLA-4 in three additional breast tumors profiled using CyTOF mass cytometry: two Treg clusters resembled clusters 82 and 46 in terms of covariance and differentially expressed genes (Figure S5C; STAR). We also observed differences in co-variation patterns across activated T cell clusters (Figure S5D). Thus, co-variation of genes has a role in defining T cell clusters, in particular Treg clusters (Figure 5G,H, SF2).

### Paired single-cell RNA and TCR sequencing reveals the range of activation states of individual T cell clonotypes

One plausible explanation for the observed continuity of intratumoral T cell activation is exposure to increasingly diverse environments. A non-mutually exclusive hypothesis is that the wide range of signal strengths afforded by a diverse repertoire of TCRs can result in a continuous spectrum of T cell activation, obscuring the transitional states. Unlike polyclonal T cell populations, activation of a monoclonal T cell population with a fixed specificity for tumor neo-antigen may yield sparse discontinuous phenotypic spaces. Supporting the latter possibility, recent bulk gene expression and chromatin accessibility analyses showed that cognate tumor neo-antigen recognition by TCR transgenic T cells results in an orderly progression of activated T cells through a reversible dysfunctional intermediate state towards an irreversible dysfunctional terminal state (Philip et al., 2017).

To gain deeper insight into whether TCR repertoire diversity contributes to the observed spectrum of T cell activation, as well as overall phenotypic diversity, we profiled paired single-cell transcriptomes and TCRs (see STAR Methods). We collected single-cell RNA-seq and paired VDJ sequencing from over 27,000 sorted CD3^+^ T cells from three additional breast cancer tumors (labeled BC9-11; Figure S6A). This data allowed direct mapping of gene expression to TCR utilization by the same individual cells at an unprecedented scale.

The transcriptomic data further provided the ability to test the generalizability of the inferred clusters to three new patients profiled with a different single cell platform. This analysis revealed that T cell clusters, identified using Biscuit on the pooled dataset generated using the 10x platform, exhibited near one-to-one mapping to the T cell clusters inferred from the inDrop dataset (Figure 6A,B; S6B).

**Figure 6:**
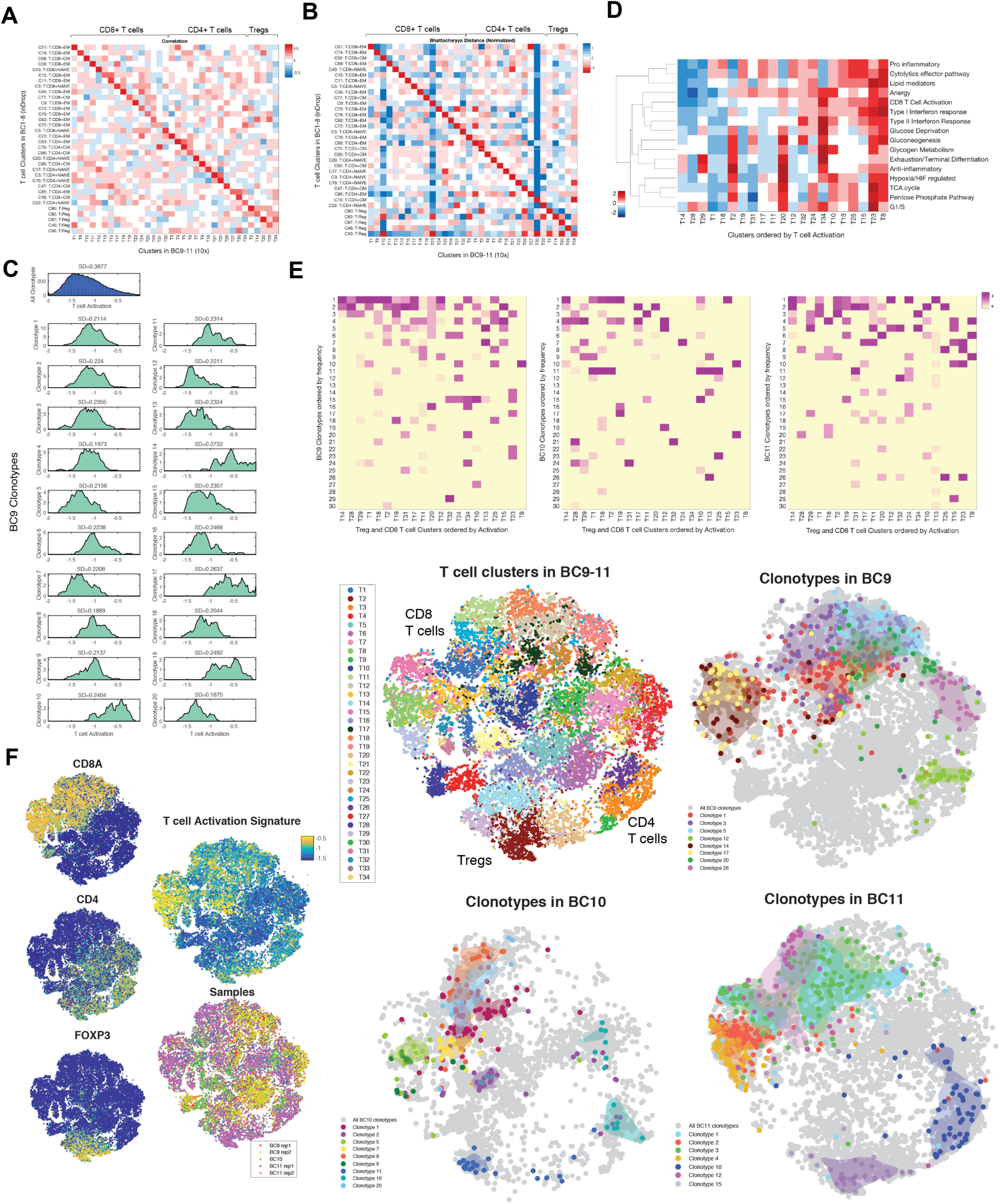
TCR repertoire shapes diverse phenotypic states. (A) Pearson correlation between centroids of differentially expressed genes in T cell clusters inferred from BC1-8 tumors using inDrop (rows) and clusters inferred from 27,000 T cells from BC9-11 tumors using 10x (columns) (see Figure S6A, B and STAR for details), showing near one-one-to-one mapping of clusters, indicating generalizability of the atlas. (B) Same as in (A) computing Bhattacharyya similarity between pairs of clusters, accounting for both mean and covariance of clusters. Further details on clusters are presented in S6B, C. (C) Histogram of activation states of (top) all T cells from three breast tumors BC9-11 and (bottom) T cells separated by each of the top 20 most dominant TCR clonotypes in BC9, mapped using paired single-cell RNA and TCR sequencing. Frequencies of clonotypes are shown in S6D. Similar figures for tumors BC10,11 are shown in Figure S6E. (D) Heatmap showing normalized mean expression levels for a curated set of transcriptomic signatures (rows, listed in Table S4) for T cell clusters found in BC9-11. (E) Distributions of each CD8+ T or Treg cluster in BC9-11 across the 30 most frequent clonotypes from each tumor. Columns (clusters) are z-scored to highlight the combinatorial impact of clonotypes in shaping each phenotypic state, and sorted by activation level. Some clusters associated with the same clonotype have the same level of activation (seen as connected horizontal stretches in the heatmap), while others have similar environmental responses. (F) t-SNE projection of normalized single-cell RNA-seq data for T cells from 3 tumors colored by markers, T cell activation signature, and patient source (left); Biscuit clusters (middle, top); and examples of dominant clonotypes from each tumor identified with paired TCR sequencing, projected in the same coordinates (right). Separate projection of each dominant clonotype for all three tumors is shown in Figure S6F.

This analysis also reproduced a continuous gradient along the T cell activation trajectory (Figure S6C), similar to that seen in Figure 4D. To evaluate the degree to which TCR diversity explains this continuity, we mapped the activation state of each clonotype separately using the paired data (Figure 6C; S6D). This revealed that a subset of the clonotypes exhibit distinct average activation levels. Further, we observed that the distribution of activation states found within any given clonotype is significantly constrained (i.e. has a higher entropy than random subsamples with size equal to that of the clonotype; p = 0, STAR Methods). This provides evidence that TCR diversity partially explains the observed continuity of T cell activation.

Nevertheless, a surprisingly wide range of activation states is present in each individual clonotype (Figure 6C; S6E). To be precise, 52%, 48%, and 32% of the variation in activation states across all cells can be explained by clonotype identities in BC9, BC10, and BC11, respectively (one-way ANOVA p<0.001, STAR Methods). Moreover, the average pairwise variation between the top 20 most frequent clonotypes is 54%, 46%, and 29% of the average variation within a clonotype in BC9, BC10, and BC11, respectively (F-test p<0.001, STAR Methods). These results together suggest that TCR diversity is not the exclusive driver of the continuity of T cell activation, implying the contribution of other factors.

### TCR repertoire shapes diverse phenotypic states in T cells

While we observed that the primary components explaining variation across immune cells, such as activation, exhibited continuity, the analysis of the entire high-dimensional data revealed discrete clusters (Figure S2B). Specifically, T cell clusters were separable on the basis of their differential expression of signatures associated with responses to environmental stimuli (Figure 5A,B). A similar trend was observed for tumor-resident T cells profiled using 10x technology (Figure 6D).

Interestingly, when analyzed jointly with TCR clonotypes, we observed that each cluster was in fact comprised of different combinatorial subsets of the clonotypes (Figure 6E). This observation provides further support for the distinctiveness of clusters, in part shaped by their TCR repertoires.

Furthermore, each clonotype was present only in a small number of related clusters that were significantly more similar phenotypically than randomly selected clusters (p<0.01 STAR Methods), and hence occupied a confined region in a 2D t-SNE projection (Figures 6F, S6F). In some cases clusters that shared a clonotype exhibited similar levels of activation (Figure 6E), whereas in other cases such clusters were similar in signatures relating to environmental stimuli. For example, clonotype 9 from tumor BC9 is present in clusters T11 and T12, which have very similar expression levels across nearly all of the environmental signatures (Figure 6D,E). Additionally, clonotype 20 in BC9 is present in clusters high in anergy and glucogenesis, and clonotype 21 in BC10 is present in clusters with high expression of the G1/S signature and low expression of other signatures.

The variable TCR clonotype composition of individual T cell clusters, together with their differential expression of key gene expression signatures, thus suggest that phenotypic states are shaped by a combination of antigenic TCR stimulation and environmental stimuli.

### Activation and differentiation define components of variation of intratumoral myeloid cells

Although myeloid lineage cells are commonly thought to be highly diverse, the heterogeneity of intratumoral monocytes and macrophages and its impact on tumor progression remains insufficiently characterized (Engblom et al., 2016). A broad survey of the monocytic clusters found in BC1-8 suggested unexplored substructure within these major cell types (Figure 7A). As with the T cells, we employed diffusion maps to assess this heterogeneity, excluding neutrophils and mast cells, as they formed much more distinct clusters (Figure 7B). This analysis revealed four major branches that displayed more distinct cell states than did the T cells (Figure S7A). The first branch almost entirely comprised intratumoral macrophages (TAMs) from three clusters (23, 25, and 28) (Figure 7B-F). The next two informative components capture a more gradual trajectory from blood monocytes to intratumoral monocytes (Figure 7B-D; S7B-D); an additional component with two discrete states distinguishes plasmacytoid DC (pDC) from the other monocytic cell clusters (Figure 7B, E; S7B, C, E).

**Figure 7:**
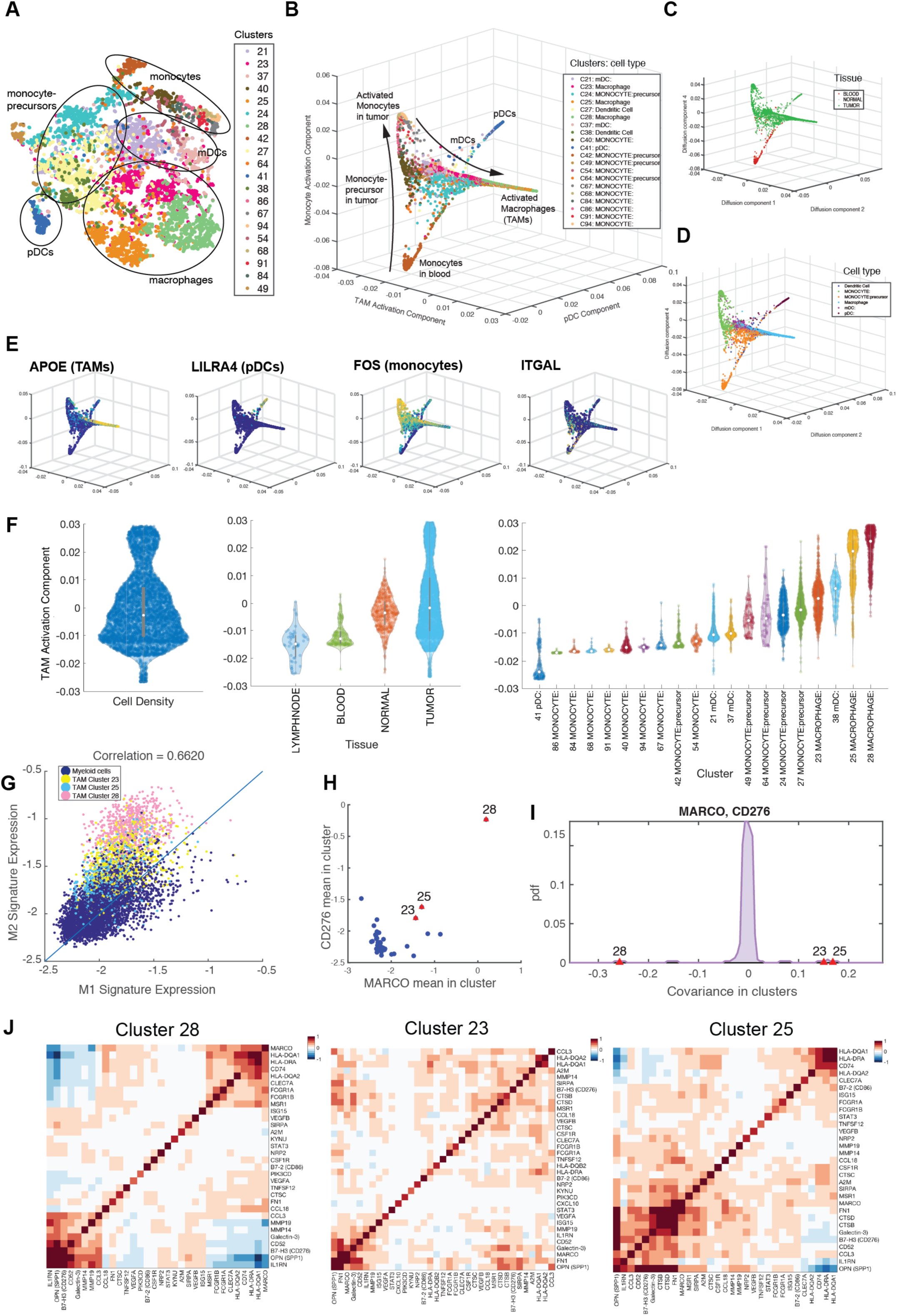
Detailed Characterization of Myeloid Cells. (A) t-SNE map projecting only myeloid cells across tissues and patients (BC1-8). Cells are colored by Biscuit cluster and cell types are labeled based on bulk RNA-seq correlation-based annotations. (B through E) Projection of myeloid cells on macrophage activation, pDC, and monocyte activation diffusion components, colored by (B) cluster, (C) tissue type, (D) cell type, and (E) expression of example lineage demarcating genes. (F) Violin plots showing the density of cells along macrophage activation component and organized by overall density (left panel), tissue type (middle panel), and cluster (right panel). See Figure S6 for other components. (G) Scatter plot of normalized mean expression of M1 and M2 signatures per cell; cells assigned to 3 TAM clusters have been highlighted by cluster; each dot represents a cell and cells are plotted in randomized order. (H) Scatterplot of mean expression of MARCO and CD276 in myeloid clusters; each dot represents a cluster. Average expression levels for the TAM clusters are marked in red, indicating high expression of both markers in macrophage clusters. (I) Distribution of covariance between MARCO and CD276 across all myeloid clusters. TAM clusters are marked in red and present substantial outliers. See Figure S7B for similar computation on the raw, un-normalized data, verifying the result. (J) Heatmaps showing covariance patterns of M1 and M2 macrophage polarization marker genes (including many current or potential drug targets) in 3 TAM clusters.

Focusing on the first branch, top correlated genes include APOE, CD68, TREM2, and CHIT1 (Figure S7B). This likely reflects differentiation and activation of either recruited or tissue-resident macrophages. Additionally, expression of genes associated with “alternatively activated” (M2) macrophages, including scavenger receptor MARCO, pro-angiogenic receptor NRP2, and inhibitory molecule B7-H3 (CD276), increased along this branch (Figure S7B). Concomitantly, immunostimulatory genes associated with “classically activated” (M1) macrophages, including chemokine CCL3 (MIP-1a), increased along the branch. All three of the TAM populations, particularly clusters 23 and 28, were among the monocytic clusters with the highest expression of the canonical M2 signature, but were likewise high in the M1 signature (Figure S7E). Quite strikingly, we found that M1 and M2 gene signatures positively correlated in the myeloid populations (Figure 7G), in line with recent findings in other tumor types (Muller et al., 2017). These findings support the idea that macrophage activation in the tumor microenvironment does not comport with the polarization model, either as discrete states or along a spectrum of alternative polarization trajectories.

Similar to the T cells, covariance parameters were key in differentiating the three TAM populations, though they shared most of their differentially expressed genes (Table S3). One example was co-expression of two M2-type markers, MARCO and B7-H3. While the TAM clusters all expressed high levels of both genes, they co-varied positively in clusters 23 and 25, but negatively in 28 (p = 0, p = 5e-06, p = 0, respectively; Figure 7H, I; S7F,G). The differing covariance patterns were also significant in raw un-normalized data (Figure S7F) (STAR Methods) and thus not an artifact of modeling. Such examples highlight the importance of co-expression patterns in defining myeloid cell states.

## DISCUSSION

Despite major clinical advances in cancer immunotherapy, our ability to understand mechanisms of action or predict efficacy is confounded by the heterogeneous and poorly understood composition of immune cells within tumors. Using unbiased single-cell RNA-seq analysis of all tumor vs. normal tissue-resident immune cell subsets we constructed an immune atlas in breast carcinomas, combining immune cells isolated from normal and cancerous breast tissue, as well as peripheral blood and the lymph node.

This atlas revealed vast diversity in immune cells of both the adaptive and innate immune systems, with the biggest change linked to the tissue environment. Interestingly, immune cell subpopulations in normal tissue were observed to be a subset of those found in tumor tissue, an observation that could not have been found with bulk gene expression measurements.

Furthermore, the diversity of cell states, as quantified with our metric of “phenotypic volume”, significantly expanded in breast tumors as compared to normal breast tissue. Three major components - T cell activation, terminal differentiation, and hypoxic response - contributed to this phenotypic expansion. The activation component argues against a view of activated T cells rapidly traversing through sparse transitional cell states towards a few predominant, discrete, and stable states, including Treg, effector, memory, and exhausted T cells. This continuous view of T cell differentiation comports with recent work demonstrating that key gene signatures in CNS-autoantigen-specific Th17 cells exhibit a continuous spectrum of cellular states, which help discern non-pathogenic and pathogenic modalities of this effector T cell type (Gaublomme et al., 2015).

TCR diversity accounts in part for the continuous spectrum of T cell activation, though a wide range of activation states is also found within each TCR clonotype. This continuity may also be attributable to asynchrony in polyclonal T cell activation or heterogeneity in the types of antigen-presenting cells, their activation status, and their anatomical distribution.

When analyzed in conjunction with TCR utilization, we found T cell populations to be associated with unique combinations of TCR clonotypes. These TCR usage patterns, together with unique environmental exposures, jointly define the discrete states of intratumoral T cells. This suggests two models to explain the tight association between T cell clonotype and transcriptional phenotype. One, TCR signaling is a strong driver of downstream transcriptional signaling, and may be sufficient to account for the restricted range of phenotypes observed within each clonotype (Figure 7E, S7D). This model has recently been proposed specifically in the case of Tregs (Zenmour et al., 2018). Alternatively, diverse TCR specificities may contribute to the spatial distribution of T cells on account of the distribution of their cognate antigens and, therefore, facilitate their exposure to the aforementioned distinct environments (“mini-niches”). Recent work using multiregional genomic sequencing of tumors has revealed a high degree of tumoral subclonality diverging between spatial regions (Sun et al., 2017), including in breast cancer (Yates et al., 2015). In support of this notion, our inferred subsets showed variable levels of responses to environmental stimuli. The quality and quantity of antigen presentation is likewise known to vary over time, as tumors respond to selective pressure by downregulating expression of both MHC complexes and individual neoantigens (Verdegaal et al., 2016).

Beyond these environmental responses, complex co-expression patterns also contributed to defining T cell states. Five Treg subsets exhibited similar mean gene expression for canonical markers, but exhibited drastic differences in gene covariance patterns. Particularly noteworthy was co-expression of checkpoint receptor genes in some Treg subpopulations as compared to mutually exclusive expression of the same genes in other Treg clusters. In this regard, varying co-expression of CTLA-4, TIGIT, and GITR and other co-receptors in multiple Treg clusters suggests that these populations may occupy different functional niches. Cells co-expressing CTLA-4 and TIGIT have been demonstrated to selectively inhibit pro-inflammatory Th1 and Th17 responses but not Th2 responses, promoting tissue remodeling (Joller et al., 2014). We also observed considerably different proportions of Treg clusters across patients, suggesting that multi-dimensional profiling might be necessary to personalize future combination therapies.

Our analyses appear to offer a more nuanced view of tumor and normal tissue-resident myeloid lineage cells in comparison to T cells, in terms of continuity vs. separation of cell states. Unlike T cells, which primarily displayed continuous activation transitions, we observed sharper state delineations in myeloid populations. This difference between T cells and myeloid cells was likely due to a less appreciated developmentally established myeloid cell heterogeneity, whose understanding has begun to emerge only recently (Perdiguero and Geissmann, 2016). However, our analyses also showed common features to those in T cells, including gene expression covariance identifying cell clusters and an expansion of immune phenotypic space in breast tumor as compared to normal breast tissue.

In macrophages, we found both M1 and M2 associated genes frequently expressed in the same cells, positively correlated with one another and following the same activation trajectory. These results challenge not only the customary model of macrophage polarization wherein M1 and M2 activation states exist as mutually exclusive discrete states, but also a refined model wherein macrophages reside along a spectrum between the two states. In fact, our data goes further than models that admit for the co-existence of M1 and M2 states (Martinez and Gordon, 2014), demonstrating a positive correlation between the two. Our findings solidify and reinforce previously reported bulk analyses of tumor-associated macrophages in mouse models of oncogene-driven breast cancer, and mass cytometry analyses of myeloid cells in lung and kidney cancer (Chevrier et al., 2017; Franklin et al., 2014; Lavin et al., 2017). Notably, we observed more patient-specific variation in myeloid lineage cells than in T cells, with the frequency of the former ranging from just over 10% to over 50% in individual patients. Individual clusters similarly exhibited ranges of patient specificity. The large patient effect in myeloid cells suggests that attempts at generalized targeting or reprogramming of suppressive myeloid cell populations may not yield uniform responses and personalization at the patient level may be needed.

Our characterization of the immune cell subsets inhabiting primary solid tumor and the corresponding normal tissue, and their heterogeneity within and between patients, revealed continuity of differentiation states and expansions of a “phenotypic space” as principal features of the two main cellular targets of cancer immunotherapy - T cells and myeloid cells. In T cells these features were in large part shaped by TCR-induced activation and TCR-dependent environmental exposures. These observations, along with the resulting extensive immune single-cell RNA- and TCR-seq datasets and comprehensive analytical platform, will facilitate better understanding of potential mechanisms behind immune cell contributions to promoting and opposing tumor progression.

## Author Contributions

E.A., AJ.C., G.P., A.E.C., C.K., A.Y.R, D.P. conceived the study. A.J.C., M.S., K.C, P.D., D.P. designed and developed SEQC. E.A., S.P., D.P. designed and developed Biscuit. G.P. provided clinical samples. K.W., R.M.F., P.M. prepared samples. J.N., V.K., L.M. performed all scRNA-seq data acquisition experiments. P.T.M., R.C.W., K.K. collected CyTOF data. E.A, A.E.C., D.P. developed new analysis methods. E.A., A.J.C., G.P., A.E.C., C.K., A.Y.R, D.P. analyzed and interpreted single-cell RNA-seq data. E.A., G.P., A.E.C., A.Y.R, D.P. analyzed and interpreted paired TCR and RNA-seq data as well as CyTOF data. E.A., A.J.C, A.E.C, A.Y.R., D.P. wrote the manuscript.

## Acknowledgements

We thank Barbara Engelhardt, David Blei, Frederic Geissmann, and Roshan Sharma for valuable conversations related to this manuscript. We thank the Flow Cytometry Core, Tissue Procurement Service, and Integrated Genomics Operation at MSKCC for assistance. We thank the specimen donors at MSKCC. This study was supported by NIH grants R37 AI034206 (A.Y.R), NIH DP1-HD084071 (D.P), NIH R01CA164729 (D.P). Cancer Center Support Grant P30 CA008748, the Hilton-Ludwig Cancer Prevention Initiative funded by the Conrad N. Hilton Foundation, Ludwig Cancer Research, and Gerry Center for Metastasis and Tumor Ecosystems. E.A. was supported by an American Cancer Society - Fairfield County Roast Committee Postdoctoral Fellowship (PF-17-243-01-RMC). A.J.C is a recipient of an HHMI international graduate fellowship. G.P. is the recipient of a Breast Cancer Alliance Young Investigator Grant and the Society of MSK Research Grant. C.K. was a recipient of a Hutham S. Olayan Graduate Fellowship. P.T.M. is the recipient of a Boehringer Ingelheim SHINE Fellowship. CyTOF experiments were supported with funds from the Boehringer Ingelheim SHINE Program (A.Y.R.). A.Y.R. is an investigator with the Howard Hughes Medical Institute.

---

**Figure S1:**
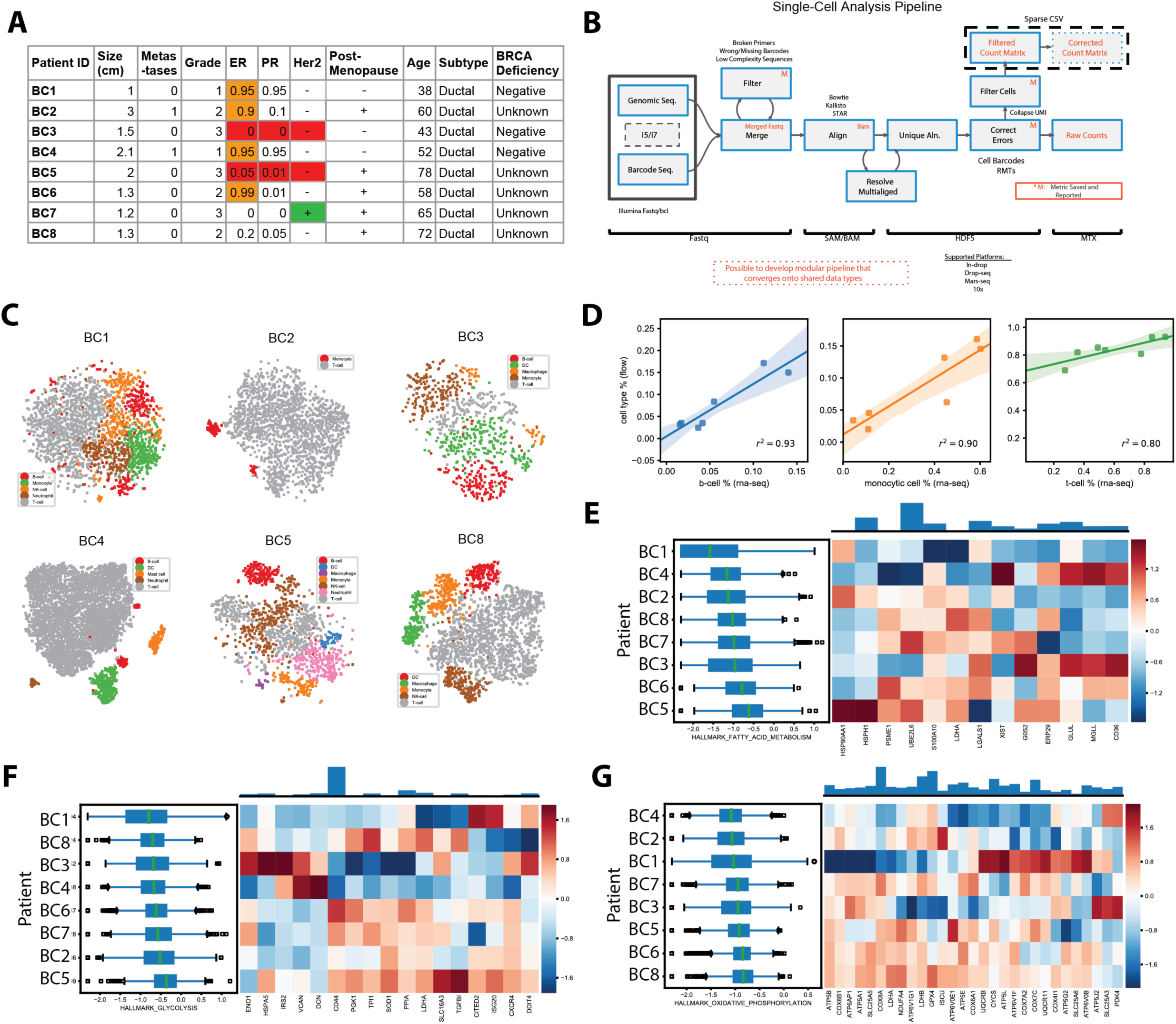
Additional details on samples and individual immune systems, related to Figure 1. (A) All clinical and related metadata for all 8 patients. (B) Bioinformatics pipeline for processing single-cell RNA-seq data; inputs boxed on left, outputs labeled in red. Each rectangle represents a processing step. Braces on bottom display file formats as data moves through the pipeline. Quality control and metrics are provided in Table S1. (C) t-SNE projection of complete immune systems from six breast cancer tumors. scRNA-seq data for each tumor is processed with pipeline in Figure S1B and library size-normalized; each dot represents a single-cell colored by PhenoGraph clustering, and clusters are labeled by inferred cell types. Two additional tumors are presented in Figure 1C. (D) Regression of flow cytometry cell type percentages in each patient against RNA-seq cell type percentages for B cells (blue), monocytic cells (orange), and T cells (green). (E-G) Expression of metabolic signatures: fatty acid metabolism (E), glycolysis (F), and phosphorylation (G), summarized as boxplots (left) showing expression of each respective signature (defined as the mean normalized expression of genes in each signature listed in Table S4) across immune cells from each patient; and heatmap (right) displaying z-scored mean expression of genes in each signature; (top) barplot showing total expression of each gene indicated in the heatmap across all patients. See Figure 1E for one additional signature.

**Figure S2:**
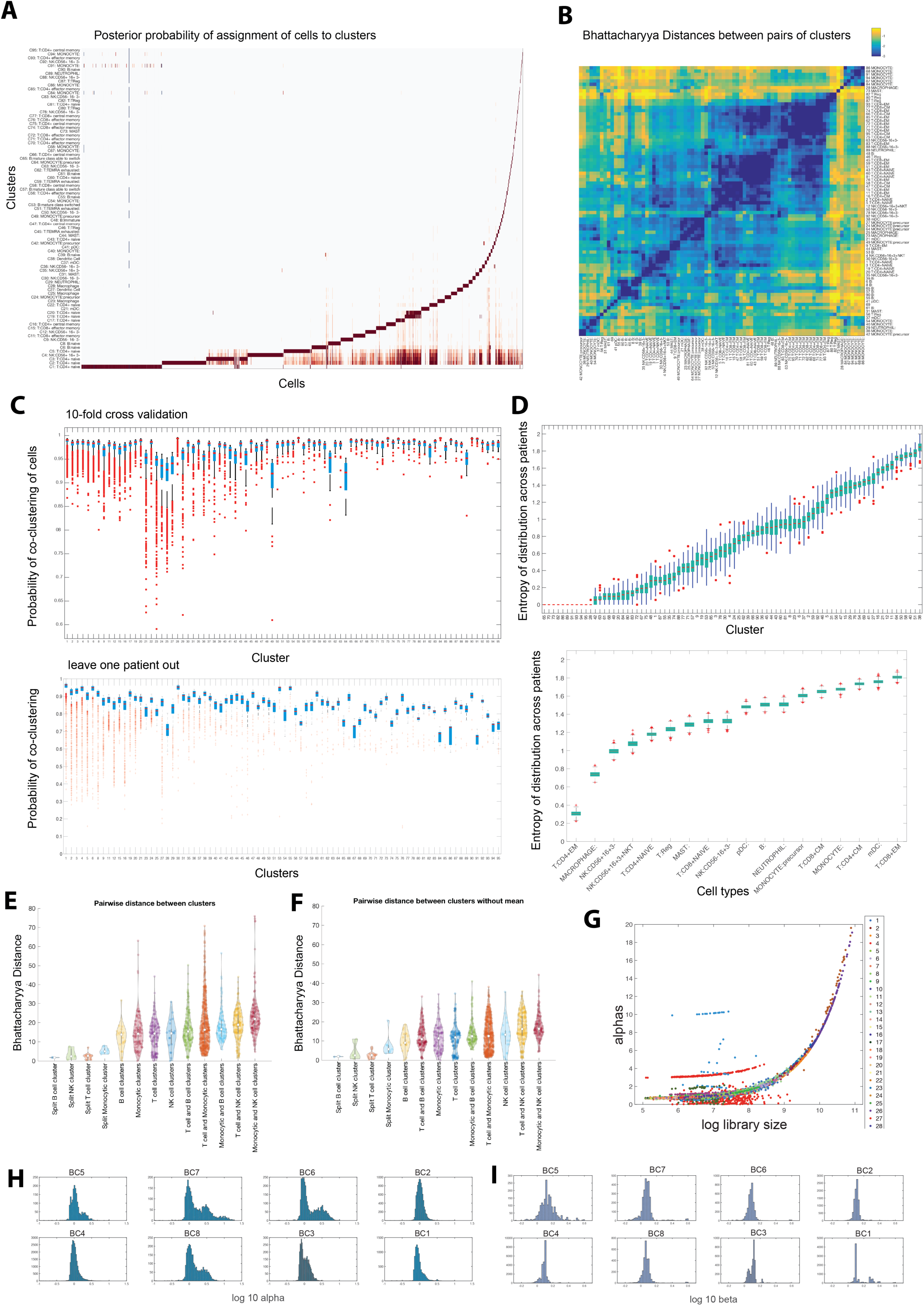
Details of inferred parameters for clustering and normalization using Biscuit, related to Figure 2. (A) Posterior probability of assignment of cells to clusters in the Biscuit model in the full immune cell atlas of combined tissues and patients presented in Figure 2E; note broad distributions in assignment of naive T cells (bottom) as compared to other cell types. (B) Bhattacharyya pairwise distances between clusters of Figure 2F, H (blue: small distance to yellow: large distance). (C) Robustness analysis of clusters performed with 10-fold cross-validation (top) and leave-one-patient-out cross-validation (bottom); boxplots summarize the probability of a pair of cells being assigned to the same final cluster across all 10 subsets/8 left-out patients. (D) Boxplots showing entropy of distribution of patients in each cluster (top) and cell type (below), computed with bootstrapping to correct for number of cells in each cluster/cell type. For clusters note that labels are ordered by size (cluster 1 has the most number of cells and cluster 95 the fewest) and ordering clusters by mean entropy in this plot indicates that entropy does not correlate with size. (E) Violin plot of pairwise Bhattacharyya distances between gene expression distribution for all pairs of clusters in the same or different cell types considering mean and covariance of expression, averaged across all genes. (F) Same as (E), but after removing the effect of cluster mean in computing similarity, thus considering only covariance. (G) Distribution of Biscuit alpha parameters per cell vs log of library size, with cells colored by clusters; Biscuit alpha parameters correct for differences in library size across and within clusters. (H-I) Distribution of inferred cell-specific parameters alpha (H) and beta (I) in Biscuit across cells from each patient. These differences were corrected in normalizing using these alpha and beta parameters.

**Figure S3.**
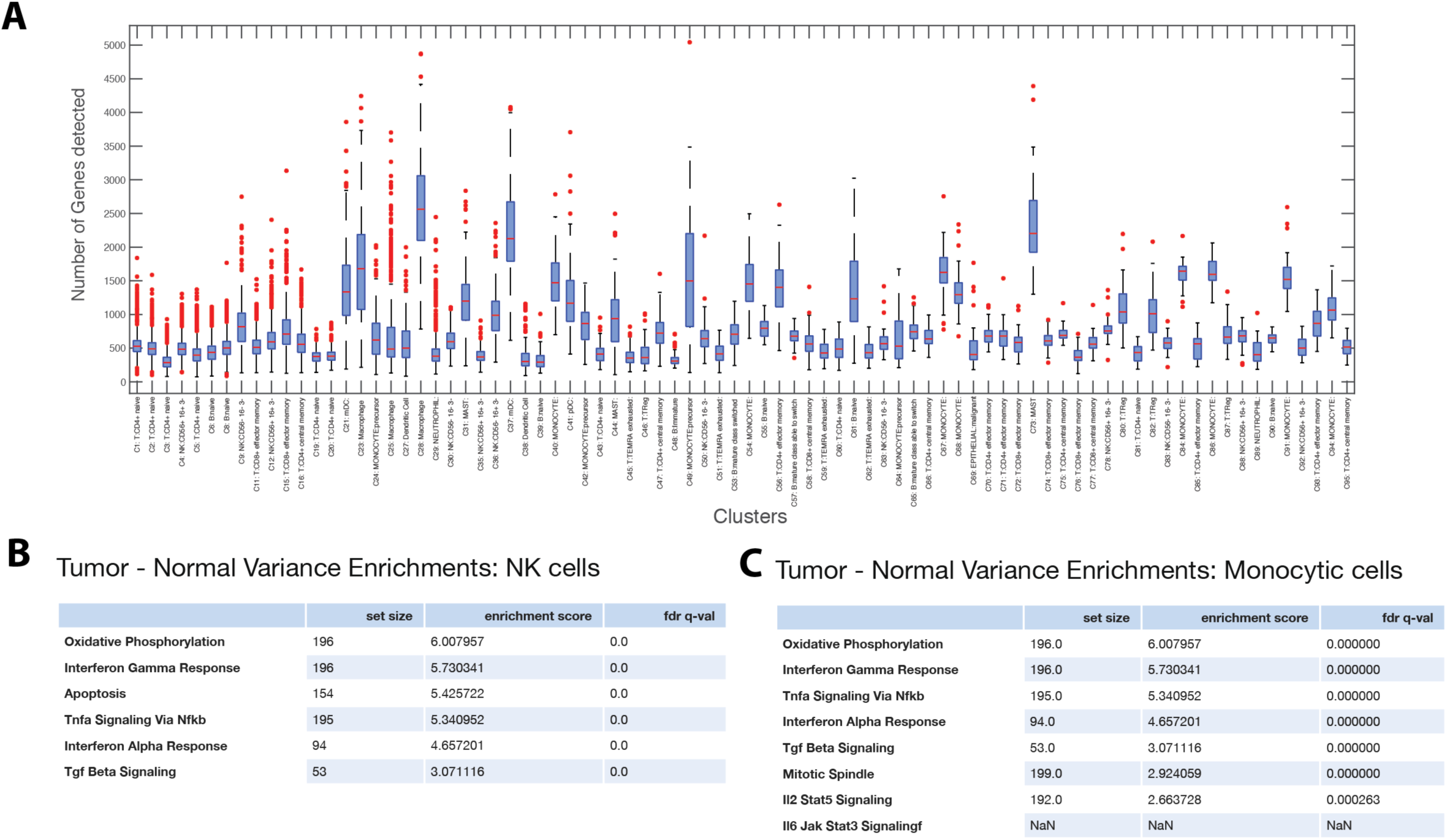
Analysis of differences in gene expression variance between immune cells in breast tumor compared to normal breast tissue, related to Figure 3. (A) Boxplot showing the number of genes detected per cell grouped by cluster. (B-C) Hallmark GSEA enrichment (Subramanian et al., 2005) results on genes with highest difference in variance in tumor vs normal tissue in (B) NK and (C) monocytic cells. See Figure 3E for enrichment in T cells; complete lists of enrichments are presented in Table S5.

**Figure S4.**
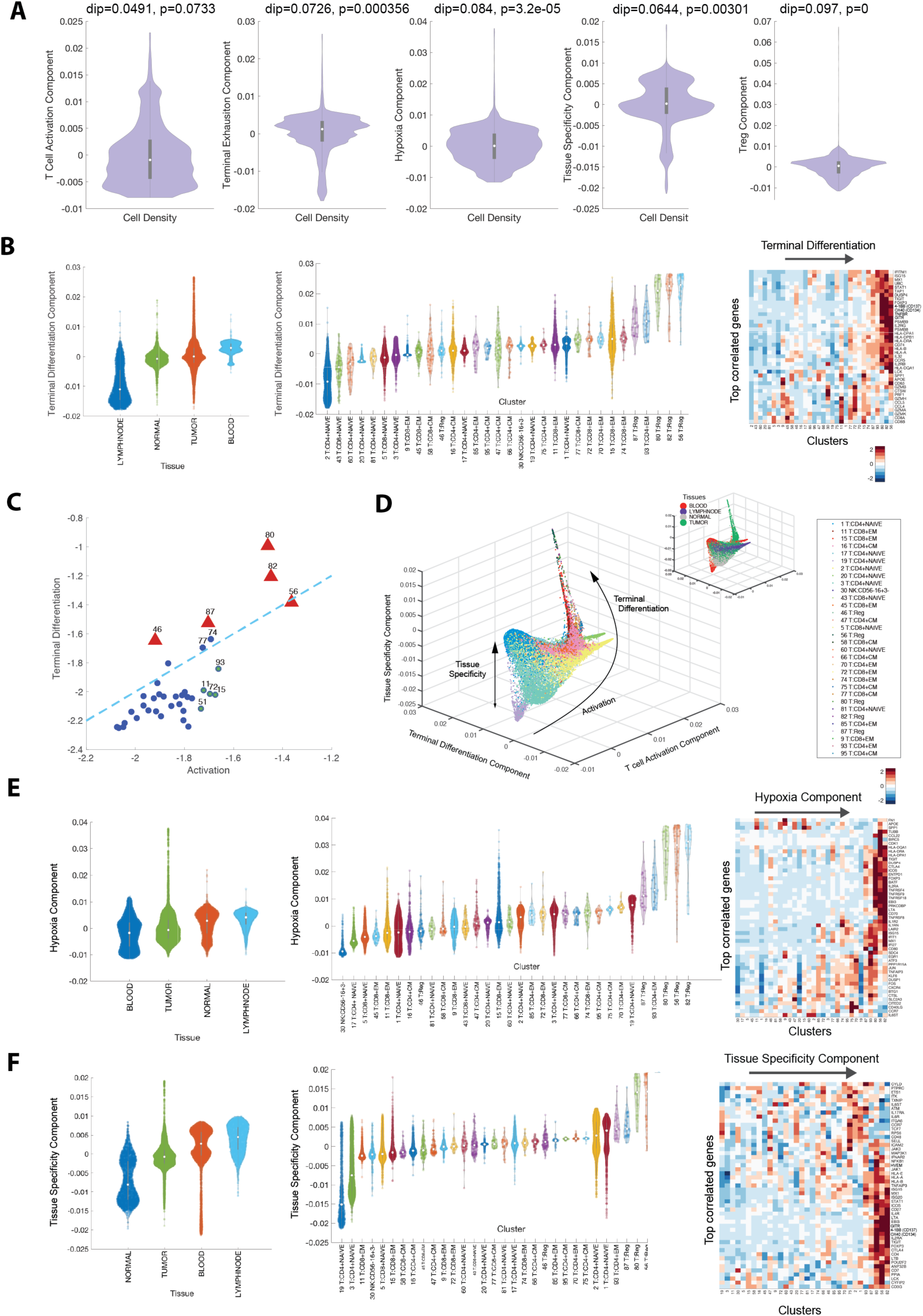
Additional details on T cell diffusion components, related to Figure 4. (A) Hartigan’s dip test on density of cells projected on diffusion components, showing statistically significant continuity (lack of “dips”) in cells along T cell activation component (component 3, third panel from left); other components exhibit more defined states (multimodality). (B) Violin plot of cells projected on terminal differentiation diffusion component (component 4): terminal differentiation component organized by tissue type (left panels) and cluster (center panel). Also, heatmap showing expression of immune-related genes with the largest positive correlations with component, averaged per cluster and z-score standardized across clusters; columns (clusters) are ordered by mean projection along the component. (C) Scatter plot showing mean expression levels of T Cell activation and terminal differentiation signatures for all T Cell clusters. Red triangles denote T Reg clusters showing high expression in both signatures. The dotted line denotes y = x. (D) Visualization of all T Cell clusters using activation (component 3), terminal differentiation (component 4), and tissue specificity (component 6) diffusion components. Cells are colored by clusters, and by tissue type in insert. The main trajectories are indicated with arrows and annotated with labels. (E,F) Same as (B) for hypoxia (E) and tissue specificity (F) components. See Figure 4 for additional component.

**Figure S5. Details of covariance patterns in T cell clusters, related to Figure 5.**

(A) Displaying null distributions and observed covariances between CTLA-4 and GITR in raw, un-normalized data using hypothesis testing, subsampling, and permutation (see STAR methods); shows that the differences in covariance shown in Figure 5F,G are also present in un-normalized and un-imputed data, and hence are not an artifact of computation.

(B) Bivariate plots of expression levels of GITR and CTLA-4 in Treg clusters based on inferred mean and covariance parameters from Biscuit. Dark blue color indicates the highest density of cells and light yellow the lowest density of cells.

(C) CyTOF data for 6556 T cells collected using panel presented in Table S9 from three tumors (BC12-14); top left: PhenoGraph clusters with two Treg clusters marked as A, B. Top right: Boxplots of expression of two markers in Treg cells in each cluster A, B. Bottom: Covariance between CTLA-4 and GITR in each Treg cluster A, B. Each dot is a cell, colored by density of cells. Cluster A resembles previous Treg cluster 82, differentially expressing CD25 and no covariance between CTLA-4 and GITR, while Treg cluster B resembles cluster 46, differentially expressing TIGIT and strong positive covariance between CTLA-4 and GITR.

(D) Network graphs showing covariance between checkpoint receptors in activated T cell clusters. Edge width denotes absolute magnitude (strength) of covariance and color denotes sign of covariance (red positive and blue negative). Note diversity across clusters. Similar graphs for T reg clusters are shown in 5G.

**Figure S6:**
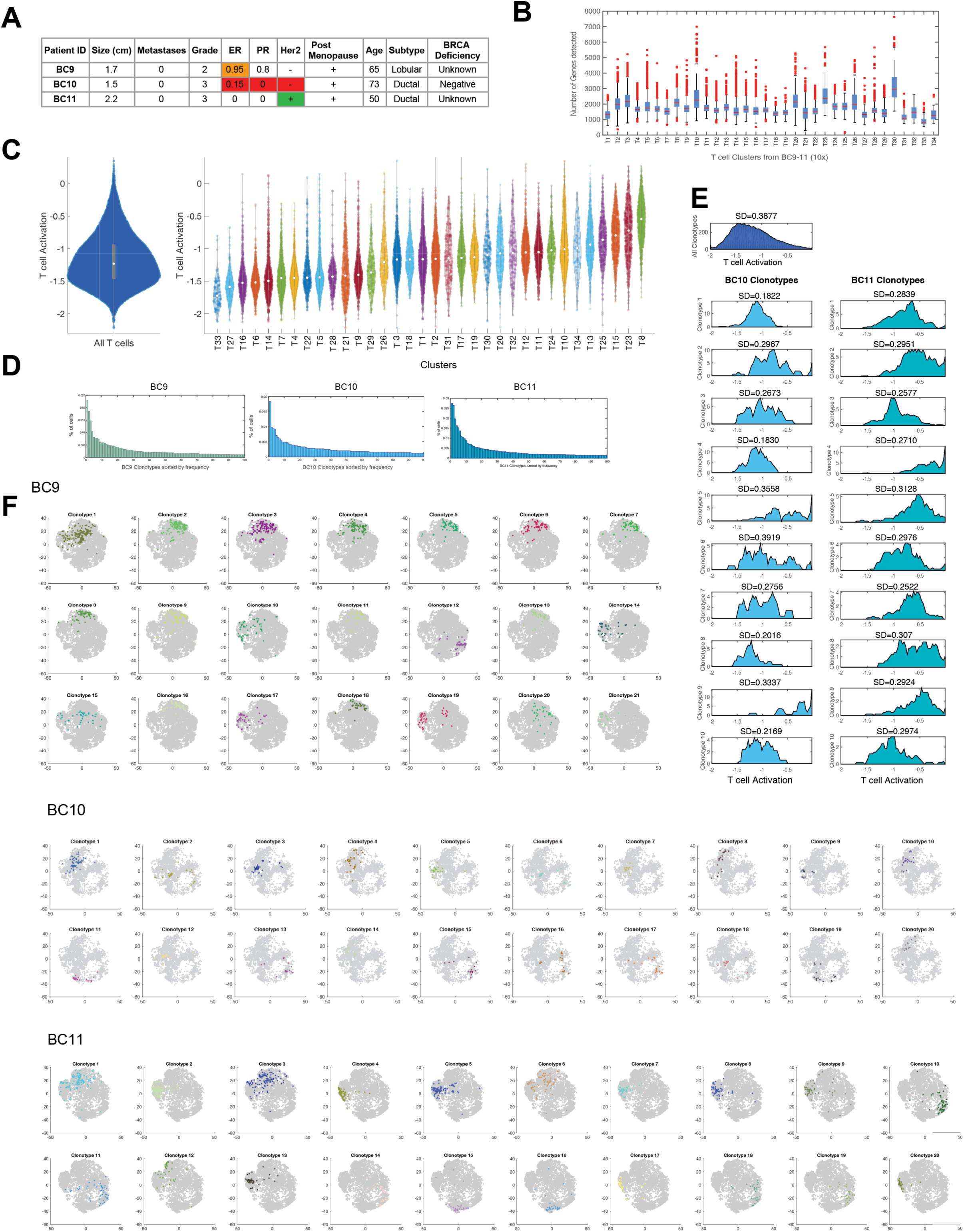
Additional Details on paired single-cell TCR sequencing and RNA-seq of T cells from 3 breast tumors, related to Figure 6. (A) Clinical and related metadata for 3 tumors (BC9-11) used for paired TCR and RNA-seq study. (B) Barplot showing the number of genes detected per cell grouped by clusters inferred from 27,000 CD3+ cells from three tumors, profiled with 10x 5’ single-cell RNA-seq technology and analyzed using Biscuit. (C) Violin plot showing the distribution of 27,000 T-cells from single-cell RNA-seq from three tumors (BC9-11) along activation signature aggregated by total density (left) and cluster (right). Number of dots inside each violin are proportional to number of cells. (D) Barplots showing frequencies of 100 most dominant clonotypes in each tumor. (E) Histogram of activation states of (top) all T cells from three breast tumors BC9-11 and (bottom) T cells separated by each of the top 10 most dominant TCR clonotypes in BC10 and BC11 mapped using paired single-cell RNA and TCR sequencing. Similar figures for tumors BC9 are shown in Figure 6C. (F) t-SNE projection of all clonotypes identified in each tumor (grey) and each of the most dominant clonotypes separately (in color); each dot is a T cell; coordinates are the same as in Figure 6F. Select dominant clonotypes from BC9 (top), BC10 (middle), and BC11 (bottom) spanning different regions of the 2D projection are overlaid in Figure 6F.

**Figure S7:**
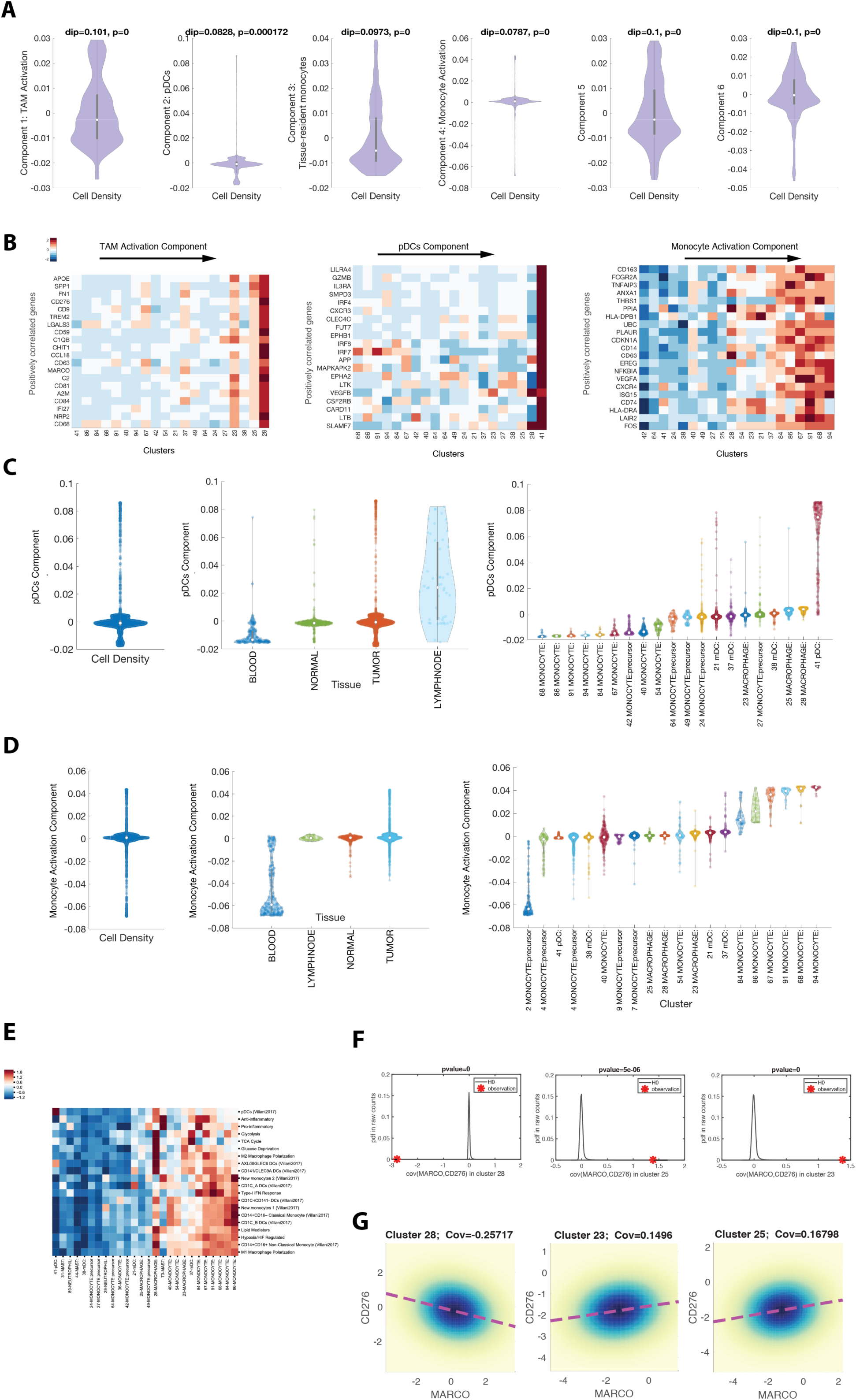
Additional details on diffusion component analysis of myeloid cells, related to Figure 7. (A) Hartigan’s dip test on density of cells projected on diffusion components: no diffusion components across myeloid cells show statistically significant continuity (unimodality), implying myeloid cells reside in defined (multimodal) states along major components explaining variation. (B) Heatmaps showing expression of immune-related markers with the largest positive correlation with TAM activation (left), pDCs (middle), and monocyte activation (right) components. (C) Violin plot showing the density of cells projected along pDC component and organized by tissue type and cluster. (D) Violin plot showing the density of cells projected along monocyte activation component and organized by tissue type and cluster. (E) Heatmap showing imputed mean expression levels in myeloid clusters for a curated set of transcriptomic signatures important to myeloid cells (listed in Table S4), z-score normalized per signature. (F) Displaying null distributions and observed covariances between MACRO and CD276 in raw, unnormalized data using hypothesis testing, subsampling, and permutation (see STAR methods), showing that the differences in covariance in normalized data as shown in Figure 7B are also present in un-normalized and un-imputed data, and hence are not an artifact of computation. (G) Bivariate plots of expression levels of MARCO and CD276 in Treg clusters based on inferred mean and covariance parameters from Biscuit. Dark blue color indicates the highest density of cells and light yellow the lowest density of cells.

**Methods Figure 1.**
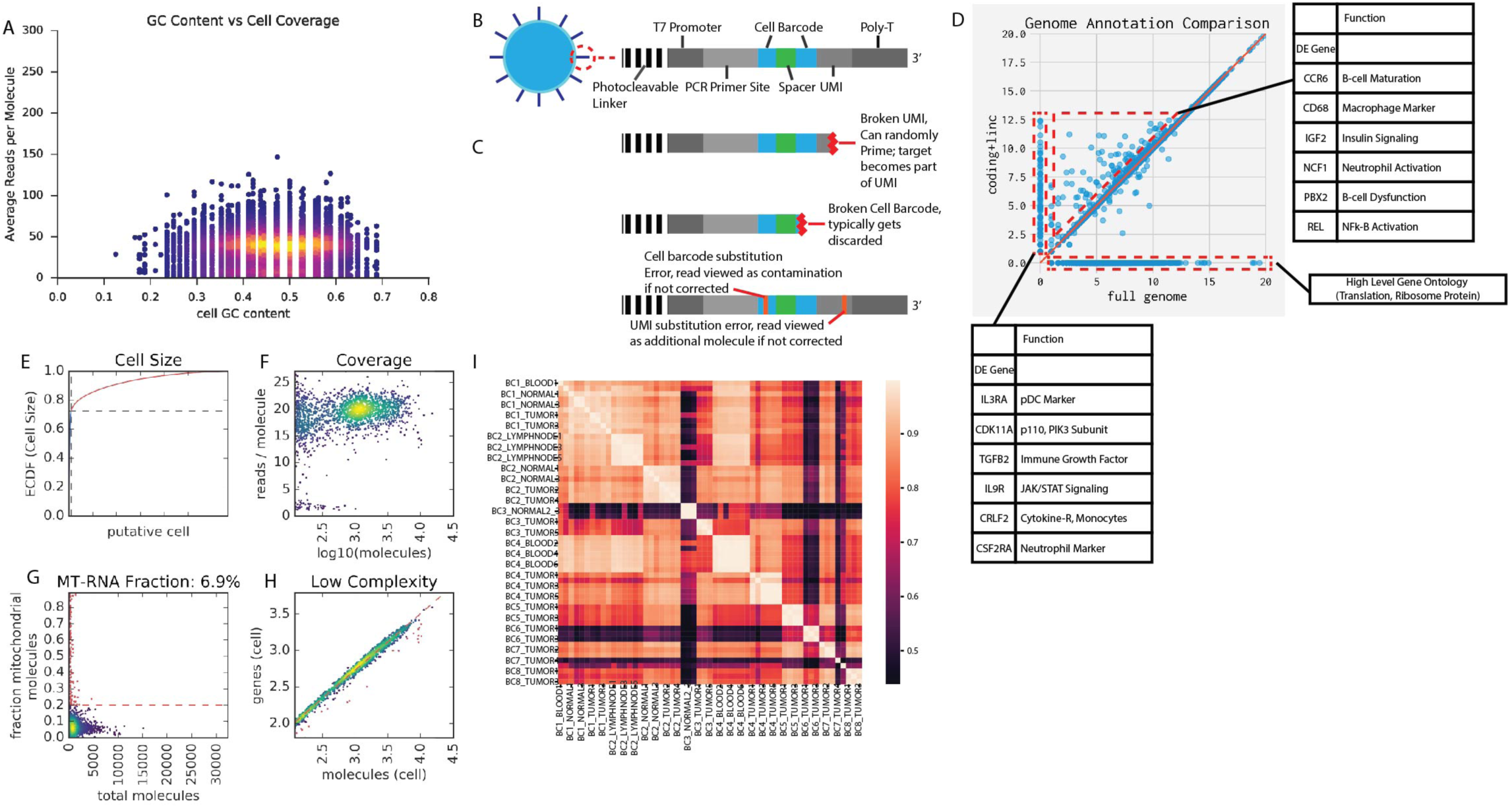
Data Driven Pipeline Construction SEQC. (A) Visualization of cell barcode GC content (percentage, x-axis) versus cell barcode read coverage (y) displaying higher coverage at 50% GC content. Yellow to purple color represents density of cell barcodes. (B) Schematic of capture primer displaying amplification machinery, cell barcodes, UMIs, and poly-T capture site. (C) Cartoon showing common sources of barcode errors including (i) breakage and (ii) substitution errors. (D) Comparison of complete GENCODE annotation against a reduced annotation containing only GENCODE-annotated lincRNA and protein coding RNA. Displaying drop-out events occurring on x-axis as well as masking events on y-axis. (E) Example cell filtering plot showing the empirical cumulative density of molecules (y-axis) per cell barcode (x-axis). Note that a small number of cell barcodes contain most of the molecules in the experiment. Dashed black lines represent cut-off points after which cell barcodes are considered to consist of contamination. Red barcodes are excluded. (F) Coverage plot comparing the total molecules in each cell (x axis) against the average coverage in each cell (y axis). Densities of cells with aberrantly low coverage such as those with lower than 5 reads / molecule are considered likely errors and are discarded. (G) Mitochondrial (MT) RNA fraction plot displaying the total number of molecules in each cell vs the fraction of those molecules that come from mitochondrial sources. Cells in red consist of more than 20% MT-RNA and are considered to be likely dying cells. These cells (red) are discarded. (H) Complexity plot displaying the number of detected molecules (x axis) vs genes (y axis). The relationship is fit with linear regression and cells (red) whose residual gene detection is greater than 3 standard deviations are removed. (I) Heatmap of pairwise pseudo-bulk sample-sample correlations (r2) across all samples and replicates in the experiment.

**Methods Figure 2.**
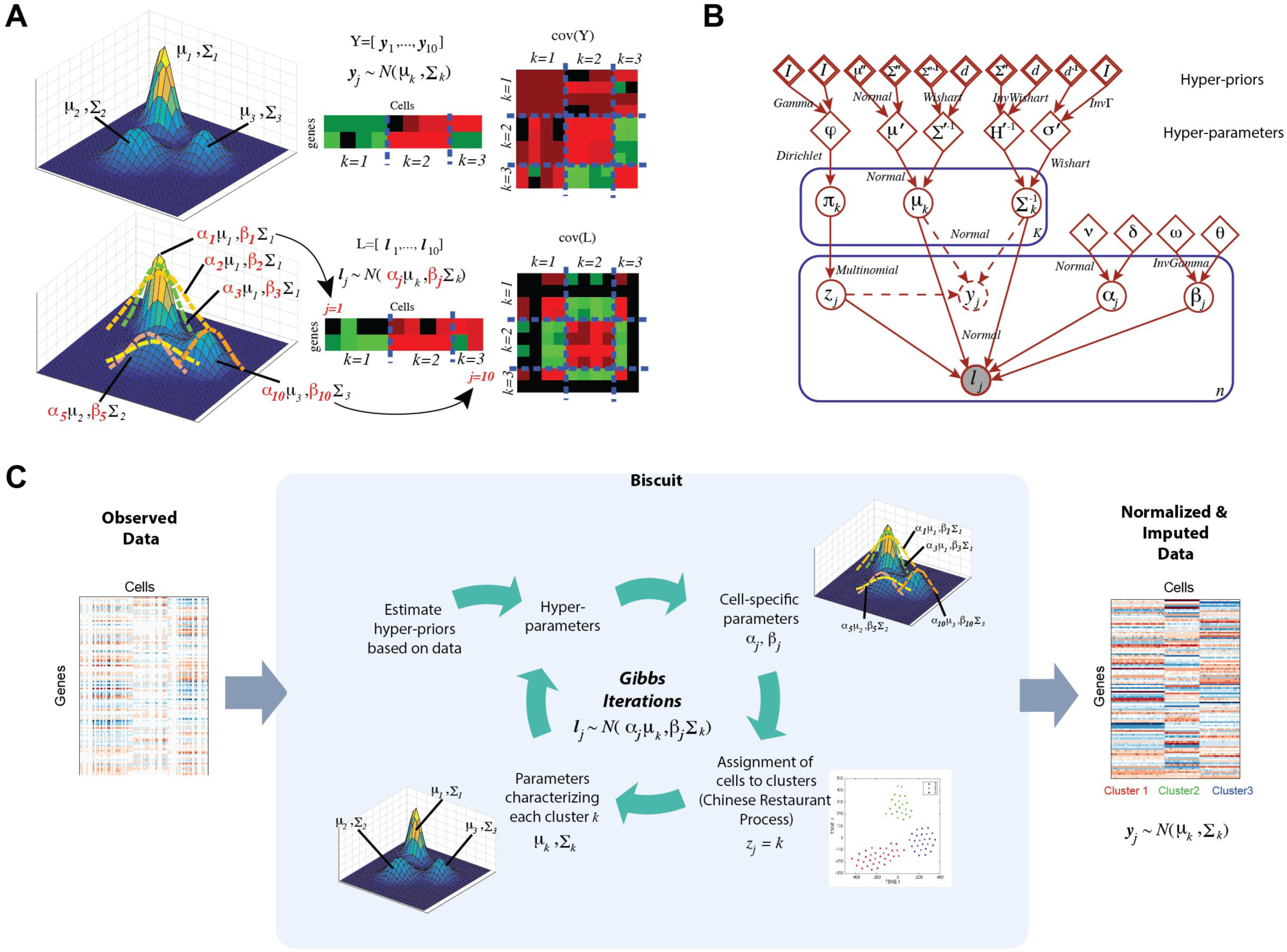
Details on Biscuit Model. (A) Stochastic data generative process for Biscuit illustrated with a toy example. **Top panel**: Left: shows 3 multivariate Gaussian densities with no technical variation. Middle: An ideal cell (yj) is simulated as a random draw from any of these 3 Gaussians. Right: The covariance matrix across 10 such randomly-drawn cells showing 3 block covariances across the diagonal corresponding to three clusters. **Bottom panel**: Left: shows 3 multivariate Gaussian densities with means and covariances scaled using (j, j) to handle cell-specific variations. Middle: A cell (lj) is simulated as a random draw from any of these 3 scaled Gaussians. Right: The covariance matrix across 10 such randomly-drawn cells showing loss of signal in the 3 block diagonal covariances. We assume the model for lj captures real single-cell measurements and the goal is to normalize data by converting it to follow the model for yj. (B). Finite state automata for Biscuit. The shaded circle denotes lj, which is observed gene expression for cell j, white circles show latent variables of interest, rectangles depict the number of replications at the bottom right corner, diamonds are hyper-parameters, and double diamonds are hyper-priors obtained empirically. Inference equations are obtained by inverting the date generative process. (C) Left panel: Input count matrix to Biscuit. Middle panel: Inference algorithm with Gibbs iterations are depicted where cell-specific (j, j) and cluster-specific (k, k) parameters are iteratively inferred leading to cell assignments to clusters. Right panel: Output from Biscuit, which is the normalized and imputed count matrix.

## STAR Methods

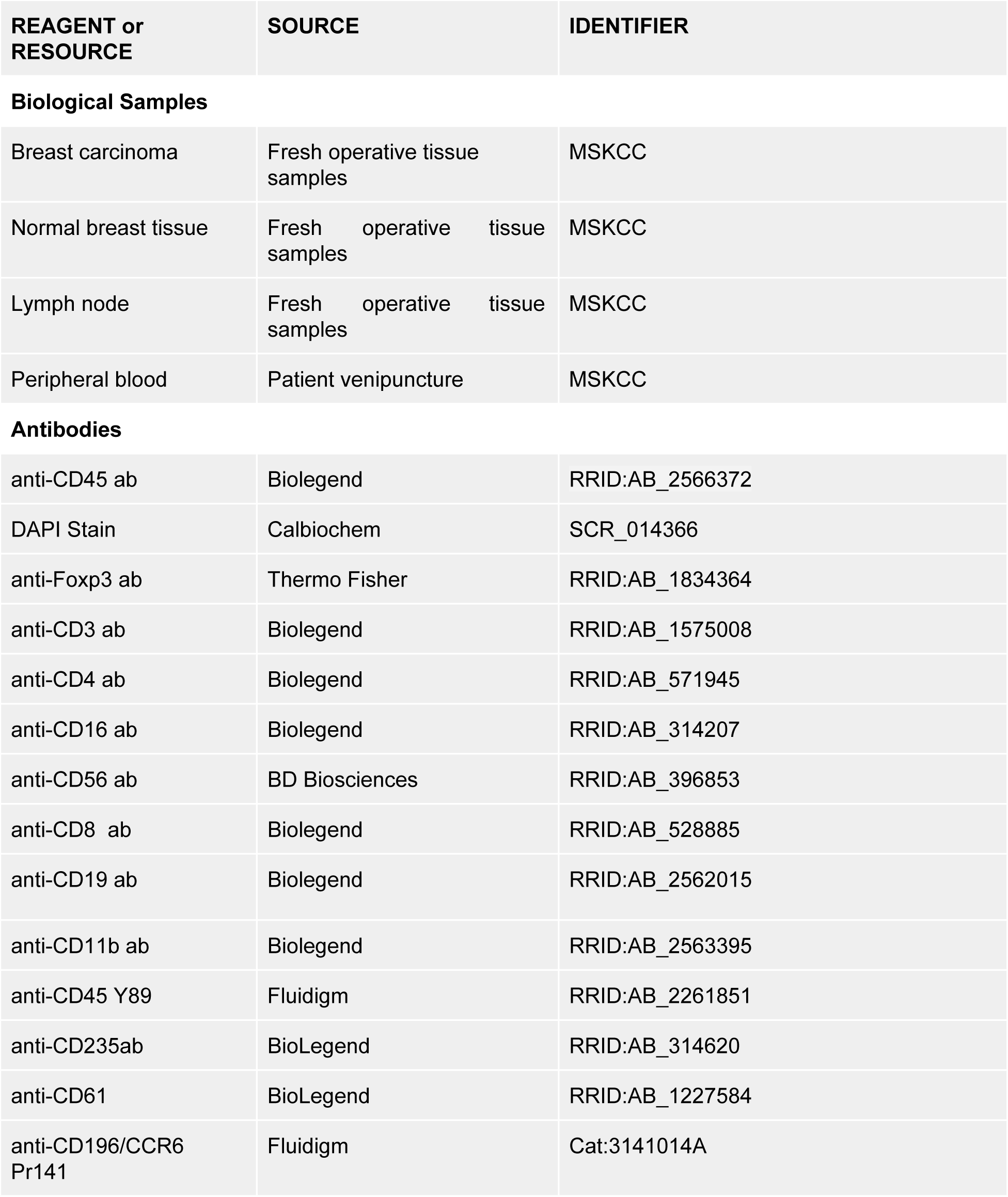
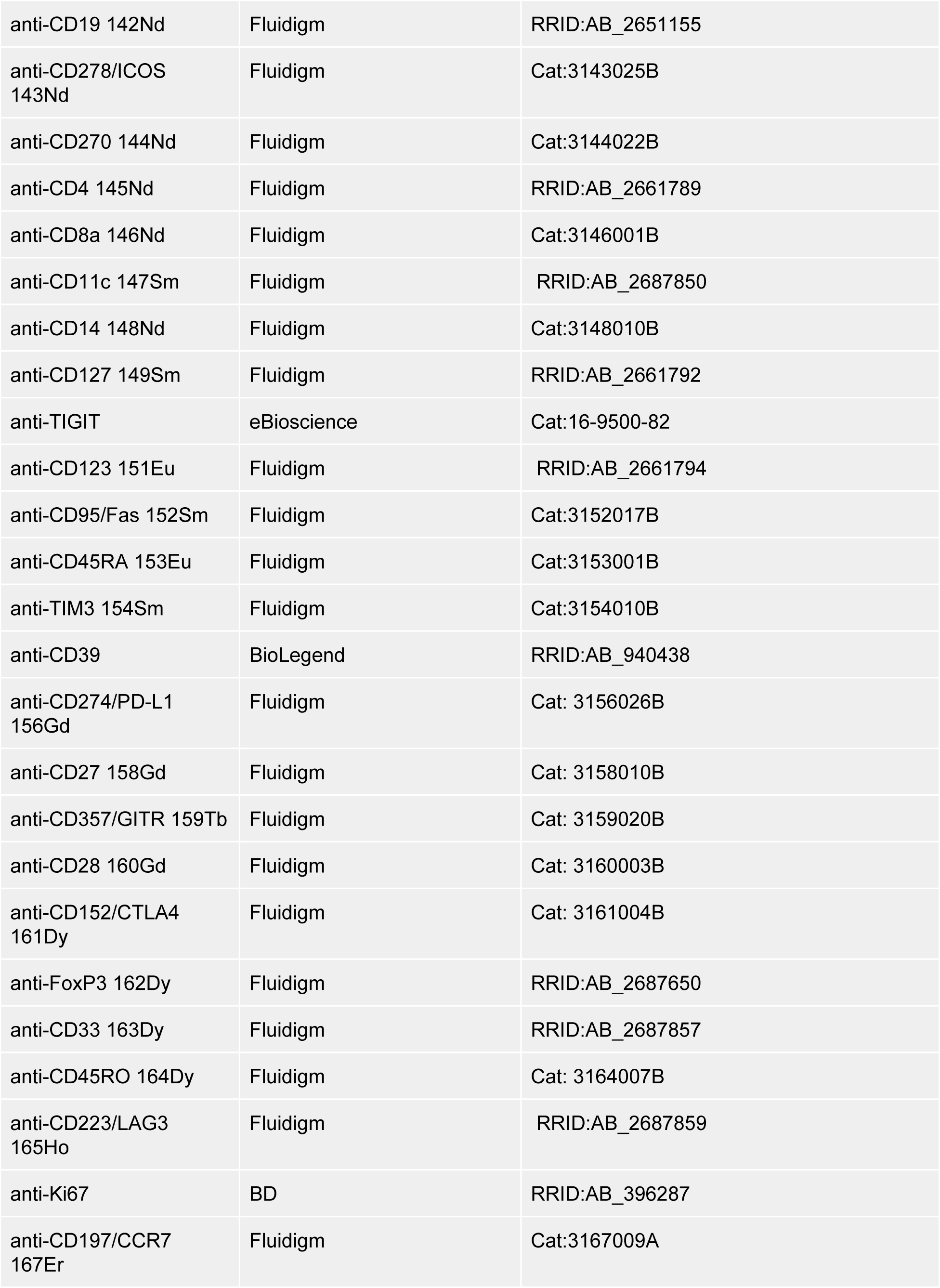
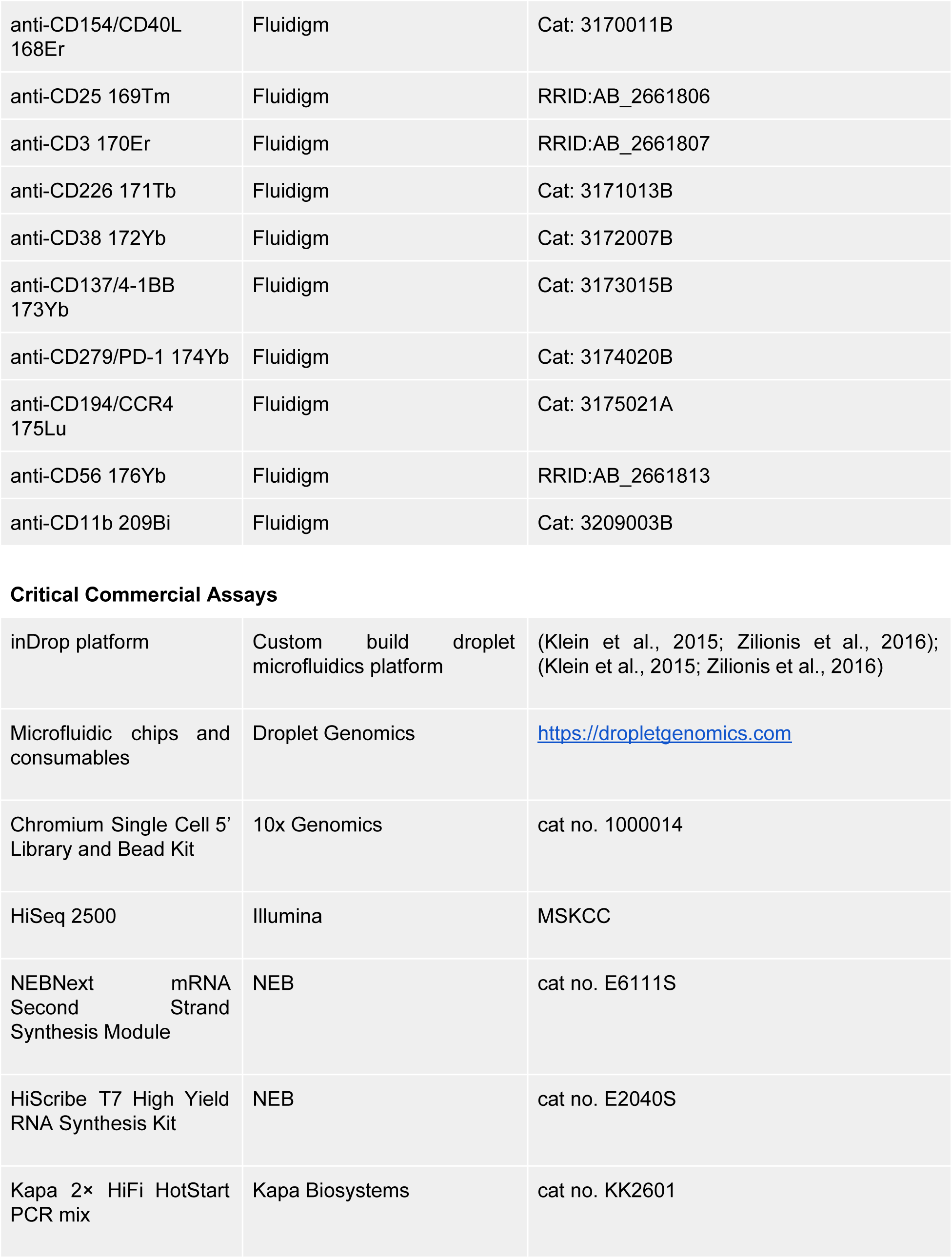
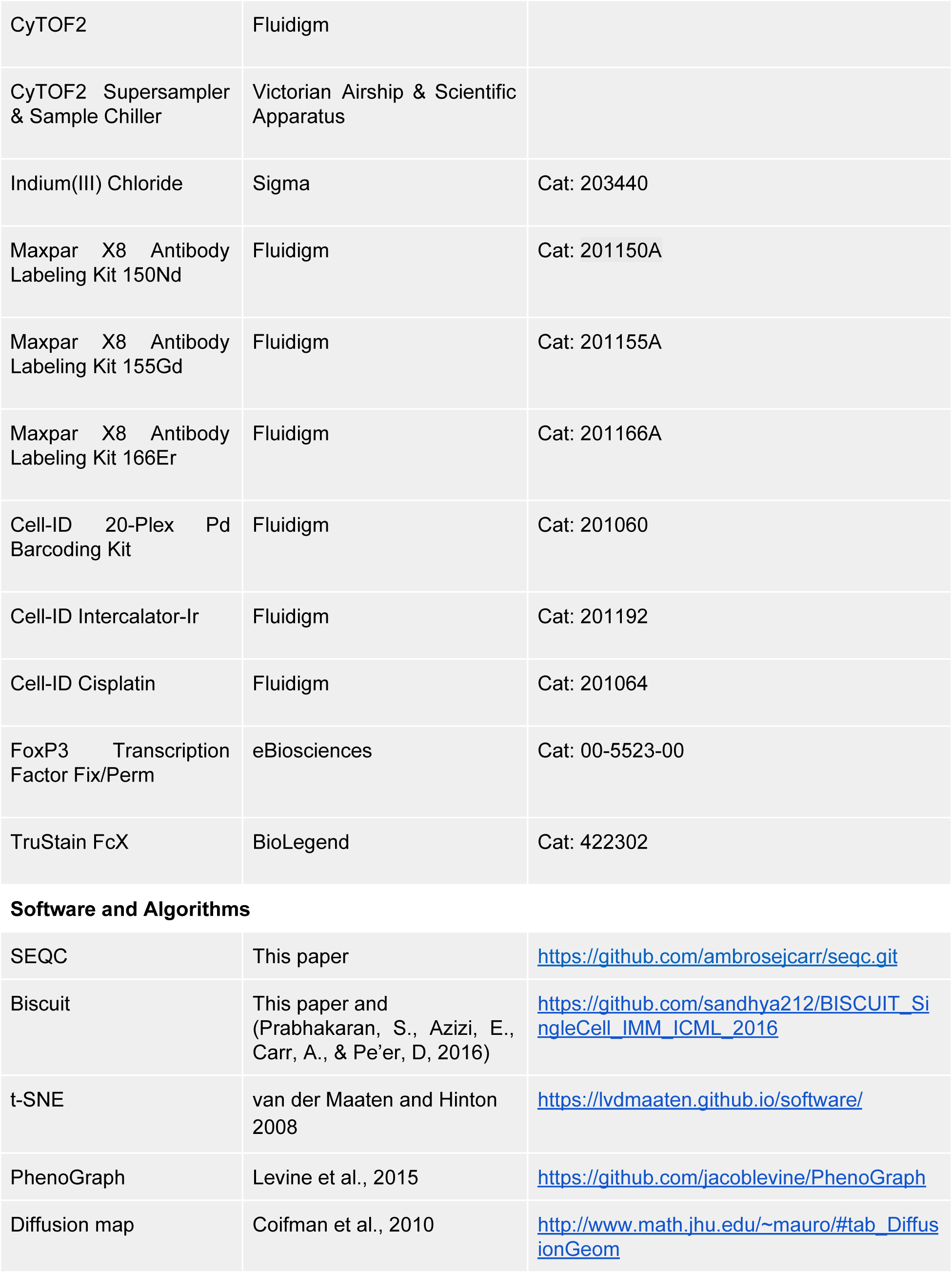
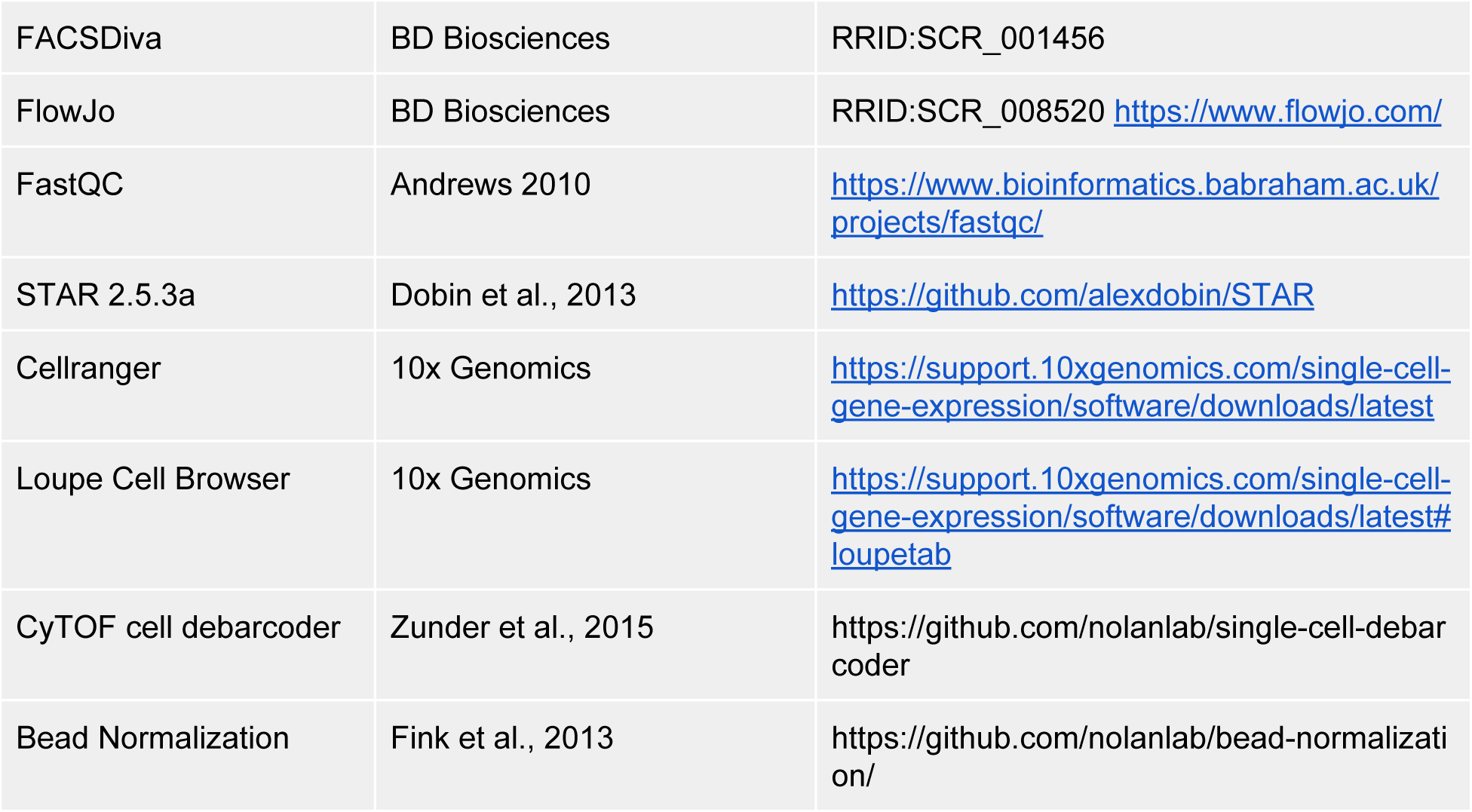
KEY RESOURCES TABLE

## CONTACT FOR REAGENT AND RESOURCE SHARING

Further information and requests for resources should be directed to and will be fulfilled by Lead Contacts Dana Pe’er and Alexander Rudensky.

## METHOD DETAILS

### Sample Collection

Tissues were collected from women undergoing surgery for primary breast cancer. Normal tissue was obtained from contralateral prophylactic mastectomies of the same cancer patients, and peripheral blood mononuclear cells (PBMCs) were obtained from patients prior to their surgical procedures. All samples were obtained after informed consent and approval from the Institutional Review Board (IRB) at Memorial Sloan Kettering Cancer Center. Clinical information and metadata for the samples are provided in Figure S1A, S6A. Samples were chosen from diverse subtypes, as we aimed to identify common changes in immune phenotypes in cancerous tissue as compared to normal tissue and blood or lymph node, rather than delineating differences across breast cancer subtypes.

CD45+ cells from tumor and normal tissues were isolated by mincing the freshly obtained surgical specimens into 1 mm cubic pieces and subsequent enzymatic digestion using Liberase TL (Sigma) for 20 min at 37°C. The digested tissues were then passed through a 100M filter and washed twice with PBS prior to surface staining. Immune cells were stained at 1*x*10^6 cells per ml for 20 min with anti-CD45 and DAPI for live-dead discrimination following Fc receptor blockade (BioLegend). Viable immune cells (CD45+DAPI-) were sorted on a FACSARIA sorter (BD Biosciences). Post-sort purity was routinely > 95% for the sorted populations.

Each sample was then divided into 10,000 cell aliquots and at least 2 technical replicates were processed through the complete experimental protocol (see below), with the exception that technical replicates were often processed on the same sequencing lane.

### Library Preparation for inDrop

We employed inDrop (Klein et al., 2015; Zilionis et al., 2016), a droplet-based single cell RNA sequencing technology. inDrop was selected over alternative technologies because it makes use of closely packed deformable hydrogel beads, ensuring that 75-90% of the input cells are paired with a unique barcode. This efficiency allowed deep sampling of many immune cells from individual patients, even in cases where immune infiltration was relatively low. As a result, we were able to compare triple-negative breast cancer (TNBC) samples with Her2+ and ER+ samples, which in some cases had as few as 50,000 tumor infiltrating immune cells.

Isolated, FACS-sorted CD45+ cells were suspended in ice-cold 1X PBS supplemented with 16% (v/v) Optiprep and 0.05% (w/v) BSA and encapsulated into 1.5 nL droplets together with custom-made DNA barcoding hydrogel beads and RT/lysis reagents. The microfluidics chip was operated at a throughput of ~30,000 cells per hour, and over 75% of cells entering microfluidics chips were co-encapsulated with one DNA barcoding hydrogel bead. The frequency of cell doublets (droplets having two cells) was low (~0.59%) due to highly diluted cell suspensions used for encapsulation, corresponding to approximately 1 cell for every 12^th^ droplet. In general, single-cell RNA-Seq library preparation was carried out following the protocol reported recently (Klein et al., 2015; Zilionis et al., 2016), with some modifications as described below.

### RNA-Seq library preparation for 10x Genomics single-cell 5’ and VDJ sequencing

The scRNA-Seq libraries were prepared following the protocol provided by the 10X genomics Chromium Single Cell Immune Profiling Solution. Briefly, approx. 12.000 FACS-sorted CD3+ immune cells (90-95% viability) were encapsulated into droplets at a concentration of 700 cells/uL, which results into expected mRNA barcoding of ~7000 single-cells with a multiplet rate 5.4%. After the RT step, droplets were broken and barcoded-cDNA was purified with DynaBeads, followed by 14-cycles of PCR-amplification (98ºC for 45s; [98ºC for 20s, 67ºC for 30s, 72ºC for 1 min] x 14; 72ºC for 1 min). The resulting amplified-cDNA was sufficient to construct 5` gene expression libraries and T cell receptor enriched libraries. The cDNA of single-cell transcriptomes (50 ng) was fragmented, double-size selected with SPRI beads (avg. size 450 bp) and sequenced on Illumina NextSeq platform (High Output V2 Kit, 150 cycles). The cDNA encoding the TCR library (10 ng) was amplified with 10-cycles of PCR (98ºC for 45s; [98ºC for 20s, 67ºC for 30s, 72ºC for 1 min] x 10; 72ºC for 1 min) followed by an additional 10-cycles of PCR (98ºC for 45s; [98ºC for 20s, 67ºC for 30s, 72ºC for 1 min] x 10; 72ºC for 1 min) using DNA primers provided in the kit. After library construction, VDJ region-enriched libraries were size selected with SPRI beads (avg. size 600 bp) and sequenced on an Illumina HiSeq 2500 instrument.

### Construction of new barcode sets for inDrop

GC content has a known impact on PCR efficiency (Mamedov et al., 2008): high or low fractions of G and C nucleotides reduce sequence amplification efficiency. We analyzed data produced with earlier version of DNA barcodes (Klein et al., 2015; Zilionis et al., 2016) and observed that barcodes with balanced GC content achieved higher molecule number (Figure M1A). We reasoned that balancing GC content across our barcodes would decrease variance across our libraries, thus increasing the average number of mRNA observed per cell. Further, we observed that the original barcode sequences had a minimum Hamming distance of 2. This is adequate to identify but not to correct single-base errors. We redesigned a library so that all barcodes had balanced GC content, with Hamming distance of >= 3, such that all single base errors are correctable, and with an average Hamming distance between pairs of barcodes of 13.3. This was done by performing a constrained optimization over barcodes of various lengths obtained from Edittag (Faircloth and Glenn, 2012). As a result, the vast majority of barcode errors are correctable and, as our results showed, the single-cell RNA-Seq libraries generated with new DNA barcoding hydrogel beads produced an overall increase in molecules/million sequencing reads of 5.3%.

The custom-made hydrogel beads carrying new DNA barcode sets were synthesized using the Agilent Bravo Automated Liquid Handling Platform following the previously described protocol (Klein et al., 2015; Zilionis et al., 2016). Before loading the DNA barcoding beads into the chip, they were washed twice in 1X SuperScript-III RT buffer and lysis reagent (1% (v/v) Igepal-CA630). In contrast to the approach in Zilionis et al., the Illumina PE Read 1 sequence was placed on the RT primer, thus the full-length primer sequence was as follows:

~~~
/5Acryd/PC/CGATGACG**TAATACGACTCACTATAG**GGATACCACCATGG<u>CTCTTTCCCTACACGACG</u> <u>CTCTTCCGATCT</u>[12345678901]GAGTGATTGCTTGTGACGCCTT[12345678]NNNNNNNNTTTT TTTTTTTTTTTTTTTV,
~~~

where 5Acryd is an acrydite moiety, PC is a photo-cleavable spacer, the letters in bold indicate T7 RNA promoter sequence, and underlined letters indicate the site for Illumina PE Read 1 Sequencing primer. The numbers indicate cell barcodes, which were specifically designed for this experiment to have 50% GC content and Hamming distance of >= 3 between each pair of barcode. Fluorescent in situ hybridization (FISH) analysis confirmed that hydrogel beads carried ~10^8 covalently-attached and photo-releasable barcoding DNA primers.

### Increasing the throughput

To increase the cell isolation throughput we used a cell barcoding chip (v2) (Droplet Genomics) and flow rates for cell suspension at 250 μl/hr, for RT/lysis mix at 250 μl/hr, and for barcoded hydrogel beads at 75 μl/hr. The flow rate for droplet stabilization oil was 550 μl/hr. Such flow parameters generated approximately 40,000 droplets an hour. After loading all components (cells, beads and RT/lysis reagents) into droplets, the final composition of a reaction under which cDNA synthesis was carried out was 155 mM KCl, 50 mM NaCl, 11 mM MgCl_2_, 135 mM Tris-HCl [pH 8.0], 0.5 mM KH_2_PO_4_, 0.85 mM Na_2_HPO_4_, 0.35 % (v/v) Igepal-CA630, 0.02 % (v/v) BSA, 4.4% (v/v) Optiprep, 2.4 mM DTT, 0.5 mM dNTPs, 1.3 U/ml RNAsIN Plus, and 11.4 U/ml SuperScript-III RT enzyme. After emulsion collection on ice the tube was exposed to 350 nm UV-light to photo-release DNA barcoding primers attached to the hydrogel beads. The RT reaction was initiated by transferring the tube to 65ºC for 1 min followed by a 1-hour incubation at 50ºC and 15 min at 75ºC. Post-RT droplets were chemically broken to release barcoded cDNA, which was then purified and amplified as described previously (Klein et al., 2015; Zilionis et al., 2016)). At the final step, libraries were amplified using trimmed PE Read 1 primer (PE1):

~~~
5’-AATGATACGGCGACCACCGAGATCTACACTCTTTCCCTACACGA
~~~

and indexing PE Read 2 primer (PE2):

~~~
5’-CAAGCAGAAGACGGCATACGAGAT[index]GTGACTGGAGTTCAGACGTGTGCTCTTCCGATCT,
~~~

where [index] encoded one of the following sequences: CGTGAT, ACATCG, GCCTAA, TGGTCA, CACTGT or ATTGGC).

Multiplexing of PCR libraries allowed for the pooling of different samples onto one lane of Illumina HiSeq2500 flow cell when desired.

### Sequencing and fastq quality control

Data were sequenced on Illumina HiSeq 2500 instruments using paired-end sequencing (PE1 54 bp and PE2 66 bp). Each replicate was sequenced on one half of a HiSeq lane, at an initial depth of approximately 100 million reads. scRNA-seq produces lower-complexity libraries than bulk sequencing techniques, which can infrequently lead to reduced base quality. Because each patient sample is precious, and because it is difficult to compare libraries with different average molecule counts, we verified the quality of each sequencing library with FastQC (Andrews, 2010), a software package that estimates the number of un-callable and low quality bases. Libraries that displayed significant (>25%) low quality bases were re-sequenced to maximize inter-sample comparability. See Supplementary Table S1 for sample sequencing depths.

### CyTOF sample preparation & data collection

CyTOF data was collected from three tumors (BC12-14) with the following metadata:

| Patient ID | Size | Metastases | Grade | ER | PR | Her2 | Postmenopausal | Age | Subtype | BRCA |
| --- | --- | --- | --- | --- | --- | --- | --- | --- | --- | --- |
| BC12 | 1.2 | n/a | 1 | 99 | 95 | - | + | 83 | Ductal | Unknown |
| BC13 | 0.9 | neg | 1 | 99 | 75 | - | - | 51 | Ductal | Unknown |
| BC14 | 2.7 | pos | 2 | 60 | 98 | - | - | 46 | Ductal | Unknown |

Custom conjugated antibodies were generated using MaxPar X8 antibody labeling kits (Fluidigm) exactly according to protocol. Tumor single-cell suspensions were quickly thawed and washed in PBS + FBS, then stained with 0.5mM Cisplatin for 5 min at room temperature. Reaction was quenched by adding Cell Staining Media (Fluidigm) containing EDTA. Cells were resuspended in FC-block (BioLegend) on ice for 20 minutes and washed. Cells were then stained with anti-CCR4 on ice for 30 minutes and washed in CSM. Samples were resuspended in Fix I Buffer (Fluidigm) for 10 minutes at room temp., then centrifuged and resuspended in Barcode Perm Buffer (Fluidigm). Appropriate palladium barcodes (Fluidigm) were added and incubated 30 minutes at room temp. Cells were washed 2x in CSM then combined. The data include tumors that were stained singly and tumors that were barcoded together with stock human PBMCs as carrier cells to increase cell yield during staining. A master mix for surface staining was prepared in CSM to yield final dilutions according to Supplementary Table S9. Cells were stained for 1 hour on ice, then washed in CSM, followed by MaxPar PBS (Fluidigm). Cells were fixed overnight in Fix/Perm buffer (eBiosciences) overnight at 4C, then washed in Perm Buffer (eBiosciences). A master mix for intracellular staining was prepared in Perm Buffer containing 2% fetal bovine serum to yield final dilutions according to Supplementary Table S9. Cells were stained for 1 hour at room temp. then washed twice in Perm buffer and once in MaxPar PBS. Cells were then resuspended in 1.6% paraformaldehyde in MaxPar PBS containing 0.5 μM Intercalator-Ir (Fluidigm) for at least 1 hour. Cells were then washed once in CSM, twice in MaxPar Water (Fluidigm) and resuspended in MaxPar water to a concentration of 1×10^6^ cells/mL. EQ 4 Element Calibration Beads (Fluidigm) were added to cells 1:10. Samples were acquired on a CyTOF2 (Fluidigm) equipped with a sample chiller and super-sampler (Victorian Airships) at a rate of < 400 events per second.

## QUANTIFICATION AND STATISTICAL ANALYSIS

### Data preprocessing: SEQC

#### Overview

At the time of data collection adequate analysis methods to construct count matrices from sequencing files for this data type were lacking. We therefore designed SEquence Quality Control (SEQC), a package that takes Illumina fastq or bcl files and generates a count matrix that is carefully filtered for errors and biases. The process is outlined in Figure S1B. We developed SEQC into a general purpose method to build a count matrix from single cell sequencing reads, able to process data from inDrop, drop-seq, 10X, and Mars-Seq2 technologies.

Briefly, SEQC begins by extracting the cell barcode and UMI from the forward read and storing these data in the header of the reverse read. This produces a single fastq file containing alignable sequence and all relevant metadata. The merged file is carefully filtered for cell barcode substitution errors, broken barcodes, and low-complexity polymers to eliminate errors early in the pipeline, saving analysis cost.

Filtered reads are aligned against the genome with STAR (Dobin et al., 2013), a high performance community-standard aligner. After alignment, minimal representations of sequencing reads are translated into an hdf5 read store object, where cell barcodes are represented in reduced 3-bit coding. Reads are annotated with a reduced set of exon and gene ids representing gene features—only the ones that are possible to detect with poly-A capture based droplet RNA sequencing—and SEQC attempts to resolve reads with multiple equal-scoring alignments.

In cases where genomic and transcriptomic alignments are present, the transcriptomic alignments are retained. Unique alignments from the previous step are corrected for errors using an enhancement of the method designed in Jaitin et al. (Jaitin et al., 2014), with an additional probability model to constrain the false positive rate. The error-reduced, uniquely-aligned data are grouped by cell, molecule, and gene annotation, and compressed into count matrices containing (1) reads and (2) molecules. This matrix is thresholded in a similar manner to what has been previously described (Macosko et al., 2015; Zheng et al., 2017).

Finally, SEQC outputs a series of QC metrics in an HTML archive that can be used to evaluate the quality of the library and the success of the run. SEQC is fully modular, and as such has been adapted to process drop-seq, 10x, and mars-seq data by switching de-multiplexing modules. In addition, it can be configured either to run on a local high-performance cluster, or can automatically initiate runs on Amazon Web Services compute platforms for those without access to local compute servers. The SEQC code is free and open-source, and can be found at https://github.com/ambrosejcarr/seqc.git, licensed under the MIT license.

#### Fastq Demultiplexing

The first stage of SEQC takes multiple fastq files containing genomic information and barcoding metadata spread across multiple sequencing files, and merges that information into a single fastq file using a “platform” class that comprises the locations of the cell barcodes and UMIs, the type of barcode and UMI correction to be run, and the number of T-nucleotides that are expected to be read from the capture primer. The merged fastq file contains genomic, alignable sequence in the sequence field, and has read metadata prepended to the name field, separated by colons. This step can be adjusted for novel sequencing approaches by adding a new platform class, often with only 10 lines of code. This allows the complete SEQC pipeline to be rapidly tested on iterations of new technologies.

For inDrop, which has variable-length cell barcodes, the description defines an additional method to localize the constant spacer sequence, which is flanked on both sides by the cell barcode (Figure M1B). The cell barcodes and molecular identifier are then extracted relative to the position of the spacer. Finally, we count the number of T-nucleotides that follow the UMI, where the poly-T spacer is supposed to be, and store this information for downstream filtering steps. The generated fastq file has the following format:

~~~
@<CELLBARCODE>:<UMI>:<#T>;GENOMIC READ NAME (read 2)
<GENOMIC SEQUENCE (read 2)>
+
<GENOMIC QUALITY (read 2)>
~~~

#### Substitution Error Rate Estimation

Two pieces of information *not* retained by the demultiplexing module are the cell barcode and UMI quality scores. Some pipelines, such as 10x Genomics’ Cell Ranger, posit that sequencing error is the major source of substitution mutations in 3’ sequencing data. Our inDrop data does not support this view of library construction. In inDrop, each read contains a 16-19 bp cell barcode selected from a whitelist of known barcodes. By examining barcodes for single base mutations, we estimated a positional, nucleotide-specific error rate for each sample (Table S1). E.g. to calculate the probability of a conversion from adenosine to cytosine, where *A* → *C* denotes this nucleotide conversion:

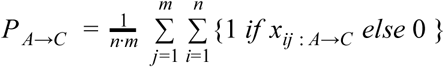

where *x*_*j*_ is a barcode, *j* ∈ {1, …, *m*} and each barcode has *n* bases.

The average observed per-barcode error rates are 4%, a number far in excess of the abundance reported by the Illumina sequencer, which can be reliably calculated from errors in phiX included in sequencing runs (mean error rate 0.2% ∓ 0.1%) (Manley et al., 2016); a 4% error rate is more in line with aggregate error rates of the enzymes used in the preparation of sequencing libraries.

To verify that quality scores do not predict error rates, we tested the correlation between the error state of the cell barcode (1 if the base contains an error and 0 otherwise) with Illumina quality scores. If quality were predictive of substitution errors, we would expect to observe strong negative correlations, suggesting that low quality implies high error probability. However, we observed no relationship (mean r^2^=0.04, max r^2^=0.06; ‘C’ errors) on either inDrop or 10x data.

In contrast, mutations to N bases produce the expected relationship, with base quality negatively correlating with → N substitutions (r^2^=-0.87). However, N base errors made up less than 1 / 100,000 of the observed errors in our experiment, and we conclude (1) that base quality is not meaningfully predictive of error rates, and (2) that most sequenced error is derived from upstream library construction steps.

While a 4% barcode error rate is higher than the error rate observed by other technologies, the use of linear amplification means that the errors we observe are non-cumulative: each transcript is generated from the original captured mRNA molecule, and as a result, 99.94% of observed barcodes have one or zero errors, all of which are correctable. This is in contrast to PCR-based amplification approaches which propagate errors that occur in early cycles, requiring more complex, graph-based correction methods, and larger Hamming buffers (see https://github.com/vals/umis).

#### Pre-alignment filtering

This module takes as input a raw merged fastq file and outputs a merged fastq file with corrected cell barcodes and full length UMI sequences (that may still contain substitution errors).

#### Cell barcode correction

Cell barcode errors in inDrop are easy to detect by design: we have a whitelist of 147,456 barcodes, each with Hamming distance >= 3. Thus, any single base substitution error is resolved by creating a lookup table for all barcodes and all single substitutions. If found in the table, the barcode is corrected. If not, it is discarded. As estimated above, the probability of a cell barcode containing an error is ~2%, and thus the expected rate of barcodes accruing 2 errors in a barcode is 1 / 2500. A 2-error lookup table has a very large memory footprint and would significantly increase computational cost of processing each experiment. Alternative algorithms have greater complexity and would increase run time. Thus, we accept this low rate of loss and proceed to correct single base errors, recovering approximately 2% additional data for each sample.

#### UMI validation

In contrast to cell barcodes, UMIs are random, and correction cannot proceed by the same strategy, so we devote a section later in the pipeline to the detection of UMI errors after the gene and mapping position of a fragment are identified.

Another source of error in scRNA-seq experiments, including inDrop, cel-seq, mars-seq, and likely drop-seq and 10x genomics, is the fracturing and random-priming of capture primers (Figure M1C) (Jaitin et al., 2014). We often observe cell-barcode prefixes followed by randomers. When fragmentation occurs at the cell barcode level, we can remove the fragments using the whitelist approach above. To remove barcodes that break in the UMI, we determined that we would sequence 5 bases into the poly-T tail of the primer, which we expect to be all T-nucleotides. By excluding reads with more than 1 non-T nucleotide, we are able to exclude most broken UMIs.

In aggregate, the filters in this section remove an average of 36% of reads (sd = 9.3%), depleting the count matrix of spurious molecules (see Table S1 for detailed values). These values are consistent with the results of running SEQC on drop-seq or MARS-seq datasets (data not shown).

### Annotation Construction

Because the genome annotation is designed to be broadly applicable across sequencing modalities, it contains many features that are theoretically undetectable by inDrop and other 3’ sequencing technologies. To address this, we constructed a custom annotation by starting with the current GENCODE genome and GTF file and removing all feature annotations that are not theoretically detectable by inDrop.

Two characteristics of inDrop limit its ability to capture certain gene biotypes. First, it employs poly-A capture, and thus will not detect non-polyadenylated transcripts. Second, it uses SPRIselect beads at several stages to deplete primers from reaction media. These beads carry out size selection, preferentially depleting primers but also small RNA species such as snoRNA, miRNA, and snRNA. Thus, libraries are expected to contain only transcribed, polyadenylated RNA of length > ~200 nt. Examining gene biotypes, this meant retaining protein coding and lncRNA biotypes, and excluding others.

To determine the impact of this change of reference on our data, we aligned the same single-cell immune dataset against the full reference and the reduced reference described above. We constructed a pseudo-bulk dataset for each reference by summing the molecules across all cells, producing an expression vector that contained a single value for each gene. We hypothesized that the reduction in reference features would result in a concentration of alignments in biologically relevant genes by depleting non-specific features, and that there would be many drop-out events where genes would be detected in the complete reference, but not the subset.

This is exactly what was observed (Figure M1D). The overall r2 value between the references is 0.94, with 93% of genes holding the exact same values in both reference alignments. In addition, information is concentrated in 35% fewer features, despite losing only 8% of the total molecules. There is also a large drop-out contingent present only when aligned against the complete reference. Gene ontology enrichment against this reference revealed high-level biologically agnostic enrichments, such as “protein coding,” “translation,” and other enrichments, which suggest a random sampling of high-expression genes.

Surprisingly, there was also a contingent of genes present only in the reduced alignment. These genes were highly enriched for immunological pathways, including JAK/STAT signaling, cytokine production, cytokine receptors, and immune growth factors, and further included critical immune genes such as IL3RA (Figure M1D). This suggests that they are likely to represent true annotations for genes in this dataset, and that reducing the annotation produces a gain in specificity. We reasoned that these genes were uncovered in the reduced annotation because there are features in the complete set, such as pseudogenes, which have high homology to transcribed genes. Including these annotations, which should not be detectable, produces illogical multi-alignment to multiple genetic locations. When such multi-alignment cannot be resolved, most pipelines (including this one) exclude those multi-aligned reads, losing valuable signal. Given these results, we believe that the 8% molecule loss is the result of correctly discarding low-complexity alignments that were spuriously assigned a low-quality transcriptomic feature.

We note that Cell Ranger, the most commonly used 10x pipeline, carries out an extreme version of this redesign: it removes any gene that is not protein coding. We believe that this is too harsh: it excludes numerous transcribed pseudogenes and lincRNA which have been previously shown to be expressed, have biological functionality, and be detectable in scRNA-seq.

The extreme case of annotation redesign is to exclusively align to expected features. Several methods exist to accomplish this, including Kallisto (Bray et al., 2016) and Tophat2 (Kim et al., 2013). However, 3’ scRNA-seq data typically contains between 10-30% genomic contamination, as identified by intergenic alignments. When we aligned directly against the transcriptome, we found that approximately 1% of intergenic reads were mistakenly aligned to exonic locations despite having higher alignment scores to genetic regions (data not shown). As a result, we align to the genome, considering only detectable features, and prefer transcriptomic alignments in cases where there are equivalent genomic and transcriptomic alignments, but remove reads that score highest against the genome.

### Alignment

The merged, single-ended fastq files are aligned to hg38 with STAR using the annotation file as described. We selected STAR because it is a fast, highly parallel, cloud-scalable aligner that benchmarks well against existing aligners (Illicic 2016). We note that STAR automatically trims bases as necessary to find alignments, and as such no pre-trimming of reads based on quality is carried out. Alignment parameters used are as follows: –outFilterType BySJout, –outFilterMultimapNmax 100, –limitOutSJcollapsed 2000000 –alignSJDBoverhangMin 8, –outFilterMismatchNoverLmax 0.04, –alignIntronMin 20, –alignIntronMax 1000000, –readFilesIn fastqrecords, –outSAMprimaryFlag AllBestScore, –outSAMtype BAM Unsorted

This module thus takes as input a fastq file and produces a bam file containing up to 20 multiple alignments per input fastq record, with all unaligned reads contained in the same file. This format is useful for archival purposes, as it can be used to reconstruct the original merged fastq file without data loss.

### Multi-Alignment Correction

Alignment algorithms aim to identify the unique portion of the genome that was transcribed to generate the read that is being aligned. In some cases, this unique source cannot be identified, and in these cases multiple possible sources are reported. These are commonly termed “multi-alignments”, and because 3’ ends of genes have higher homology than other parts of the genome, multi-alignments are more common in 3’ sequencing data than in approaches that cover the full-transcriptome, such as Smart-seq2. Despite the increased frequency, most 3’ pipelines discard multi-alignments and deal exclusively with unique genes. This module is designed to resolve all multiple alignments, producing an output that contains resolved (now unique) alignments.

There are several existing approaches to resolving multi-alignments, of which transcriptomic pseudo-alignment, such as that done by Salmon (Patro et al., 2017) and Kallisto (Bray et al., 2016), and EM approaches, such as RSEM (Li and Dewey, 2011), are the most common. However, both methods have the effect of decreasing signal-to-noise ratios for inDrop sequencing data. For RSEM, low-coverage 3’ sequencing data contains considerable uncertainty, which RSEM is designed to pass into the count matrix. This uncertainty is normally removed by UMI-aware count based methods, although it incurs some data loss. Kallisto and Salmon, in contrast, both pseudo-align and resolve multi-alignments, but only against the transcriptome. This causes alignments from contamination that is of genomic origin to be pseudo-aligned to transcriptomic positions, producing inflated and spurious alignments for low-homology genes.

Of the high-throughput droplet-based approaches, inDrop has a unique combination of linear amplification and UMIs, which produces high fragment coverage per UMI. Although individual alignments are often ambiguously aligned to more than one location, it is often possible to look at the set of fragments assigned to an UMI and identify a unique gene that is compatible with all the observed fragments. Here we implement an efficient method to find the unique genes that generate each fragment set. When a fragment set cannot be attributed to a specific gene, it is discarded.

Starting with all reads attributed to a cell, we begin by grouping reads according to their UMI, producing “fragment sets” *S*. Typically, these fragment sets represent trivial problems, such as *s*_1_ = {*A*, *A*, *AB*}, a set with two unique alignments to gene A and a third ambiguous alignment to genes A and B. In this case all three observations support the gene A model, while only one observation supports the gene B model.

In cases of UMI collisions, where two mRNA molecules were captured by different primers that happen to share the same UMI sequence, this can lead to problems wherein reads from these merged fragment sets are mistakenly discarded as multi-aligning. However, because the probability of two genes sharing significant homology is low, it is usually possible to recover these molecules by first separating fragment sets into disjoint sets. For example, if a fragment set

*s*_2_ = {*A*, *AB*, *B*, *B*, *C*, *CD*, *ABC*, *E*, *EF*, *EF*},

it is broken into two disjoint groups:

*s*_2_ = *s*_3_ ⋃ *s*_4_; *s*_3_ ⋂ *s*_4_ = ∅ where: *s*_3_ = {*A*, *AB*, *B*, *B*, *C*, *CD*, *ABC*} and *s*_4_ = {*E*, *EF*, *EF*}.

This is biologically reasonable, as molecule collisions are the only way to reasonably obtain a group of molecules that covers two non-overlapping gene annotations.

To calculate disjoint sets efficiently, we utilize a Union-Find data structure (Aho and Ullman, 1983), which finds disjoint sets in *O*(*log*(*n*)) time. Pseudo-code is as follows:

~~~
summarize alignments as data = {cell=c, umi=u, gene=g}
sort data by c > u > g
for fragment_set in data[{c, u}]:
    for disjoint_set in union_find(fragment_set):
         find number_shared_genes in disjoint_set
         if number_shared_genes == 1:
                resolve disjoint_set
~~~

We note that this allows us to more accurately identify the alignment rates for each gene, build better error models for barcode correction, and recover cases where reads align multiply to the same gene. More critically, it gives us the ability to recover fragments that would otherwise not be resolvable due to sequence homology, and these improved fragment counts per molecules act as significant predictors of molecule likelihood and UMI quality. We note that a similar strategy has since been published by (Klein et al., 2015) and a comparable logic underlies the concept of transcript compatibility in Kallisto (Bray et al., 2016).

We had previously created a model wherein disjoint sets with more than 1 common gene could also be disambiguated by calculating the probability of gene-gene multi-alignments from their homology, by comparing gene sequences using a Suffix Array built from the final 1000 bases of each gene. With this strategy, we could estimate the relative probability that genes were generated from each potential candidate molecule shared across all reads in the fragment set. However, the relative rarity of such events (<1% of data) combined with the additional run-time complexity of this method caused us to omit it from the production version of SEQC. This module typically resolves approximately 1M reads per hiseq lane. The result of this module is a bam or h5 file containing only unique alignments to gene features.

### Molecular Identifier Correction

Errors in molecular identifiers are well-known to introduce noise in sequencing experiments (Jaitin et al., 2014), since undetected errors induce spurious increases in molecule counts. This module takes an hdf5 read store, identifies errors in UMIs, and replaces them with their corrected value. The most common approach, published in (Jaitin et al., 2014) for MARS-seq, does a very good job of detecting and removing molecule errors in inDrop (due to similarity in the Cell-seq protocol used in both technologies). This approach deletes any UMI for which a higher-abundance donor UMIs can be identified that (1) lies within a single base error (2) has higher count (3) and contain all observed alignment positions of the recipient RNA. This results in removal of approximately 20% of observed UMIs. However, we observed that this model can be overly stringent, correcting UMIs when the donor molecule has as few as one read count higher than the recipient.

We apply a modified version of the (Jaitin et al., 2014) approach, where we replace errors with corrected barcodes instead of deleting them, and where we only eliminate errors when we have adequate statistical evidence. To accomplish this, we utilize the spacer and cell-barcode whitelist to empirically estimate a per-base error UMI error rate of approximately 0.2% per base, e.g. to calculate the probability of a conversion from adenosine to cytosine, where *A* → *C* denotes this nucleotide conversion:

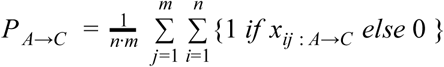

where *x*_*j*_ is a barcode, *j* ∈ {1, …, *m*} and each barcode has *n* bases.

To calculate the probability a target read was generated in error from a specific donor molecule, we calculate the product of the errors that could potentially convert a donor into the observed molecule. To convert, for example, ACGTACGT into TTGTACGT, having one *A* → *T* and one *C* → *T* conversion:

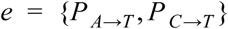

The probability of the above conversion is

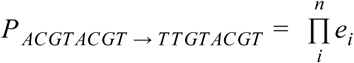

Because there are multiple potential donors for each molecule, we calculate the conversion probability for each molecule. Assuming errors are randomly distributed, they can be modeled by a Poisson process, and Poisson rate term can be estimated from the data:

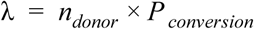

where *n*_*donor*_ is the number of observations (reads) attributed to the donor molecule in the data. Since the sum of multiple Poisson processes is itself Poisson, the rate of conversion from each donor can be combined into a single rate λ^*agg*^ for each target molecule. The set of conversions *s* that we consider for each target molecule are all conversions that can occur with two or fewer nucleotide substitutions, in other words, all molecules within a Hamming distance *D*_*h*_ ≤ 2, where *D*_*h*_ is a matrix of pairwise Hamming distances between barcodes.

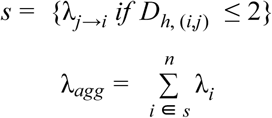

Finally, given the probability of a molecule being observed via the substitution errors that are corrected by the Jaitin method, we can calculate the probability that *n* observations of a specific molecule *x* were generated via the Poisson process with rate λ_*agg*_:

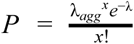

Cases that are very unlikely (p < 0.05) are *not* corrected. For inDrop experiments, this results in a recovery of an additional 3-5% of molecules in the data that are otherwise error-corrected. We note that this model is not applicable to all data; it is useful in this instance because we have relatively high coverage (10 reads / molecule) that allows us to evaluate our confidence in molecule observations. For lower-coverage data, it may be appropriate to err towards removing molecules instead of retaining them.

### Raw Digital Expression Matrix Construction

To create a digital expression matrix, the uniquely-aligned, error-corrected hdf5 read store is de-duplicated by counting unique groups of reads with the same UMI, cell barcode, and gene annotation. A single molecule then replaces each set, and those molecules are summed to create a cells x genes matrix. scRNA-seq count matrices are often over 95% sparse, and thus are stored in matrix market format and operated on as coordinate sparse or compressed sparse row matrices. We call these count matrices “raw” count matrices because they contain all barcodes observed in an experiment.

### Cell Selection and Filtering

#### Size Selection

Barcoding beads are loaded into inDrop at higher rates than cells in order to ensure that a high fraction of cells are encapsulated with exactly one bead. As a result, the raw count matrix contains a mixture of cell barcodes that were encapsulated with cells and cell barcodes that were encapsulated alone, but may nevertheless capture some ambient mRNA molecules that float in solution due to premature lysis or cell death in the cell solution. We separate these by finding the saddle point in the distribution of total molecule counts per barcode and excluding the mode with lower mean. In practice, we accomplish this by constructing the empirical cumulative density function of cell sizes and finding the minimum of the second derivative (Figure M1E) of the distribution. For typical inDrop runs, this results in the elimination of over 95% of the cell barcodes, but retains as many as 95% of the molecules.

#### Coverage Selection

Molecule size alone is not adequate to remove all barcodes that were not paired with real cells. Some barcodes appear to aggregate higher numbers of errors, and as such we often see a bimodal distribution of molecule coverage: a higher mode that represents real cells, and a smaller mode that represents aggregated errors (Figure M1F). We remove the low-count density by fitting 2-component and 1-component Gaussian mixture models to each axis and comparing their relative fits using the Bayesian information criterion. When the 2-component model’s difference in likelihood is at least 5% larger than the 1-component model, we exclude the densities with the smaller mean (Figure M1G).

#### Filtration of dead or dying cells

We score cells for mitochondrial RNA content, which is widely used as a proxy for cell death in scRNA-seq. We observe that a small fraction of cells contain a higher abundance of molecules annotated by this signature, as much as 20−95% of their RNA. Since inDrop does not lyse mitochondria, we reason that these are likely to be cells dying due to stress imposed on them by the inDrop procedure or prior sorting, and remove them from further analysis. This filter may be turned off for studies where apoptosis is a relevant phenotype.

#### Low-complexity cell filtration

Finally, we regress the number of genes detected per cell against the number of molecules contained in that cell. We observe that there are sometimes cells whose residuals are significantly negative, indicating a cell which detects many fewer genes than would be expected given its number of molecules. We exclude these cells whose residual genes per cell are more than 3 standard deviations below the mean (Figure M1H).

### Information Storage & Run Time

scRNA-seq generates large volumes of data whose storage can be costly and onerous. We thus store only aligned, barcode-tagged bam files which losslessly retain all information from the original multiplex fastq files in small storage space. SEQC supports reprocessing of these files, and backwards conversion into fastq files, if users desire the ability to process their data on other platforms or reprocess with updated versions of SEQC. Additional metadata files take up nominal space, and generated count matrices are stored in matrix market sparse format in light of the sparsity of the data. SEQC requires approximately 8 hours to run on a standard 32 GB / 16 core Amazon c4.4xlarge, and costs $5.84 on on-demand or $0.88 on preemptible (spot) instances to process an inDrop, drop-seq or 10x genomics experiment. In contrast, the 10x genomics commercial pipeline Cell Ranger costs approximately $20, in large part due to high RAM requirements.

### Data Quality Analysis of Breast Leukocytes

We applied SEQC to each of the 14 inDrop samples and 61 replicates in our data. Each sample had a minimum of 2 replicates (Table S1). Samples were sequenced such that each cell was covered by an average of 22,000 reads. Cells contained on average 15 reads per molecule, and cell saturation was 91% across all samples and replicates. On average, 20% of cells were excluded due to high mitochondrial content, a proxy for cell death and stress. Samples obtained from tissue requiring dissociation had significantly higher mitochondrial transcripts (25%) than those obtained from blood (13%) (t=2.42, p=0.018). Small numbers of low-complexity cells displaying fewer than expected genes for their molecule count were detected and removed, removing 1.2 ∓ 2.3% of total molecules. A complete summary of read abundances, genes detected, and number of cells can be found in Table S6. Having excluded non-viable cells from downstream analysis, we shifted to examining within-sample consistency across technical replicates.

### Library Consistency and Quality Control

The inDrop encapsulation procedure runs 10,000 cell aliquots in series, leaving open the possibility of batch-to-batch variation within technical replicates of patient samples. To determine the magnitude of batch-to-batch variation, we compared the variation within patient replicates to between-patient comparison. Each sample was collapsed into pseudo-bulk by summing over the cells of the digital expression matrix. We determined intra-patient consistency by determining the average pairwise Pearson correlation between pairs of samples, and compared that against the inter-patient correlations. After excluding one aberrant sample with a sample-sample correlation of 0.6, we observed Pearson correlations with a minimum of r2=0.92, a mean of 0.97, and a standard deviation of 0.02 (full pairwise correlation matrix in Figure M1I). This is in contrast to the complete pairwise correlation matrix, where the average correlation between patients is 0.72. This suggests that patient-to-patient variation is primarily biological, and that we have low technical variation between inDrop runs. These comparisons are generated automatically by SEQC, and serve as an internal control for batch variation.

### Individual Sample Normalization and Clustering

To characterize the immune cells extracted from patients in this study, we began by analyzing samples independently to identify their cellular composition and cell type abundances (Figures 1C, S1C). Cells were first normalized by median library size.

The normalized data was then decomposed using randomized principal component analysis (Halko et al., 2009). We selected the number of principal components to retain using the knee point (Valle et al., 1999). This resulted in 6-10 principal components per sample. The dimension-reduced PCA projection was used as the input to PhenoGraph (Levine et al., 2015), which was used to cluster the data with default parameters (k=30 nearest neighbors). The same principal components were used to generate tSNE projections (Maaten and Hinton, 2008), which were generated with barnes-hut tSNE, implemented in the bhtsne package https://github.com/lvdmaaten/bhtsne (Figure 1C, S1C).

### Cluster Cell Type Annotation

As reference data, to facilitate the interpretation and labeling of clusters derived from PhenoGraph and Biscuit, we collected previously generated bulk gene expression profiles of sorted cells from a number of sources. To label our immune clusters we collected profiles from several published datasets on sorted immune populations (Novershtern et al., 2011) (Jeffrey et al., 2006), which provided 37 and 32 sorted populations, respectively. In addition, samples from the ENCODE consortium were added as negative controls to identify and subsequently remove contaminating stroma and tumor cells which may have infiltrated the sample due to low-level CD45 expression or auto-fluorescence. To determine the gross cell types of the clustered immune cells extracted in this experiment, we examined the correspondence between cluster centroids for PhenoGraph clusters or mean parameters for Biscuit clusters and the collected cohort of bulk profiles.

Bulk samples were library-size normalized and pre-processed by mean centering, and in the case of microarray data from (Novershtern et al., 2011) (Jeffrey et al., 2006), the data was also scaled by standard deviations. In the case of PhenoGraph clusters (Figures 1C, S1C), cells were library size normalized, clusters were mean centered, and low-expression genes with an average of fewer than 1 count per cell were removed to ensure the correlations were based on the highly expressed genes in each cell type. Cluster centroids were then correlated with bulk profiles. In the case of Biscuit clusters (Figures 2E,F), mean parameters for each cluster were correlated with bulk profiles.

Each of the bulk profiles was marked as having derived from one of several major cell types: B-cells; T-cells (naive, central memory, cytotoxic, Treg); Monocytic cells (monocytes, dendritic cells, macrophages); Mast cells; Neutrophils; or NK-cells. The highest-scoring bulk profile for each centroid was used to categorize each cluster by its type, and types were split for downstream analysis.

Cells were also typed by examining expression of known marker genes. In this analysis, cells were scored as detecting a marker gene if the cell contained a non-zero molecule count for that gene. Each cell was corrected for its detection rate (the fraction of total genes detected in that cell) and the marker detection rate was then averaged across cells of a cluster. Markers used to type cells included NCAM1, NCR1, NKG2 (NK-cells), GNLY, PFN1, GZMA, GZMB, GMZM, GZMH (cytotoxic T, NK), FOXP3, CTLA4, TIGIT, TNFRSF4, LAG3, PDCD1 (Exhausted T-cell, T-regulatory Cell), CD8, CD3, CD4 (T-cells), IL7R (Naive T-cells), CD19 (B-cells), ENPP3, KIT (Mast cells), IL3RA, LILRA4 (plasmacytoid DC), HLA-DR, FCGR3A, CD68, ANPEP, ITGAX, CD14, ITGAM, CD33 (Monocytic Lineage). For all retained clusters, the two typing methods were in agreement (Figure 2H).

Twelve populations did not correlate highly with any of the immune populations sorted and sequenced in Dmap and Garvan, and were unique among the Biscuit clusters. However, when the same analysis was performed with ENCODE data that included non-immune populations, we found that these clusters, which are marked in Table S2 (clusters 7, 10, 13, 14, 17, 18, 26, 32, 33, 34, 52, and 79) correlated much more highly with various bulk-sorted epithelial, stromal, and endothelial cell populations. Further, manually examining their Biscuit-identified differentially expressed genes, provided in Table S3, led to an identical conclusion. They were thus labelled as non-immune and excluded from the present study.

### Gene Signature Summarization

While attempting to interpret biological signals that were observed in the studied immune cells, we realized that it was important to consider several facets of gene signature enrichment: (1) the classically studied mean value of the signature across cells in the cluster, but also (2) the marginal distribution of cell loadings across the signature, and (3) the relative contribution of each gene. Therefore, for plots of signature expression by patient we began by constructing a barplot of the counts for each gene in the signature, corrected for cellular observation rate (the total number of genes observed with molecule count > 1). This displays the contribution of each gene to the signature (top panel in Figure 1E, 2B, and S1E-G). The normalized values for each signature, per cell, are then summarized as a box plot to display the variation of cells in each patient (left panels). Finally, the cluster median of each gene is taken per patient, and the cluster medians are z-scored across patients. The z-scored values are plotted as a heatmap (center-right panel in Figure 1E, S1E-G and 2B), facilitating a comparison of signatures across patients.

The full lists of compiled gene signatures used throughout this paper can be found in Table S4 and may serve as a valuable resource for additional investigations. To create these lists we broadly surveyed the extant literature and manually curated consensus lists of genes to be included. The relevant literature surveyed to form these signatures includes the following:

For the M1 and M2 macrophage polarization signatures we merged gene lists from (Sica and Mantovani, 2012); (Biswas and Mantovani, 2010); (Bronte et al., 2016) (Ugel et al., 2015) (Gabrilovich, 2017). For other myeloid-specific signatures we used (Villani et al., 2017) (pDCs, AXL/SIGLEC6 DCs, CD141/CLEC9A DCs, CD11C_A DCs, CD1C-/CD141-DCs, CD1C_B DCs, New Monocytes 1, New Monocytes 2, CD14+CD16-Classical Monocytes, and CD14+CD16+ Non-Classical Monocytes); and (Gesta et al., 2007), (Perera et al., 2006), (Farmer, 2006; Lefterova and Lazar, 2009) (Lipid Mediators).

For T Cell-specific signatures we used (Wherry and Kurachi, 2015), (Wherry, 2011), and (Schietinger et al., 2012) (Exhaustion and Anergy); (Glimcher et al., 2004) (Cytolytic Effector Pathway); and (Smith-Garvin et al., 2009), (Chtanova et al., 2005), and (Adam Best et al., 2013) (T Cell Activation).

For gene signatures used across cell types we used (Mantovani et al., 2008), and (Grivennikov et al., 2010) (Pro and Anti-Inflammatory); (Platanias, 2005) (Type I and II Interferon Responses); (Ho et al., 2015) (glucose deprivation); (Benita et al., 2009; Makino et al., 2003) (Hypoxia/HIF Regulated); (Moreno-Sánchez et al., 2009), (Caton et al., 2010; Funes et al., 2007; Mues et al., 2009), (Beale et al., 2007) (Glycolysis, Gluconeogenesis, TCA Cycle, Pentose Phosphate Pathway, and Glycogen Metabolism), and (Whitfield et al., 2002) (G1/S).

### Biscuit Clustering and Normalization for Merging Samples

When attempting to merge immune systems from multiple patients, we observed that in some cases, cells were more similar to other cells from the same patient than they were to those of the same cell type. Figure 2A (left) shows scRNA-seq data from 9K immune cells from 4 breast cancer patients after normalization of cells to median library size, suggesting large differences between patients. Moreover, the tSNE projection does not suggest diverse subpopulations and structure beyond two main cell types of lymphoid and myeloid cells (Figure 2A left).

The differences across patients are likely caused by a large number of factors, both technical and biological. Technical factors include differences in machine, enzyme activity, lysis efficiency, and experimental protocol. These samples were also subject to operational variation during the clinical resection, transport, and handling. These factors all impact cell viability, which in turn affects the single cell RNA-seq library preparation, in particular molecular capture rate and sampling. Because molecular capture is a binary event, and the capture rates are very low, these technical variations often determine whether a given gene feature is observed in the data for a given cell.

These more technical artifacts, particularly in capture rate, are confounded with biological differences. This is particularly challenging in the case of immune cells, where activated cells have higher transcription rates. Thus, the number of captured molecules may be higher in activated immune cells, and therefore activation of immune cells can be convolved with total counts (Blackinton and Keene, 2016; Cheadle, 2005; Singer et al., 2016). As a result, methods that normalize all cells together may remove critical cell type-specific biological variation in the systems of interest such as immune activation and environmental response. Indeed, we see large differences in the number of activated T-cells across patients (Figure 2B), with more activated T-cells in the Triple Negative subtype as expected (Dushyanthen et al., 2015). Hence, normalizing by library size will likely remove these biological variations. Cell type-specific normalization is especially crucial in cases involving vast subtype diversity, such as immune cells including large macrophages and smaller lymphocytes (Vallejos et al., 2017).

To solve this problem we developed and applied the method “Biscuit” (short for Bayesian Inference for Single-cell ClUstering and ImpuTing) to simultaneously cluster cells and normalize according to their assigned clusters (Figure M2). This is done through incorporating parameters denoting cell-specific technical variation into a Hierarchical Dirichlet Process mixture model (HDPMM) (Görür and Rasmussen, 2010) (Figure M2A). This allows for inference of cell clusters based on similarity in gene expression as well as in co-expression patterns, while identifying and accounting for technical variation per cell (Figure M2B,C). Two key ideas that power Biscuit are the use of gene co-expression as a more robust means to identify cell types, and the normalization of each cell type separately to better account for cell type-specific effects on technical variation. The main idea behind the use of co-expression is that cell types not only share similar mean expression, but also share similar co-expression patterns (covariance) between genes. While mean expression can be more sensitive to capture efficiency, covariation is more robust to such effects. This similarity in co-variation can be used to improve normalization and in turn improve clustering, through the learning of cluster specific parameters.

By jointly performing normalization and clustering, we retain biological heterogeneity and avoid biases that result from independent clustering and normalization, and instead are able to match cells to clusters of the same cell type from different patients which may have very different sampling rates. Figure 2A (right) shows the same data from 4 tumors after normalization with Biscuit. Note that Biscuit does not use any information on sample IDs in the normalization, and normalization is only driven by cluster assignments. The Biscuit-normalized data shows that the differences in library-size normalized data were largely artifacts of normalization and batch effects. Furthermore, data from Biscuit shows richer structure suggesting diversity in cell types. Hence, we then applied Biscuit to data from all 8 tumors to infer the full diversity of immune cell types in the breast tumors, which identified 67 clusters, indicating significant diversity in both lymphoid and myeloid cell types (Figure 2C).

#### Summary of Biscuit model

Starting with the count matrix 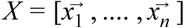, where each column 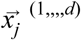 contains the expression (number of unique mRNA molecules) of *d* genes in cell *j*, the model assumes the log of counts 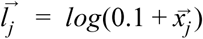 for each cell *j* = (1, …, *n*) follow a multivariate Normal distribution: 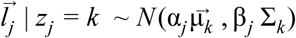 where *z*_*j*_ denotes the assignment of cell *j* to cluster *k*, and 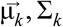 are the mean and covariance, respectively, of the *k*-th mixture component (cluster), and scalars α_*j*_, β_*j*_ are cell-dependent scaling factors used for normalization. Note that the assumption of log Normality has been verified using model mismatch and Lilliefors tests (Prabhakaran, S., Azizi, E., Carr, A., & Pe’er, D, 2016). Within a Bayesian model setting, we assign conjugate prior distributions to the parameters, namely a symmetric Dirichlet prior of the order *K*, a conjugate-family prior over μ_*k*_ as Normal and for Σ_*k*_ as Wishart (Dı́az-Garcı́a et al., 1997). A noninformative Normal prior is set for α*j* and an Inverse-gamma is set for β_*j*_. The full model specification is thus as below:

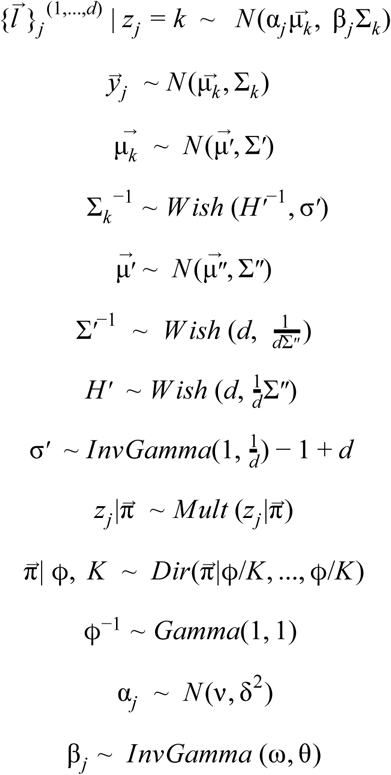

where 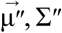 are the empirical mean and covariance across all cells, which are used to allow priors to adapt to different datasets (Figure M2B).

Parameters are inferred through a scalable Gibbs algorithm using the Chinese restaurant process (CRP) (Pitman, 2006), which also infers the number of clusters (*K*) (Figure M2C). The conditional posterior distributions for model parameters 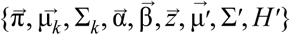 have analytical forms which we derived in (Prabhakaran, S., Azizi, E., Carr, A., & Pe’er, D, 2016).

Final cluster assignments were estimated from the mode of inferred distribution for assignment of cells to clusters (*z*_*j*_; ∀*j*) after removing a burn-in period. The mean of Gibbs samples for cluster-specific parameters (μ_*k*_, Σ_*k*_; ∀*k* = 1, …, *K*) and cell-specific parameters (α_*j*_, β_*j*_; ∀*j*) were used for further analysis and interpretation.

The goal of normalization is to transform the data from 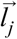 to 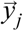 in which the expression is corrected for cell-specific factors α_*j*_, β_*j*_ using a linear transformation 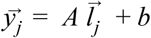 such that imputed expression for cell *j* follows *N*(μ_*k*_, Σ_*k*_) and hence all cells assigned to the same cluster follow the same distribution after correction. One transformation satisfying the above distributions for 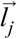 and 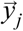 is 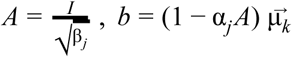.

This transformation also imputes dropouts in a cell by using the parameters of the cluster to which it has been assigned. The use of covariance parameters in the model ensures that intra-cluster heterogeneity is preserved after imputing. We show a systematic evaluation of the algorithm performance (on synthetic and real single cell data), its robustness, and its ability to impute dropouts, in (Prabhakaran, S., Azizi, E., Carr, A., & Pe’er, D, 2016).

#### Biscuit Implementation

The joint distribution of Biscuit with cluster- and cell-specific parameters is non-conjugate. Although inference of these parameters via MCMC-Gibbs is possible as presented in (Prabhakaran, S., Azizi, E., Carr, A., & Pe’er, D, 2016), overall runtime of the algorithm was much slower due to multiple covariance matrix inversions leading to slower chain convergence. In order to reduce the number of matrix inversions and enable faster chain mixing, we make use of the conjugate prior for the multivariate Gaussian, namely the Normal-inverse Wishart distribution, where the cluster means and covariances can be jointly inferred (Murphy., 2007).

The Normal-inverse-Wishart (NIW) distribution is the conjugate prior to the multivariate Gaussian distribution with unknown mean and covariance. It is a multivariate four-parameter continuous distribution with probability density function defined as follows:

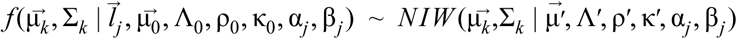

The inference equations for parameters based on NIW also have closed-forms:

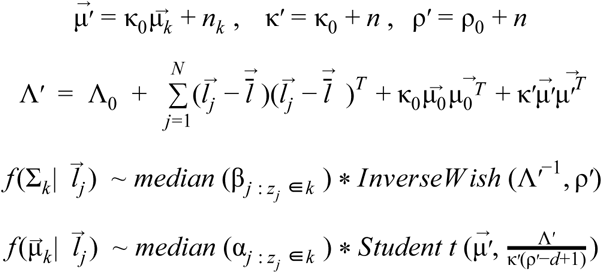

where 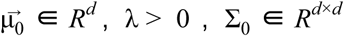 and is positive semi-definite andC ν > (*d* − 1). This scalable implementation of Biscuit in R can be found in:

https://github.com/sandhya212/BISCUIT_SingleCell_IMM_ICML_2016. The code is parallelized at multiple stages - the NIW-based inference engine works in parallel across blocks of genes ordered through the Fiedler vector. Next, cell assignments per block are consolidated into a confusion matrix via further parallelisation and the overall cluster means and covariances are computed using the law of total expectation, also performed in parallel. A dataset of 3000 cells and 1000 genes takes under two hours on a Cloud platform such as Amazon Web Services (AWS), which is at least 6 times faster than the implementation of the initial model. For the full dataset, the DPMM dispersion parameter was set to 100, gene batches were set to 50, and the number of iterations was set to 150.

### Entropy Metric to Evaluate Batch Effect Correction

To evaluate Biscuit’s ability to correct batch effects across data from all eight tumors (Figure 2C) and match immune subtypes across the tumors, we devised an entropy-based metric that quantifies the mixing of the normalized data across samples. We used this quantification since discerning biases in 8 patients on a 2D t-SNE projection is difficult, whereas visualizing differences between 4 patients is easier (Figure 2A). The entropy-based metric is computed as follows: We constructed a k-NN graph (k=30) on the normalized data using Euclidean distance, and computed the distribution of patients (tumors) *m* = 1, …, 8 in the neighborhood of each cell *j*, denoted as 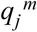. Then we computed Shannon entropy 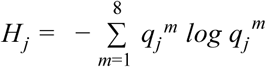 as a measure of mixing between patients, resulting in one entropy value *H*_*j*_ per cell *j*. High entropy indicates that the most similar cells come from a well mixed set of additional tumors, whereas low entropy indicates that the most similar cells largely come from the same tumor. Prior to Biscuit, most cells in the data had low entropy values, with 40% of the cells residing in a neighborhood of cells purely from the same tumor. We compare the distribution of entropies across all cells from all 8 tumors, before and after Biscuit, which reveals that the median of entropy shifts significantly towards higher mixing of samples after processing with Biscuit (Mann-Whitney U-test: U=1.7721e+09, p=0; Figure 2D). We thus conclude that Biscuit substantially corrected batch effects in this data.

### Quantification of Cell Type Enrichment in Tissues

To calculate cell type enrichments in each tissue (Figure 3B), we normalized each tissue to have equal cell count and created a 2-factor contingency table of cell types versus tissues. We then calculated *χ*^2^ enrichments for each tissue type and reported the test statistic and p-value.

Next, we wished to identify which pair of tissues had the greatest phenotypic overlap. To assess similarity in phenotype, we constructed a k-nearest neighbor graph (with k=10) in a uniformly selected subset of n=3000 cells from each tissue, reasoning that a cell’s closest neighbors in this high-dimensional embedding are the cells with the closest phenotypes. We then examined the overlap between each pair of tissues *u* and *v*, where n is the number of cells in the subset, k is the number of neighbors, and *u*, *v* ∈ {*tumor*, *normal*, *lymph node*, *blood*}:

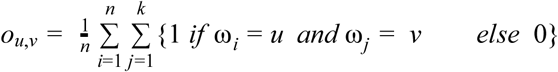

with ω_*i*_ denoting the tissue for cell *i* and *j* = 1, …, *k* denoting the neighbors of cell *i*. We built a null distribution from all overlaps between all pairs of tissues, and identified tumor and normal as having the highest frequency of being co-identified as neighbors. To test if this enrichment was significantly higher than the other shared-neighbor overlaps, we calculated the z-score of *o*_*tumor, normal*_ compared to the population of all pairwise overlaps (z=2.68), for which a z-test conferred a p-value of p=1.4e-4. Thus, we conclude that tumor and normal have the highest phenotypic overlap between any pair of tissues in our data. Both z-scores and p-values are reported. The above statistics are reported under the section “Environmental impact on the diversity of immune phenotypic states”.

### Creating a Global Immune Atlas using Biscuit

To generate a global atlas of immune cell types, we combined samples from all patients and tissues by applying Biscuit to the full set of *n*=62024 cells and *d*=14875 genes, resulting in a global atlas of *K*=95 clusters (Table S2) in which *n*=57143 cells had statistically significant cluster assignments. The remainder of cells had low library size and were hence removed from further analysis.

A subset of these clusters were identified as probable carcinoma or stromal populations through correlation with bulk gene expression datasets and marker gene expression. While these non-immune clusters may be of significant interest in their own right, they were beyond the scope of this paper and were therefore excluded from downstream analysis, leaving 47,016 cells in 83 clusters (Table S2).

Biscuit is a probabilistic model; hence, we inferred a probability distribution for all parameters. Figure S2A shows the posterior of assignment of cells to clusters, demonstrating a broader distribution for most naïve T cell clusters, suggesting that these clusters are less distinct as compared to other clusters. Other cell types exhibited a distinct, well separated mode. For simplicity, all further analysis is based on the mode of the probability of assignment of cells to clusters, such that each cell is assigned to one cluster only (Figure 2E).

Biscuit captures cell-type specific scaling factors using parameters α_*j*_, β_*j*_. Figure S2G shows that α_*j*_ values capture variation in library size (sum of total counts per cell *j*). Moreover, we observe different relationships with library size based on assigned clusters, highlighting the cluster-specific normalization that is done in Biscuit. We also observe prominent differences in distributions of α_*j*_, β_*j*_ parameters across cells from different patients (Figure S2H,I), emphasizing the differences resulting from technical factors across patients that are captured in Biscuit. The α_*j*_, β_*j*_ parameters represent each cell’s translational and rotational shifts within the Multivariate Normal distribution associated with the cluster assigned to that cell, by scaling the moments. By correcting these shifts, cells assigned to the same cluster will be normalized and follow the same distribution 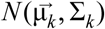.

In other words, Biscuit seeks to find if a cell can fit the distribution characterizing a given cluster, either if its mean and covariance match that of a cluster as is or match that of the cluster after scaling the cluster mean with its alpha and cluster covariance with its beta. The incorporation of covariance is key in that even if a cell has a low capture rate and thus a low number of detected genes, additional information from the gene-gene covariance patterns (modules of co-expressed genes) can be utilized to identify its cluster. Once a cell is assigned to a cluster, the distribution characterizing that cluster is then used to normalize and impute the expression of a cell, and thus corrects for downsampling and also imputes dropouts.

### Cluster Robustness

To evaluate cluster robustness, we performed 10-fold cross-validation, independently clustering and normalizing on random subsets of data. For each of 10 subsets, we ran Biscuit to obtain a set of clusters. To compare the results across the 10 subsets, we computed the confusion matrix indicating the probability of each pair of cells *j*, *j*′ being assigned to the same cluster: *P* (*z*_*j*_ = *z*_*j*_′). Figure S2C illustrates box plots for the probabilities of co-clustering (across 10 subsets) for every pair of cells that are assigned to the same cluster in the analysis of the full dataset. The average co-clustering probability in each cluster ranges between 92%-100%, showing remarkable robustness of clusters.

Additionally, we performed leave-one-patient-out analysis similar to the 10-fold cross-validation above. In this analysis we hold out one patient at a time and compute the probability of co-clustering of cells in the other 7 patients. As expected, this probability slightly drops for smaller clusters biased to one patient, but is otherwise highly robust and stable (Figure S2C).

### Mixing of Samples in Clusters

We observed that clusters displayed differing amounts of mixing between samples (Figure 2G). To further quantify the exact degree of mixing (between patients) in each cluster, we defined an entropy-based metric. We used bootstrapping to correct for cluster size (which ranged from over 8900 cells to just over 30 cells), such that we uniformly sampled 100 cells from each cluster and computed the distribution of patients across these cells, and then computed the Shannon entropy for this distribution. We repeated this procedure 100 times for each cluster, to achieve a range of entropy values per cluster. Figure S2D shows box plots for entropy values for each cluster, with the order of clusters based on their mean entropy. It also shows box plots for entropy values for each cell type, computed in a similar fashion. Clusters with entropy of 0 denote entirely patient-specific clusters. Figure S2D shows that there is a continuous range of entropy values, and thus a full range of sample specificity versus mixing, across clusters. For cell types it similarly shows a range of entropy values, with intratumoral macrophages and CD4+ T EM cells showing the highest degree of patient specificity.

The 10 patient-specific clusters with zero entropy are listed in the table below, showing that they span all cell types but are all specific to tumor tissue:

| Cluster ID | Most correlated cell type in Dmap (Novershtern 2011) | Most correlated cell type in Garvan (Jeffrey 2006) | Final Annotation based on DEGs and correlations | Cluster size (#cells) | % in Tumor | Patient |
| --- | --- | --- | --- | --- | --- | --- |
| 65 | BCELLA2 | bcells | B: | 56 | 100 | BC5 |
| 70 | TCELLA7 | central_memory_T_cells | T:CD4+EM | 49 | 100 | BC5 |
| 73 | ERY1 | mast_cells | MAST: | 47 | 100 | BC4 |
| 82 | TCELLA7 | th1_cells | T:Reg | 38 | 100 | BC5 |
| 86 | MONO2 | neutrophils | MONOCYTE: | 37 | 100 | BC5 |
| 89 | GRAN2 | neutrophils | NEUTROPHIL: | 35 | 100 | BC5 |
| 91 | MONO2 | macrophages_LPSH | MONOCYTE: | 35 | 100 | BC5 |
| 93 | TCELLA7 | central_memory_T_cells | T:CD4+EM | 33 | 100 | BC5 |
| 94 | MONO2 | neutrophils | MONOCYTE: | 33 | 100 | BC5 |
| 95 | TCELLA8 | central_memory_T_cells | T:CD4+CM | 33 | 100 | BC5 |

Two patients (BC4, BC5) contribute to these clusters. BC5 is a TNBC case, and hence is expected to have differences in immune phenotypes.

### Distances between Clusters

Biscuit provides a parametric characterization of each cluster using a multivariate Normal distribution, which allowed us to directly compare the distributions that define these clusters. While Euclidean distances are defined for vectorial objects and operate under a Cartesian coordinate system, Euclidean distance with non-vectorial objects such as probability distributions requires embedding them in Euclidean space. Such embeddings are non-unique and lead to loss of information. It is therefore advisable to use these non-vectorial objects as they are and to work with the objects’ pairwise similarities or distances instead. One such distance metric, which is effective at comparing pairwise probability distributions, is the Bhattacharyya distance (BD) (Bhattacharyya, 1990).

We defined distances between each pair of clusters *k*, *k*′ with distributions *p*_*k*_ and *p*_*k*′_ as *BD* =− *log*(*BC*(*p*_*k*_, *p*_*k*′_)) where *BC* is the Bhattacharyya coefficient measuring similarity (overlap) of the distributions. We use the BD to compute distances between pairs of inferred clusters’ moments to create the Bhattacharyya kernel. The Bhattacharyya kernel has closed forms for any exponential distribution including the (multivariate) Gaussian distribution (Jebara, T., Kondor, R., & Howard, A., 2004), which is Biscuit’s underlying data-generation distribution. For the case of multivariate normal distributions: 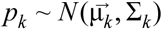 and 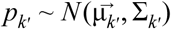:

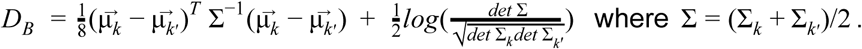

Figure S2B shows the heatmap of pairwise distances between all pairs of clusters.

A geometric interpretation of BD is that, via its cosine formulation, the distance subsumes a full hypersphere and the centre of the hypersphere is the centroid (mean) of the cluster, whereas the Euclidean distance only covers a quarter of the hypersphere with the center at the origin.

### Contribution of Covariance in Defining Clusters

We used the above Bhattacharyya distance (BD) metric to study the contribution of Biscuit’s covariance parameters to characterizing clusters of different cell types. First, we computed the BD distance between pairs of clusters of the same type (T, monocytic, NK, B) and compared these to distances between pairs of clusters of different cell types (e.g. a T cell cluster and a monocytic cluster). Figure S2E shows violin plots for distances between pairs of clusters with dots (overlaid on violins) representing cluster pairs; violins are sorted based on median distance. As reference, we also split each cluster into two halves and computed the empirical BD between two splits (shown at the left end in Figure S2E). We observe that, overall, pairs of clusters of different types are more distant than pairs of clusters from the same type, as expected.

We then computed these same pairwise distances while removing the contribution of mean parameters for each cluster, via setting 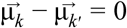 and computing the distance only based on covariance parameters of the pair of clusters Σ_*k*_, Σ_*k*′_ (Figure S2F). We observed that pairs of T cell clusters or monocytic clusters still show prominent distances, and therefore covariance parameters have a crucial role in defining these clusters.

Such varying covariance parameters span the entire genome, but include functionally important genes. For example, the degree of co-expression of genes associated with M1 and M2 signatures also varied widely within clusters in a manner not fitting the functional M1/M2 annotation. As an example, expression of CD64 in cluster 23 exhibited varying degrees of positive covariance with FN1, MMP14, MSR1, cathepsins, MARCO, and VEGFB, but co-varied slightly negatively with chemokine CCL18 (Figure 7J).

### Defining Phenotypic Volume

One of the main aims of this paper was to compare immune phenotypes across tissues, including peripheral blood and lymph node tissue as references in which the immune phenotypes typically present are better understood. The global atlas from merged tissues revealed that clusters differ appreciably in their proportions across tissues (Figure 3A-C). For example, we observed 3 blood-specific clusters with CD4+ naive characteristics, as well as a lymph node-specific T cell cluster. Conversely, we observed a large range of shared clusters between normal and tumor tissues (Figure 3B; Table S2). Interestingly, the clusters observed in normal tissue were a subset of those observed in tumor tissue. We also observed an increase in the variance of gene expression in tumor compared to normal tissue (Figure 3D,E; S3B,C), which prompted us to explore further the range of phenotypic diversity between normal and tumor tissue. We were especially interested to find whether the observed increase in variance of expression is due to activation of additional processes in tumor that are independent of those active in normal.

Hence, to quantify the diversity of phenotypes, we defined a metric of phenotypic volume or space occupied by a cell type (e.g. T, NK, or monocytic), such that if more independent phenotypes are active, the total volume spanned by the phenotypes would be larger than an alternative case wherein phenotypes are dependent.

We therefore defined “phenotypic volume” (*V*) for a subpopulation of cells as the determinant of the gene expression covariance matrix for that subpopulation, which considers covariance between all gene pairs in addition to their variance. The (symmetric) covariance matrix can be written as 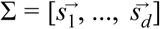 where 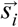 for *i* = 1, …, *d* is a vector containing covariance between gene *i* and all other genes. Its determinant *det* (Σ) is equal to the volume of a parallelepiped spanned by vectors of the covariance matrix (Tao and Vu, 2005).

For example, if the covariance values between a gene *i* and other genes is very similar to that of another gene *i*′ and other genes, such that 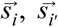 are dependent, gene *i*′ does not add to the volume. Extending this to all genes, we sought to evaluate whether the increase in expression variances (Figure 3D,E) are associated with phenotypes activated in tumor that are independent from those in normal tissue, i.e. are novel independent phenotypes observed in tumor that suggest additional mechanisms and pathways being activated in tumor.

In a simplified case with only two phenotypes, the determinant, which is equal to the area of the parallelogram spanned by two vectors representing the phenotypes, is larger if the phenotypes are independent, but would be equal to zero if they are fully dependent. With more than two phenotypes, we are then interested in measuring the volume of the parallelepiped spanned by these phenotypes. The (pseudo-)determinant can also be computed as the product of nonzero eigenvalues of the covariance matrix:

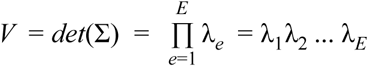

To quantify the change in phenotypic volume from normal to tumor, we computed this volume metric for each major cell type of T, monocytic, and NK cells. To correct for the effect of differences in the number of cells across cell types and tissues, we uniformly sampled 1000 cells with replacement from each cell type per tissue and computed the empirical covariance between genes based on imputed expression values for that subset of cells. This was followed by singular-value decomposition (SVD) of each empirical covariance matrix and computation of the product of nonzero eigenvalues as stated in the equation above. B cells were not included in this comparison due to the very small number of B cells in normal tissue. Given the high number of dimensions (genes), the volumes were normalized by the total number of genes (*d*). For robustness, this process was repeated 20 times to achieve a range of computed volumes for each cluster in each tissue, which are summarized with box plots (Figure 3F) showing statistically significant expansions of volume in tumor compared to normal in all three cell types (Mann-Whitney U-test p=0 for all three cell types).

### Diffusion Component Analysis

We used diffusion maps (Coifman et al., 2010; Setty et al., 2016) as a nonlinear dimensionality reduction technique to find the major non-linear components of variation across cells. We computed diffusion components in each cell type separately using the Charlotte Python package, which implements diffusion maps as described in (Coifman et al., 2005). To account for differences in cell density and cluster size, we used a fixed perplexity Gaussian kernel with perplexity 30, with symmetric Markov normalization and t=1 diffusion steps. We selected t=1 diffusion steps because this approximates diffusion of information for each cell through its 20 nearest neighbors in our data. This is a conservative value, but we wanted to ensure that information did not diffuse beyond the borders of our smallest cluster (30 cells). To focus on processes explaining variation specifically within major cell types, we performed diffusion map analyses separately on all cells labeled as T cells and myeloid cells, respectively.

In the case of T cells, the first two diffusion components identified two isolated clusters (Figure 3A): The first was cluster 9, which is quite distinct from other T cell clusters as measured by Bhattacharyya distance (Figure S2B) and shows characteristics similar to NKT cells (Table S3); the second was cluster 20, which is a blood-specific naive T cell cluster predominantly from one patient (Table S2). Since these two clusters were very distant from other T cell clusters according to a variety of comparative metrics, the two components corresponding to them were removed from further analysis, as we wished to focus on studying components that are informative of heterogeneity across multiple clusters.

The next components were labeled as T cell activation, Terminal Exhaustion, and Hypoxia, respectively, (Figure 4A) as they were most highly correlated with the corresponding gene signatures (Table S4). The subsequent component is labeled as Tissue Specificity, as it separates cells primarily on the basis of their tissue of origin and helps explain heterogeneity in T cells across tissues.

In the separate analysis of myeloid cells, several isolated clusters, namely neutrophils and mast cells, were not considered in the analysis, with the goal of identifying components explaining variation across multiple monocytic clusters (Figure 7B, S7A). The first component identified a trajectory of macrophage (TAM) activation. The second and third branches together captured a more gradual trajectory from blood monocytes to intratumoral monocytes (Figure 7B-D; S7B-D). The “blood terminus” of the trajectory correlated with expression of co-stimulatory gene ITGAL, but also with several tumor growth-promoting genes, i.e. fibroblast and epidermal growth factors, as well as IL-4 (Figure S7B), proposed to support the M2 type of macrophage activation (Mantovani and Locati, 2013; Mills et al., 2000). The other end of the trajectory, populated by intratumoral monocytes, was characterized by high expression of activation and antigen presentation-related genes encoding CD74 and HLA-DRA, but also an IFN-inducible gene encoding ISG15, which has been described to be secreted by TAMs and enhance stem-like phenotypes in pancreatic tumor cells (Figure S7B) (Sainz et al., 2014).

The fourth branch correlated with canonical plasmacytoid dendritic cell (pDC) markers such as LILRA4, CLEC4C (CD303), and IL3RA. The most discrete of the myeloid components, this branch separated the lone pDC cluster (41) from the other myeloid-monocytic cell clusters (Figure 7B, E, G, H; S7B, C, E). This subset was also the only monocytic cluster common between the tumor and the lymph node; it featured high levels of granzyme B (GZMB) (Figure S7B), which has been proposed to be a means, by which pDCs may suppress T cell proliferation in cancer (Jahrsdörfer et al., 2010; Swiecki and Colonna, 2015).

### Significance of Differences in Covariances in Raw Data

To verify that the differing covariance patterns in Figures 5 and 7 were not the result of computational modeling decisions, we tested the difference in covariance in raw unnormalized data (prior to Biscuit). As the raw data involves significant amount of dropouts, co-expression patterns and their signs cannot be easily visualized or inferred.

Hence, we performed hypothesis testing accounting for the level of dropouts by comparing the observed empirical covariance between a pair of genes *i*, *i*′ to a null distribution for the gene pair in which co-expression patterns are removed. We assume the null hypothesis to be the case where covariance between a specific gene pair for a given cluster is the same across all clusters.

Specifically, to test whether 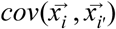 in a cluster *k*, with 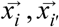 being expressions of genes *i*, *i*′ across cells assigned to cluster *k*, is significantly different from that in all other clusters, we used bootstrapping and permutation testing as follows: We started by generating a null distribution for the covariance between a pair of genes by first uniformly sampling a subset of cells from all clusters, with the subset being the same size as cluster *k*. Then, to further remove existing structures of co-expression in cells, we permuted the cell labels for gene *i*′ (while retaining labels for gene *i*) and computed empirical covariance between the two genes in this subset of “scrambled” cells. We repeated this on 10,000 subsets to achieve a null distribution of 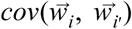 where *w*_*i*_, *w*_*i′*_ are the expressions of gene *i*, *i*′ in the sets of scrambled cells. We then compared the observed 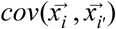 (marked with a star in Figure S5A, S7F) to the null distribution, which was rejected for that pair of genes if p-value<0.05 indicating that the covariance is significantly different in cluster *k* compared to all other clusters.

We concluded that the signal is also apparent in raw unnormalized data for all the aforementioned clusters and we observe a range of covariance values with different signs between GITR and CTLA4 across Treg clusters (Figure S5B), and similarly different values in covariance between MARCO and CD276 in TAM clusters (Figure S7G).

### Comparison of Treg clusters to previous studies

Recent studies examining Treg cell gene expression profiles in breast, lung, and colorectal tumors found differences in gene expression between tumor and normal tissue-associated cells (De Simone et al., 2016; Plitas et al., 2016). CCR8, CD177, LAYN, and MAGEH1 were expressed highly by tumor Treg cells. Flow cytometric analyses revealed that CD177 was expressed on a subset of tumor-associated Treg cells, while CCR8 was expressed on all Treg cells. Our single cell analyses confirmed that CD177 is expressed highly in some Treg clusters. While flow cytometric analysis of LAYN and MAGEH1 was not attainable due to the absence of reagents, we observed high expression of LAYN and MAGEH1 in all Treg cell clusters, suggesting that the pattern of expression of these two genes in Treg cells is similar to that of CCR8.

### Continuity of Cells along Components

The prevalent view is that relatively few states of differentiation of lymphoid and myeloid cells define the local tumor environment (Philip et al., 2017); (Gabrilovich, 2017); (Wherry and Kurachi, 2015); (Akondy et al., 2017; Kumar et al., 2018; Youngblood et al., 2017), and that these distinct states are linked to clinical outcomes such as cancer progression and response to anti-tumor therapies. We were interested to know whether cells show continuity as opposed to defined cell states along various diffusion components. For example, we were interested to know whether T cells exhibit defined states with different activation levels. For this, we computed the distribution of cells projected on each diffusion component and then used Hartigan’s Dip Test (Hartigan and Hartigan, 1985) to test whether the distribution of cells is unimodal (broad continuum of cells) or alternatively multimodal (representative of multiple defined states) with p<0.01. In Figure S4A, we observe that in the case of the T Cell Activation component, the null hypothesis of unimodality is not rejected, indicating that the distribution of cells is similar to a broad unimodal distribution as opposed to a multimodal distribution with defined states. In contrast, other components (such as the Tissue Specificity Component) exhibit multimodal distributions with distinct modes implying distinct states (in this case corresponding to various tissues).

In the case of myeloid cells, the null hypothesis of unimodality is rejected in all diffusion components, indicating that myeloid cells lie in distinct states along all major components explaining variation across cells that were analyzed (Figure S7A).

### Differences across patients

We performed Hallmark gene set enrichment on standard deviations of genes across patients, to identify signatures most variable across patients. Further details of distributions of a number of these signatures such as Hypoxia are shown in Figure 1E and S1E,F,G.

| NAME | FDR q-val | FWER p-val |
| --- | --- | --- |
| HALLMARK_TNFA_SIGNALING_VIA_NFKB | 0 | 0 |
| HALLMARK_ANGIOGENESIS | 0 | 0 |
| HALLMARK_COMPLEMENT | 0 | 0 |
| HALLMARK_ALLOGRAFT_REJECTION | 0 | 0 |
| HALLMARK_INTERFERON_GAMMA_RESPONSE | 0 | 0 |
| HALLMARK_APOPTOSIS | 0 | 0 |
| HALLMARK_INTERFERON_ALPHA_RESPONSE | 0 | 0 |
| HALLMARK_COAGULATION | 0 | 0 |
| HALLMARK_EPITHELIAL_MESENCHYMAL_TRANSITION | 0 | 0 |
| HALLMARK_HYPOXIA | 0 | 0 |
| HALLMARK_IL6_JAK_STAT3_SIGNALING | 0 | 0 |
| HALLMARK_MYC_TARGETS_V1 | 0 | 0 |
| HALLMARK_OXIDATIVE_PHOSPHORYLATION | 0 | 0 |
| HALLMARK_PI3K_AKT_MTOR_SIGNALING | 0 | 0 |
| HALLMARK_REACTIVE_OXYGEN_SPECIES_PATHWAY | 2.67E-04 | 0.004 |
| HALLMARK_MTORC1_SIGNALING | 3.13E-04 | 0.005 |
| HALLMARK_P53_PATHWAY | 2.94E-04 | 0.005 |
| HALLMARK_INFLAMMATORY_RESPONSE | 2.78E-04 | 0.005 |
| HALLMARK_KRAS_SIGNALING_UP | 3.16E-04 | 0.006 |
| HALLMARK_UV_RESPONSE_UP | 3.00E-04 | 0.006 |
| HALLMARK_CHOLESTEROL_HOMEOSTASIS | 3.33E-04 | 0.007 |
| HALLMARK_ANDROGEN_RESPONSE | 5.46E-04 | 0.012 |

More importantly, we observe more heterogeneity in these signatures within each tumor (Figure 1E and S1E,F,G), motivating the characterization of various cell states that vary in frequency between patients.

### Preprocessing of paired 5’ scRNA-seq and TCR-seq data from 10x

Data was collected from FACS-sorted CD3+ T cells generating two technical replicates from the BC9 tumor, one replicate from the BC10 tumor (due to a lower number of T cells), and two technical replicates from the BC11 tumor. Technical replicates were run on separate 10x lanes, but originate from the same sample after tissue dissociation. The sequencing data from each replicate was preprocessed separately using Cell Ranger 2.1.1, available from 10x Genomics. The clonotype comparison feature in Loupe Cell Browser (also from 10x Genomics) combined with custom scripts was then used to pool TCR clonotypes across replicates by matching CDR3 sequences from both alpha and beta chains across replicates of the same tumor. For two cells to be assigned to the same clonotype they had to share both alpha and beta sequences. We achieved a paired transcriptome and TCR sequence for 12962, 4677, and 9436 T cells from BC9, 10, and 11, respectively, resulting in a total dataset of 27,075 T cells. The median number of molecules per cell in this dataset was 4780. The frequencies of the most dominant clonotypes per patient are shown in FIgure S6D.

### Analysis of 10x Genomics paired TCR and scRNA sequencing data

The single-cell expression count matrices for 27,075 T cells from three tumors (BC9-11) were combined, and these cells were normalized and clustered with Biscuit with parameters: 500 iterations; gene batch size set to 50; alpha (dispersion parameter) set to 200, identifying 34 clusters. The number of genes detected in each of the 34 clusters are shown Figure S6B, showing an increase in capture rate compared to inDrop data shown in Figure S3A.

To evaluate the extent to which the 38 T cell clusters previously inferred from BC1-8 (using inDrop) generalize to these new cases BC9-11 (profiled with 10x), we computed the correlations between empirical means of previous and new clusters (Figure 6A). The correlations were computed using the expression of previously identified differentially expressed genes (listed in Table S3) for each of the 38 clusters. Furthermore, to correct for different distributions between the two technologies, the datasets were first quantile normalized prior to computing these correlations. Rows in Figure 6A are ordered based on the cluster from the previous set most correlated with each of the new 34 T cell clusters, and columns (new clusters) are ordered by size, i.e. number of cells in the cluster. This heatmap shows near one-to-one mapping of each cluster identified in the three new tumors to a cluster previously inferred from eight tumors, confirming the generalizability of the clusters.

Furthermore, to account for the gene distributions used to characterize clusters in Biscuit (recall, Biscuit clusters are defined by both mean and covariance, with covariance playing an important role, see figure S2E,F), we also computed the Bhattacharyya coefficient between the old and new clusters as a metric for the similarity of distributions of the differentially expressed genes (Figure 6B), showing even clearer near one-to-one-mapping of clusters characterized by both mean and covariance.

### Evaluation of the role of TCR diversity in driving a continuous spectrum of T cell activation

The activation state of each T cell was computed by taking the mean expression of all genes in the T cell activation signature (Table S4). The distribution of activation states across all T cells is shown in Figure S6C, reproducing the observation of continuity along activation as opposed to defined states in three new cases profiled with a different technology (10x rather than inDrop). We confirmed this continuity with Hartigan’s Dip test (dip statistic = 0.0578; p=0). We also observed continuity of T cell distribution along the first diffusion component across these cells from BC9-11, similar to the first informative diffusion component across T cells from BC1-8.

Using paired TCR sequencing of the same cells, we identified 9562, 3080, 6423 unique clonotypes from BC9, 10, and 11, respectively. The distributions of activation states across T cells from each clonotype are shown in Figure 6C and Figure S6E.

To determine whether clonotype identity has any role in explaining the continuity of T cell activation, we compared the entropy of the distribution of activation states in each clonotype with that of random T cells from various clonotypes. Specifically, for each clonotype, we first computed the entropy for its distribution of activation states. Then we created a null distribution by uniformly subsampling the same number of T cells as in the clonotype (to control for sample size) from the pool of all cells regardless of clonotype. This was followed by computing the entropy in each subsample, achieving a null distribution for entropy. We then compared the clonotype entropy to this null to test whether the clonotype has significantly less entropy, which would indicate it is not as uniformly distributed as random subsamples of T cells. We indeed observed significantly low entropy values for all top 30 clonotypes in each tumor (p=0), showing that individual clonotypes are confined to certain activation states and are not as widely distributed as are all T cells, and further that this is not an artifact of clonotypes containing less cells than the entire dataset. This suggests that individual clonotypes are not as uniformly distributed as are the aggregation of all clonotypes. Thus, TCR diversity partially explains the observed broad, continuous range of T cell activation, and aggregating all clonotypes thus obscures transitional activation states. For example, clonotypes 10, 11, and 12 in BC9 are centered in distinct regions of the activation component, and if the clonotype identities were not known, the aggregation would result in a more uniform distribution. Extending this to thousands of clones observed per tumor in part explains why we observe a continuous distribution of activation across all T cells.

Nevertheless, a surprisingly wide range of activation states was observed in each individual clonotype (Figure 6C; S6E). This is summarized with the distribution standard deviations of each clonotype listed in these figures. To assess the extent to which clonotype identities explain variability in T cell activation, we performed one-way ANOVA tests on the top 100 most frequent clonotypes in each tumor. ANOVA partitions the total variation into variation between versus within clonotypes, and tests if the means of clonotypes are significantly different. The percentage of variation in activation states across all cells that can be explained by clonotype identities in each tumor are listed below:

BC9: 52%, F-statistic=28.2095, p=3.8298e-100

BC10: 48%, F-statistic=35.2584, p= 4.5597e-139

BC11: 32%, F-statistic=10.6126, p=9.8365e-40

Moreover, we computed the average pairwise variation between the top 20 most frequent clonotypes using the one-way ANOVA F-test. The ratio of average pairwise variation between the top 20 most frequent clonotypes to the average variation within a clonotype in each tumor is as follows:

BC9: 0.54, F-statistic=49.0512, p=4.0576e-119

BC10: 0.46, F-statistic=42.1998, p=3.7453e-111

BC11= 0.29, F-statistic=24.3593, p= 2.8834e-71

These results together suggest that TCR diversity may not be the primary driver of the continuity of T cell activation, implying the existence of additional key factors contributing to activation state.

### Evaluation of the role of TCR repertoire in explaining phenotypic states in T cells

The proportions of the most frequent clonotypes composing clusters are shown in Figure 6E. In these heatmaps only CD8+ T cell and Treg clusters are shown, as the majority of dominant clonotypes were present in CD8 and Treg clusters (Figure S6F). Clusters (columns) are ordered by T cell activation.

In some cases, clusters that shared a clonotype exhibited similar levels of activation, apparent in Figure 6E as horizontal stretches in the heatmap. In other cases clusters that shared a clonotype were similar in gene expression signatures relating to environmental stimuli. For example, clonotype 9 from tumor BC9 is present in clusters T11 and T12, which have very similar expression levels across nearly all of the environmental signatures (Figure 6D,E). Additionally, clonotype 16 in BC9 is present in two clusters that are both high in exhaustion, whereas clonotype 20 in BC9 is present in clusters high in anergy and glucogenesis. Clonotype 21 in BC10 is present in clusters with high expression in the G1/S signature and low expression of exhaustion, hypoxia, TCA cycle, and anti-inflammatory signatures. The full list of differentially expressed genes (computed with t-test p<0.1) for each clonotype in each tumor are listed in supplementary Table S8.

All 27,075 T cells are projected together using t-SNE in Figure 6F, and are colored by specific marker genes, mean expression of the T cell activation signature, sample identity, and finally Biscuit assignment. Individual select clonotypes for each tumor are projected in Figure S6F (right) using the same t-SNE coordinates, showing that each clonotype spans a confined region and is present in a limited set of similar clusters. Note that clonotypes are not shared across patients, and hence these projections showing clonotype identity are provided separately for each patient. To confirm the statistical significance of the similarity of clusters sharing clonotypes, we computed the sum of pairwise Bhattacharyya coefficient (similarities) between clusters present in each clonotype and compared that value to a null distribution of the sum of pairwise similarities between 1000 random subsamples of (the same number of) clusters, resulting in p<0.001 for each of the top 20 most frequent clonotypes in all three tumors.

### CyTOF Data Processing and Analysis

To confirm the role of covariance in characterizing Treg clusters, we collected CyTOF data from three additional breast tumors. We designed a panel that included markers which are either differentially expressed in a Treg cluster and/or show different covariance patterns across Treg clusters. This panel is included in Table S9.

Cells from each tumor sample were barcoded together with stock human PBMCs as carrier cells, using three non-overlapping isotope channels for each. Thus two separate samples, TILs from a tumor and stock PBMCs, were run together and present in subsequent .fcs files. These two samples were debarcoded as previously described (Zunder et al., 2015), and the resulting debarcoded tumor files were normalized together using bead standards (Finck et al., 2013). Bead-normalized single cell measurement values were then transformed using the hyperbolic inverse sine function with a cofactor of 5. Samples were manually gated by event length, live/dead discrimination, and CD3, resulting in 6559 T cells from three tumors available for downstream analysis.

The data was then clustered using PhenoGraph (Levine et al., 2015) on 10 principal components (chosen based on the knee point of eigenvalues) calculated using all channels, with the parameter k=25 for the number of nearest neighbors. Data was visualized using t-SNE (Maaten and Hinton, 2008)), shown in Figure S5C.

We observed two Treg clusters shown in Figure S5C (labeled cluster A, B) resembling two previously identified Treg clusters: Treg cluster A is similar to cluster 82 differentially expressing CD25 (t-test p=0) and showing no covariance between GITR and CTLA-4 while Treg cluster B resembles previously inferred cluster 46, differentially expressing TIGIT (t-test p=0) and showing strong positive covariance between CTLA-4 and GITR.

### DATA AND SOFTWARE AVAILABILITY

The sequencing data presented in this paper will be available for download on GEO data repository.

SEQC is available on https://github.com/ambrosejcarr/seqc.git and Biscuit is available on https://github.com/sandhya212/BISCUIT_SingleCell_IMM_ICML_2016.

## Supplementary Table Legends

**Table S1.** Aggregated quality control and metric information for each sequencing sample. Columns A-C contain metadata on the patient, tissue of origin, and sample number. Related to Figure S1B.

**Table S2.** Annotations of clusters inferred in full breast immune atlas (across all patients and tissues) and their proportions across tissues and patients.

**Table S3.** List of differentially expressed genes in clusters listed in Table S2 (sheet 1); the subset of differentially expressed immune-related genes (sheet 2).

**Table S4.** List of gene signatures; sources for T cell signatures are stated in STAR Methods; M1 and M2 signatures are listed with sources in second and third sheet.

**Table S5.** Full hallmark GSEA enrichment results for gene variance in T-cells (Sheet 1), NK cells (Sheet 2), and Monocytic Cells (Sheet 3). These enrichments expand upon partial lists shown in Figure 3 and S3.

**Table S6.** List of capture reads, molecules, cells, and genes per sample replicate

**Table S7.** Frequencies of lymphocyte subsets analyzed by flow cytometry. The data represent the percent of all CD45+ cells.

**Table S8.** List of differentially expressed genes in the top 30 most frequent TCR clonotypes in each of the three tumors (BC9, BC10, BC11) in separate sheets.

**Table S9.** CyTOF Panel designed to validate different covariances in Treg clusters

## References

Amir el, A.D., Davis, K.L., Tadmor, M.D., Simonds, E.F., Levine, J.H., Bendall, S.C., Shenfeld, D.K., Krishnaswamy, S., Nolan, G.P., and Pe’er, D. (2013). viSNE enables visualization of high dimensional single-cell data and reveals phenotypic heterogeneity of leukemia. Nature biotechnology 31, 545–552.

Cheadle, C., Fan, J., Cho-Chung, Y.S., Werner, T., Ray, J., Do, L., Gorospe, M., and Becker, K.G. (2005). Control of gene expression during T cell activation: alternate regulation of mRNA transcription and mRNA stability. BMC Genomics 6, 75.

Chevrier, S., Levine, J.H., Zanotelli, V.R.T., Silina, K., Schulz, D., Bacac, M., Ries, C.H., Ailles, L., Jewett, M.A.S., Moch, H., et al. (2017). An Immune Atlas of Clear Cell Renal Cell Carcinoma. Cell 169, 736–749 e718.

Coifman, R.R., Lafon, S., Lee, A.B., Maggioni, M., Nadler, B., Warner, F., and Zucker, S.W. (2005). Geometric diffusions as a tool for harmonic analysis and structure definition of data: Diffusion maps. Proceedings of the National Academy of Sciences of the United States of America 102, 7426–7431.

Dushyanthen, S., Beavis, P.A., Savas, P., Teo, Z.L., Zhou, C., Mansour, M., Darcy, P.K., and Loi, S. (2015). Relevance of tumor-infiltrating lymphocytes in breast cancer. BMC Med 13, 202.

Engblom, C., Pfirschke, C., and Pittet, M.J. (2016). The role of myeloid cells in cancer therapies. Nature reviews Cancer 16, 447–462.

Fan, X., and Rudensky, A.Y. (2016). Hallmarks of Tissue-Resident Lymphocytes. Cell 164, 1198–1211.

Finger, E.C., and Giaccia, A.J. (2010). Hypoxia, inflammation, and the tumor microenvironment in metastatic disease. Cancer metastasis reviews 29, 285–293.

Franklin, R.A., Liao, W., Sarkar, A., Kim, M.V., Bivona, M.R., Liu, K., Pamer, E.G., and Li, M.O. (2014). The cellular and molecular origin of tumor-associated macrophages. Science 344, 921–925.

Gaublomme, J.T., Yosef, N., Lee, Y., Gertner, R.S., Yang, L.V., Wu, C., Pandolfi, P.P., Mak, T., Satija, R., Shalek, A.K., et al. (2015). Single-Cell Genomics Unveils Critical Regulators of Th17 Cell Pathogenicity. Cell 163, 1400–1412.

Haghverdi, L., Buettner, F., and Theis, F.J. (2015). Diffusion maps for high-dimensional single-cell analysis of differentiation data. Bioinformatics 31, 2989–2998.

Jeffrey, K.L., Brummer, T., Rolph, M.S., Liu, S.M., Callejas, N.A., Grumont, R.J., Gillieron, C., Mackay, F., Grey, S., Camps, M., et al. (2006). Positive regulation of immune cell function and inflammatory responses by phosphatase PAC-1. Nat Immunol 7, 274–283.

Jimenez-Sanchez, A., Memon, D., Pourpe, S., Veeraraghavan, H., Li, Y., Vargas, H.A., Gill, M.B., Park, K.J., Zivanovic, O., Konner, J., et al. (2017). Heterogeneous Tumor-Immune Microenvironments among Differentially Growing Metastases in an Ovarian Cancer Patient. Cell 170, 927–938 e920.

Joller, N., Lozano, E., Burkett, P.R., Patel, B., Xiao, S., Zhu, C., Xia, J., Tan, T.G., Sefik, E., Yajnik, V., et al. (2014). Treg cells expressing the coinhibitory molecule TIGIT selectively inhibit proinflammatory Th1 and Th17 cell responses. Immunity 40, 569–581.

Josefowicz, S.Z., Lu, L.F., and Rudensky, A.Y. (2012). Regulatory T cells: mechanisms of differentiation and function. Annu Rev Immunol 30, 531–564.

Kumar, V., Patel, S., Tcyganov, E., and Gabrilovich, D.I. (2016). The Nature of Myeloid-Derived Suppressor Cells in the Tumor Microenvironment. Trends in immunology 37, 208–220.

Lavin, Y., Kobayashi, S., Leader, A., Amir, E.D., Elefant, N., Bigenwald, C., Remark, R., Sweeney, R., Becker, C.D., Levine, J.H., et al. (2017). Innate Immune Landscape in Early Lung Adenocarcinoma by Paired Single-Cell Analyses. Cell 169, 750–765 e717.

Levine, J.H., Simonds, E.F., Bendall, S.C., Davis, K.L., Amir el, A.D., Tadmor, M.D., Litvin, O., Fienberg, H.G., Jager, A., Zunder, E.R., et al. (2015). Data-Driven Phenotypic Dissection of AML Reveals Progenitor-like Cells that Correlate with Prognosis. Cell 162, 184–197.

Mantovani, A., and Locati, M. (2013). Tumor-associated macrophages as a paradigm of macrophage plasticity, diversity, and polarization: lessons and open questions. Arteriosclerosis, thrombosis, and vascular biology 33, 1478–1483.

Marrack, P., Mitchell, T., Hildeman, D., Kedl, R., Teague, T.K., Bender, J., Rees, W., Schaefer, B.C., and Kappler, J. (2000). Genomic-scale analysis of gene expression in resting and activated T cells. Curr Opin Immunol 12, 206–209.

Martinez, F.O., and Gordon, S. (2014). The M1 and M2 paradigm of macrophage activation: time for reassessment. F1000prime reports 6, 13.

Mills, C.D., Kincaid, K., Alt, J.M., Heilman, M.J., and Hill, A.M. (2000). M-1/M-2 macrophages and the Th1/Th2 paradigm. Journal of immunology 164, 6166–6173.

Muller, S., Kohanbash, G., Liu, S.J., Alvarado, B., Carrera, D., Bhaduri, A., Watchmaker, P.B., Yagnik, G., Di Lullo, E., Malatesta, M., et al. (2017). Single-cell profiling of human gliomas reveals macrophage ontogeny as a basis for regional differences in macrophage activation in the tumor microenvironment. Genome biology 18, 234.

Novershtern, N., Subramanian, A., Lawton, L.N., Mak, R.H., Haining, W.N., McConkey, M.E., Habib, N., Yosef, N., Chang, C.Y., Shay, T., et al. (2011). Densely interconnected transcriptional circuits control cell states in human hematopoiesis. Cell 144, 296–309.

Pauken, K.E., and Wherry, E.J. (2015). Overcoming T cell exhaustion in infection and cancer. Trends Immunol 36, 265–276.

Perdiguero, E.G., and Geissmann, F. (2016). The development and maintenance of resident macrophages. Nat Immunol 17, 2–8.

Philip, M., Fairchild, L., Sun, L., Horste, E.L., Camara, S., Shakiba, M., Scott, A.C., Viale, A., Lauer, P., Merghoub, T., et al. (2017). Chromatin states define tumour-specific T cell dysfunction and reprogramming. Nature 545, 452–456.

Prabhakaran, S., Azizi, E., Carr, A., and Pe’er, D. (2016). Dirichlet Process Mixture Model for Correcting Technical Variation in Single-Cell Gene Expression Data. In Proceedings of The 33rd International Conference on Machine Learning, B. Maria Florina, and Q.W. Kilian, eds. (Proceedings of Machine Learning Research: PMLR), pp. 1070–1079.

Rooney, M.S., Shukla, S.A., Wu, C.J., Getz, G., and Hacohen, N. (2015). Molecular and genetic properties of tumors associated with local immune cytolytic activity. Cell 160, 48–61.

Senbabaoglu, Y., Gejman, R.S., Winer, A.G., Liu, M., Van Allen, E.M., de Velasco, G., Miao, D., Ostrovnaya, I., Drill, E., Luna, A., et al. (2016). Tumor immune microenvironment characterization in clear cell renal cell carcinoma identifies prognostic and immunotherapeutically relevant messenger RNA signatures. Genome biology 17, 231.

Singer, M., Wang, C., Cong, L., Marjanovic, N.D., Kowalczyk, M.S., Zhang, H., Nyman, J., Sakuishi, K., Kurtulus, S., Gennert, D., et al. (2017). A Distinct Gene Module for Dysfunction Uncoupled from Activation in Tumor-Infiltrating T Cells. Cell 171, 1221–1223.

Sun, R., Hu, Z., Sottoriva, A., Graham, T.A., Harpak, A., Ma, Z., Fischer, J.M., Shibata, D., and Curtis, C. (2017). Between-region genetic divergence reflects the mode and tempo of tumor evolution. Nature genetics 49, 1015–1024.

Tanaka, A., and Sakaguchi, S. (2017). Regulatory T cells in cancer immunotherapy. Cell research 27, 109–118.

Tirosh, I., Izar, B., Prakadan, S.M., Wadsworth, M.H., 2nd, Treacy, D., Trombetta, J.J., Rotem, A., Rodman, C., Lian, C., Murphy, G., et al. (2016). Dissecting the multicellular ecosystem of metastatic melanoma by single-cell RNA-seq. Science 352, 189–196.

Topalian, S.L., Drake, C.G., and Pardoll, D.M. (2015). Immune checkpoint blockade: a common denominator approach to cancer therapy. Cancer cell 27, 450–461.

van der Maaten, L., and Hinton, G. (2008). Visualizing Data using t-SNE. J Mach Learn Res 9, 2579–2605.

Verdegaal, E.M., de Miranda, N.F., Visser, M., Harryvan, T., van Buuren, M.M., Andersen, R.S., Hadrup, S.R., van der Minne, C.E., Schotte, R., Spits, H., et al. (2016). Neoantigen landscape dynamics during human melanoma-T cell interactions. Nature 536, 91–95.

Yates, L.R., Gerstung, M., Knappskog, S., Desmedt, C., Gundem, G., Van Loo, P., Aas, T., Alexandrov, L.B., Larsimont, D., Davies, H., et al. (2015). Subclonal diversification of primary breast cancer revealed by multiregion sequencing. Nature medicine 21, 751–759.

Zheng, C., Zheng, L., Yoo, J.K., Guo, H., Zhang, Y., Guo, X., Kang, B., Hu, R., Huang, J.Y., Zhang, Q., et al. (2017). Landscape of Infiltrating T Cells in Liver Cancer Revealed by Single-Cell Sequencing. Cell 169, 1342–1356 e1316.

Zilionis, R., Nainys, J., Veres, A., Savova, V., Zemmour, D., Klein, A.M., and Mazutis, L. (2017). Single-cell barcoding and sequencing using droplet microfluidics. Nat Protoc 12, 44–73.

## References

Adam Best, J., Blair, D.A., Knell, J., Yang, E., Mayya, V., Doedens, A., Dustin, M.L., Goldrath, A.W., and The Immunological Genome Project Consortium (2013). Transcriptional insights into the CD8+ T cell response to infection and memory T cell formation. Nat. Immunol. 14, 404.

Aho, A.V., and Ullman, J.D. (1983). Data structures and algorithms.

Akondy, R.S., Fitch, M., Edupuganti, S., Yang, S., Kissick, H.T., Li, K.W., Youngblood, B.A., Abdelsamed, H.A., McGuire, D.J., Cohen, K.W., et al. (2017). Origin and differentiation of human memory CD8 T cells after vaccination. Nature 552, 362–367.

Andrews, S. (2010). FastQC: a quality control tool for high throughput sequence data.

Beale, E.G., Harvey, B.J., and Forest, C. (2007). PCK1 and PCK2 as candidate diabetes and obesity genes. Cell Biochem. Biophys. 48, 89–95.

Benita, Y., Kikuchi, H., Smith, A.D., Zhang, M.Q., Chung, D.C., and Xavier, R.J. (2009). An integrative genomics approach identifies Hypoxia Inducible Factor-1 (HIF-1)-target genes that form the core response to hypoxia. Nucleic Acids Res. 37, 4587–4602.

Bhattacharyya, A. (1990). On a Geometrical Representation of Probability Distributions and its use in Statistical Inference. Calcutta Statist. Assoc. Bull. 40, 23–49.

Biswas, S.K., and Mantovani, A. (2010). Macrophage plasticity and interaction with lymphocyte subsets: cancer as a paradigm. Nat. Immunol. 11, ni.1937.

Blackinton, J.G., and Keene, J.D. (2016). Functional coordination and HuR-mediated regulation of mRNA stability during T cell activation. Nucleic Acids Res. 44, 426–436.

Bray, N.L., Pimentel, H., Melsted, P., and Pachter, L. (2016). Near-optimal probabilistic RNA-seq quantification. Nat. Biotechnol. 34, 525–527.

Bronte, V., Brandau, S., Chen, S.-H., Colombo, M.P., Frey, A.B., Greten, T.F., Mandruzzato, S., Murray, P.J., Ochoa, A., Ostrand-Rosenberg, S., et al. (2016). Recommendations for myeloid-derived suppressor cell nomenclature and characterization standards. Nat. Commun. 7, 12150.

Caton, P.W., Nayuni, N.K., Kieswich, J., Khan, N.Q., Yaqoob, M.M., and Corder, R. (2010). Metformin suppresses hepatic gluconeogenesis through induction of SIRT1 and GCN5. J. Endocrinol. 205, 97–106.

Cheadle, C. (2005). Stability Regulation of mRNA and the Control of Gene Expression. Ann. N. Y. Acad. Sci. 1058, 196–204.

Chtanova, T., Newton, R., Liu, S.M., Weininger, L., Young, T.R., Silva, D.G., Bertoni, F., Rinaldi, A., Chappaz, S., Sallusto, F., et al. (2005). Identification of T Cell-Restricted Genes, and Signatures for Different T Cell Responses, Using a Comprehensive Collection of Microarray Datasets. The Journal of Immunology 175, 7837–7847.

Coifman, R., Coppi, A., Hirn, M., and Warner, F. (2010). Diffusion Geometry Based Nonlinear Methods for Hyperspectral Change Detection.

Coifman, R.R., Lafon, S., Lee, A.B., Maggioni, M., Nadler, B., Warner, F., and Zucker, S.W. (2005). Geometric diffusions as a tool for harmonic analysis and structure definition of data: diffusion maps. Proc. Natl. Acad. Sci. U. S. A. 102, 7426–7431.

De Simone, M., De Simone, M., Arrigoni, A., Rossetti, G., Gruarin, P., Ranzani, V., Politano, C., Bonnal, R.J.P., Provasi, E., Sarnicola, M.L., et al. (2016). Transcriptional Landscape of Human Tissue Lymphocytes Unveils Uniqueness of Tumor-Infiltrating T Regulatory Cells. Immunity 45, 1135–1147.

Díaz-García, J.A., Jáimez, R.G., and Mardia, K.V. (1997). Wishart and Pseudo-Wishart Distributions and Some Applications to Shape Theory. J. Multivar. Anal. 63, 73–87.

Dobin, A., Davis, C.A., Schlesinger, F., Drenkow, J., Zaleski, C., Jha, S., Batut, P., Chaisson, M., and Gingeras, T.R. (2013). STAR: ultrafast universal RNA-seq aligner. Bioinformatics 29, 15–21.

Dushyanthen, S., Beavis, P.A., Savas, P., Teo, Z.L., Zhou, C., Mansour, M., Darcy, P.K., and Loi, S. (2015). Relevance of tumor-infiltrating lymphocytes in breast cancer. BMC Med. 13.

Faircloth, B.C., and Glenn, T.C. (2012). Not all sequence tags are created equal: designing and validating sequence identification tags robust to indels. PLoS One 7, e42543.

Farmer, S.R. (2006). Transcriptional control of adipocyte formation. Cell Metab. 4, 263–273.

Finck, R., Simonds, E.F., Jager, A., Krishnaswamy, S., Sachs, K., Fantl, W., Pe’er, D., Nolan, G.P., and Bendall, S.C. (2013). Normalization of mass cytometry data with bead standards. Cytometry A 83, 483–494.

Funes, J.M., Quintero, M., Henderson, S., Martinez, D., Qureshi, U., Westwood, C., Clements, M.O., Bourboulia, D., Pedley, R.B., Moncada, S., et al. (2007). Transformation of human mesenchymal stem cells increases their dependency on oxidative phosphorylation for energy production. Proc. Natl. Acad. Sci. U. S. A. 104, 6223–6228.

Gabrilovich, D.I. (2017). Myeloid-Derived Suppressor Cells. Cancer Immunol Res 5, 3–8.

Gesta, S., Tseng, Y.-H., and Ronald Kahn, C. (2007). Developmental Origin of Fat: Tracking Obesity to Its Source. Cell 131, 242–256.

Glimcher, L.H., Townsend, M.J., Sullivan, B.M., and Lord, G.M. (2004). Recent developments in the transcriptional regulation of cytolytic effector cells. Nat. Rev. Immunol. 4, 900–911.

Görür, D., and Rasmussen, C.E. (2010). Dirichlet Process Gaussian Mixture Models: Choice of the Base Distribution. J. Comput. Sci. Technol. 25, 653–664.

Grivennikov, S.I., Greten, F.R., and Karin, M. (2010). Immunity, inflammation, and cancer. Cell 140, 883–899.

Halko, N., Martinsson, P.-G., and Tropp, J.A. (2009). Finding structure with randomness: Probabilistic algorithms for constructing approximate matrix decompositions.

Hartigan, J.A., and Hartigan, P.M. (1985). The Dip Test of Unimodality. Ann. Stat. 13, 70–84.

Ho, P.-C., Bihuniak, J.D., Macintyre, A.N., Staron, M., Liu, X., Amezquita, R., Tsui, Y.-C., Cui, G., Micevic, G., Perales, J.C., et al. (2015). Phosphoenolpyruvate Is a Metabolic Checkpoint of Anti-tumor T Cell Responses. Cell 162, 1217–1228.

Jahrsdörfer, B., Vollmer, A., Blackwell, S.E., Maier, J., Sontheimer, K., Beyer, T., Mandel, B., Lunov, O., Tron, K., Nienhaus, G.U., et al. (2010). Granzyme B produced by human plasmacytoid dendritic cells suppresses T-cell expansion. Blood 115, 1156–1165.

Jaitin, D.A., Kenigsberg, E., Keren-Shaul, H., Elefant, N., Paul, F., Zaretsky, I., Mildner, A., Cohen, N., Jung, S., Tanay, A., et al. (2014). Massively parallel single-cell RNA-seq for marker-free decomposition of tissues into cell types. Science 343, 776–779.

Jebara, T., Kondor, R., & Howard, A. (2004). Probability product kernels. J. Mach. Learn. Res. 5, 819–844.

Jeffrey, K.L., Brummer, T., Rolph, M.S., Liu, S.M., Callejas, N.A., Grumont, R.J., Gillieron, C., Mackay, F., Grey, S., Camps, M., et al. (2006). Positive regulation of immune cell function and inflammatory responses by phosphatase PAC-1. Nat. Immunol. 7, 274–283.

Kim, D., Pertea, G., Trapnell, C., Pimentel, H., Kelley, R., and Salzberg, S.L. (2013). TopHat2: accurate alignment of transcriptomes in the presence of insertions, deletions and gene fusions. Genome Biol. 14, R36.

Klein, A.M., Mazutis, L., Akartuna, I., Tallapragada, N., Veres, A., Li, V., Peshkin, L., Weitz, D.A., and Kirschner, M.W. (2015). Droplet barcoding for single-cell transcriptomics applied to embryonic stem cells. Cell 161, 1187–1201.

Kumar, B.V., Connors, T.J., and Farber, D.L. (2018). Human T Cell Development, Localization, and Function throughout Life. Immunity 48, 202–213.

Lefterova, M.I., and Lazar, M.A. (2009). New developments in adipogenesis. Trends Endocrinol. Metab. 20, 107–114.

Levine, J.H., Simonds, E.F., Bendall, S.C., Davis, K.L., Amir, E.-A.D., Tadmor, M.D., Litvin, O., Fienberg, H.G., Jager, A., Zunder, E.R., et al. (2015). Data-Driven Phenotypic Dissection of AML Reveals Progenitor-like Cells that Correlate with Prognosis. Cell 162, 184–197.

Li, B., and Dewey, C.N. (2011). RSEM: accurate transcript quantification from RNA-Seq data with or without a reference genome. BMC Bioinformatics 12, 323.

Maaten, L. van der, and Hinton, G. (2008). Visualizing Data using t-SNE. J. Mach. Learn. Res. 9, 2579–2605.

Macosko, E.Z., Basu, A., Satija, R., Nemesh, J., Shekhar, K., Goldman, M., Tirosh, I., Bialas, A.R., Kamitaki, N., Martersteck, E.M., et al. (2015). Highly Parallel Genome-wide Expression Profiling of Individual Cells Using Nanoliter Droplets. Cell 161, 1202–1214.

Makino, Y., Nakamura, H., Ikeda, E., Ohnuma, K., Yamauchi, K., Yabe, Y., Poellinger, L., Okada, Y., Morimoto, C., and Tanaka, H. (2003). Hypoxia-inducible factor regulates survival of antigen receptor-driven T cells. J. Immunol. 171, 6534–6540.

Mamedov, T.G., Pienaar, E., Whitney, S.E., TerMaat, J.R., Carvill, G., Goliath, R., Subramanian, A., and Viljoen, H.J. (2008). A fundamental study of the PCR amplification of GC-rich DNA templates. Comput. Biol. Chem. 32, 452–457.

Manley, L.J., Ma, D., and Levine, S.S. (2016). Monitoring Error Rates In Illumina Sequencing. J. Biomol. Tech. 27, 125–128.

Mantovani, A., and Locati, M. (2013). Tumor-associated macrophages as a paradigm of macrophage plasticity, diversity, and polarization: lessons and open questions. Arterioscler. Thromb. Vasc. Biol. 33, 1478–1483.

Mantovani, A., Allavena, P., Sica, A., and Balkwill, F. (2008). Cancer-related inflammation. Nature 454, 436–444.

Mills, C.D., Kincaid, K., Alt, J.M., Heilman, M.J., and Hill, A.M. (2000). M-1/M-2 macrophages and the Th1/Th2 paradigm. J. Immunol. 164, 6166–6173.

Moreno-Sánchez, R., Rodríguez-Enríquez, S., Saavedra, E., Marín-Hernández, A., and Gallardo-Pérez, J.C. (2009). The bioenergetics of cancer: Is glycolysis the main ATP supplier in all tumor cells? Biofactors 35, 209–225.

Mues, C., Zhou, J., Manolopoulos, K.N., Korsten, P., Schmoll, D., Klotz, L.-O., Bornstein, S.R., Klein, H.H., and Barthel, A. (2009). Regulation of glucose-6-phosphatase gene expression by insulin and metformin. Horm. Metab. Res. 41, 730–735.

Murphy., K.P. (2007). Conjugate bayesian analysis of the gaussian distribution. Tech. Rep.

Patro, R., Duggal, G., Love, M.I., Irizarry, R.A., and Kingsford, C. (2017). Salmon provides fast and bias-aware quantification of transcript expression. Nat. Methods 14, 417–419.

Perera, R.J., Marcusson, E.G., Koo, S., Kang, X., Kim, Y., White, N., and Dean, N.M. (2006). Identification of novel PPARγ target genes in primary human adipocytes. Gene 369, 90–99.

Pitman, J. (2006). Combinatorial Stochastic Processes (Springer Science & Business Media).

Platanias, L.C. (2005). Mechanisms of type-I- and type-II-interferon-mediated signalling. Nat. Rev. Immunol. 5, 375–386.

Plitas, G., Konopacki, C., Wu, K., Bos, P.D., Morrow, M., Putintseva, E.V., Chudakov, D.M., and Rudensky, A.Y. (2016). Regulatory T Cells Exhibit Distinct Features in Human Breast Cancer. Immunity 45, 1122–1134.

Prabhakaran, S., Azizi, E., Carr, A., & Pe’er, D (2016). Dirichlet Process Mixture Model for Correcting Technical Variation in Single-Cell Gene Expression Data. Journal of Machine Learning Research, W&CP (ICML) 48, 1070–1079.

Sainz, B., Jr, Martín, B., Tatari, M., Heeschen, C., and Guerra, S. (2014). ISG15 is a critical microenvironmental factor for pancreatic cancer stem cells. Cancer Res. 74, 7309–7320.

Schietinger, A., Delrow, J.J., Basom, R.S., Blattman, J.N., and Greenberg, P.D. (2012). Rescued Tolerant CD8 T Cells Are Preprogrammed to Reestablish the Tolerant State. Science 335, 723–727.

Setty, M., Tadmor, M.D., Reich-Zeliger, S., Angel, O., Salame, T.M., Kathail, P., Choi, K., Bendall, S., Friedman, N., and Pe’er, D. (2016). Wishbone identifies bifurcating developmental trajectories from single-cell data. Nat. Biotechnol. 34, 637–645.

Sica, A., and Mantovani, A. (2012). Macrophage plasticity and polarization: in vivo veritas. J. Clin. Invest. 122, 787.

Singer, M., Wang, C., Cong, L., Marjanovic, N.D., Kowalczyk, M.S., Zhang, H., Nyman, J., Sakuishi, K., Kurtulus, S., Gennert, D., et al. (2016). A Distinct Gene Module for Dysfunction Uncoupled from Activation in Tumor-Infiltrating T Cells. Cell 166, 1500–1511.e9.

Smith-Garvin, J.E., Koretzky, G.A., and Jordan, M.S. (2009). T Cell Activation. Annu. Rev. Immunol. 27, 591–619.

Subramanian, A., Tamayo, P., Mootha, V.K., Mukherjee, S., Ebert, B.L., Gillette, M.A., Paulovich, A., Pomeroy, S.L., Golub, T.R., Lander, E.S., et al. (2005). Gene set enrichment analysis: a knowledge-based approach for interpreting genome-wide expression profiles. Proc. Natl. Acad. Sci. U. S. A. 102, 15545–15550.

Swiecki, M., and Colonna, M. (2015). The multifaceted biology of plasmacytoid dendritic cells. Nat. Rev. Immunol. 15, 471–485.

Tao, T., and Vu, V. (2005). On random ±1 matrices: Singularity and determinant. Random Struct. Algorithms 28, 1–23.

Ugel, S., De Sanctis, F., Mandruzzato, S., and Bronte, V. (2015). Tumor-induced myeloid deviation: when myeloid-derived suppressor cells meet tumor-associated macrophages. J. Clin. Invest. 125, 3365–3376.

Valle, S., Li, W., and Joe Qin, S. (1999). Selection of the Number of Principal Components: The Variance of the Reconstruction Error Criterion with a Comparison to Other Methods†. Ind. Eng. Chem. Res. 38, 4389–4401.

Vallejos, C.A., Risso, D., Scialdone, A., Dudoit, S., and Marioni, J.C. (2017). Normalizing single-cell RNA sequencing data: challenges and opportunities. Nat. Methods 14, 565–571.

Villani, A.-C., Satija, R., Reynolds, G., Sarkizova, S., Shekhar, K., Fletcher, J., Griesbeck, M., Butler, A., Zheng, S., Lazo, S., et al. (2017). Single-cell RNA-seq reveals new types of human blood dendritic cells, monocytes, and progenitors. Science 356.

Wherry, J.E. (2011). T cell exhaustion. Nat. Immunol. 12, ni.2035.

Wherry, J.E., and Kurachi, M. (2015). Molecular and cellular insights into T cell exhaustion. Nat. Rev. Immunol. 15, nri3862.

Whitfield, M.L., Sherlock, G., Saldanha, A.J., Murray, J.I., Ball, C.A., Alexander, K.E., Matese, J.C., Perou, C.M., Hurt, M.M., Brown, P.O., et al. (2002). Identification of Genes Periodically Expressed in the Human Cell Cycle and Their Expression in Tumors. Mol. Biol. Cell 13, 1977–2000.

Youngblood, B., Hale, J.S., Kissick, H.T., Ahn, E., Xu, X., Wieland, A., Araki, K., West, E.E., Ghoneim, H.E., Fan, Y., et al. (2017). Effector CD8 T cells dedifferentiate into long-lived memory cells. Nature 552, 404–409.

Zheng, G.X.Y., Terry, J.M., Belgrader, P., Ryvkin, P., Bent, Z.W., Wilson, R., Ziraldo, S.B., Wheeler, T.D., McDermott, G.P., Zhu, J., et al. (2017). Massively parallel digital transcriptional profiling of single cells. Nat. Commun. 8, 14049.

Zilionis, R., Nainys, J., Veres, A., Savova, V., Zemmour, D., Klein, A.M., and Mazutis, L. (2016). Single-cell barcoding and sequencing using droplet microfluidics. Nat. Protoc. 12, nprot.2016.154.

Zunder, E.R., Finck, R., Behbehani, G.K., Amir, E.-A.D., Krishnaswamy, S., Gonzalez, V.D., Lorang, C.G., Bjornson, Z., Spitzer, M.H., Bodenmiller, B., et al. (2015). Palladium-based mass tag cell barcoding with a doublet-filtering scheme and single-cell deconvolution algorithm. Nat. Protoc. 10, 316–333.

